# Inferring domestic goat demographic history through ancient genome imputation

**DOI:** 10.1101/2025.04.18.649576

**Authors:** Jolijn A. M. Erven, Alice Etourneau, Marjan Mashkour, Mahesh Neupane, Phillipe Bardou, Alessandra Stella, Andrea Talenti, Clet Wandui Masiga, Curt Van Tassell, Emily Clark, François Pompanon, Licia Colli, Marcel Amillis, Marco Milanesi, Paola Crepaldi, The VarGoats Consortium, Bertrand Servin, Ben Rosen, Gwenola Tosser-Klopp, Kevin G. Daly

**Affiliations:** UCD School of Agriculture and Food Science, University College Dublin, Belfield, Ireland; UCD Conway Institute of Biomolecular and Biomedical Research, University College Dublin, Belfield, Ireland; GenPhySE, Université de Toulouse, INRAE, ENVT, F-31326, Castanet Tolosan, France; Archéozoologie et Archéobotanique: Sociétés, Pratiques et Environnements UMR 7209 du Centre national de la recherche scientifique (CNRS) et Muséum national d’Histoire naturelle (MNHN), Paris, France; Bioarchaeology Laboratory, Central Laboratory, Archaeozoology section, University of Tehran, Tehran, Iran; Animal Genomics and Improvement Laboratory, Agricultural Research Service, United States Department of Agriculture, Beltsville, Maryland, USA; Sigenae, GenPhySE, Université de Toulouse, INRAE, ENVT, F-31326, Castanet Tolosan, France; National Research Council of Italy, CNR, Milan, Italy; The Roslin Institute, Royal (Dick) School of Veterinary Studies, University of Edinburgh, Midlothian, EH25 9RG, United Kingdom; Tropical Institute of Development Innovations (TRIDI), Uganda; EMBL’s European Bioinformatics Institute, Wellcome Genome Campus, CB10 1SD, United Kingdom; Université Grenoble Alpes, Université Savoie Mont Blanc, CNRS, LECA, Grenoble, France; Dipartimento di Scienze Animali, della Nutrizione e degli Alimenti (DIANA), Università Cattolica del Sacro Cuore, via Emilia Parmense 84, 29122 Piacenza (PC), Italy; BioDNA Centro di ricerca sulla Biodiversità e sul DNA Antico, Università Cattolica del Sacro Cuore, Facoltà di Scienze Agrarie, Alimentari e Ambientali, via Emilia Parmense 84, 29122 Piacenza (PC), Italy; Centre for Research in Agricultural Genomics (CRAG), CSIC-IRTA-UAB-UB, Campus Universitat Autònoma de Barcelona, Bellaterra, Spain; Department for Innovation in Biological, Agro-Food and Forest Systems, University of Tuscia, Viterbo, Italy; Dipartimento di Scienze Agrarie e Ambientali, Produzione, Territorio, Agroenergia, Università di Milano, Italy

**Author notes:** Author for Correspondence: Kevin G. Daly, School of Agriculture and Food Science, University College Dublin, Belfield, Ireland. These authors contributed equally (Ben Rosen, Gwenola Tosser-Klopp, Kevin G. Daly).

**Keywords:** Imputation, aDNA, Goat domestication

## Abstract

Goats were among the earliest managed animals, making them a natural model to explore the genetic consequences of domestication. However, a challenge in ancient genomic analysis is the relatively low genome coverage for most samples, limiting analysis to pseudohaploid genotypes. Genotype imputation offers potential to alleviate this limitation by improving information content and accuracy in low coverage genomes. To test this we used published high coverage (>8✕) goat palaeogenomes, imputing downsampled genomes using the VarGoats dataset (1,372 individuals) as a reference panel. Measuring concordance between imputed and high coverage genotypes, we find high concordance after filtering for common (>5%), high confidence variants, with 0.5✕ genomes reaching >0.97 concordance. There is a trade-off between coverage, genotype probability (GP) thresholds, and genotype recovery, where higher coverage and more lenient GP thresholds result in higher recovery, and a reduction in heterozygous false-positive rates with stricter thresholds. We then imputed 36 goat palaeogenomes with ≥0.5✕ coverage to examine runs-of-homozygosity (ROH) and identity-by-descent (IBD) patterns. Using a novel approach combining ROH profiles across tools, we find that among Neolithic goats, ROH increases with distance from the Zagros Mountains, suggesting a large effect of the initial dispersal of managed herds. Inbreeding levels decrease across Southwest Asia in more recent periods. IBD mirrored this pattern, with less relatedness in the early herding site of Ganj Dareh compared to higher relatedness in goats from later in the dispersal process. These findings provide insights into the genetic consequences of early goat management on demography, and confirm the utility of imputation in leveraging low coverage palaeogenomes.

**Significance:** Paleogenomics offers crucial insight into how animals were domesticated, but poor DNA preservation in ancient remains often limits the reach of genetic analyses. We utilise a cutting-edge technique, genotype imputation, to recover missing genetic information from ancient low coverage goat genomes and shine light on their domestication process. Early domestic goats showed low overall runs-of-homozygosity (ROH) and relatedness. We find that during the Neolithic, runs-of-homozygosity (ROH) and relatedness among goats increased, likely a consequence of the movement of herds beyond their natural range by humans. Inbreeding levels decline in more recent periods, potentially due to expanded herd sizes, animal trade networks, or improved husbandry practices. These findings challenge long-standing assumptions about domestication bottlenecks and highlight how ancient DNA can be used to uncover complex evolutionary histories, even from low coverage ancient samples.

## Introduction

The domestic goat (*Capra hircus*) was one of the earliest animals brought under human management, likely beginning before 8,200 years BCE (Before Common Era) in Southwest Asia (Zeder 2008). Faunal assemblages from the Central Zagros Mountains of Iran show demographic indicators of young slaughtering consistent with a preference for kid- and milk-producing females (Zeder & Hesse 2000). Genomic data from these assemblages support this conclusion, showing differential uniparental diversity patterns and signs of recent inbreeding (homozygosity tracts >5Mb, tracing back to the last 4-8 generations or more recently (Stoffel et al. 2021)), expected within a more limited breeding population (Daly et al. 2021). Control of goats is posited at other regions during this Aceramic/Pre-Pottery Neolithic period. For example, at Aşıklı Höyük in Central Anatolia, there is evidence for a gradual development of goats and sheep being kept within enclosed spaces of the settlement, and with tethering of animals likely practiced (Abell et al. 2019; Zimmermann et al. 2018). Across the Southern Levant, there are indications of fodder usage at Abu Ghosh ∼8,000 BCE, and a widespread increase in the occurrence of goat remains relative to wild animals (Makarewicz & Tuross 2012; Munro et al. 2018). Goat management practices — and managed goats themselves — dispersed throughout neighbouring environs in the following millennia. Targeted slaughtering of young, grown males became ubiquitous in Southwest Asia by the mid 7th millennium BCE (Arbuckle & Atici 2013), while managed goat herds reached Northeast Iran in the mid-to-late 8th millennium BCE (Roustaei et al. 2015) and Europe by the late 7th millennium BCE (Zeder 2008; Arbuckle et al. 2014).

The consequences of these initial phases of management and diffusion have been broadly characterized using palaeogenomes, with evidence of recent inbreeding among early managed goats in the Zagros Mountains (Daly et al. 2018, 2021). However, genetic data from ancient remains is defined by its relatively poor quality, due to post-mortem degradation and associated low sequencing depth. Consequently, most palaeogenomic analyses are limited to pseudohaploid genotypes, and lack the richer demographic inferences possible from diploid genotypes. Similarly, missing genotypes restrict the accurate detection of shared haplotypes between chromosomes, both within-individual (runs-of-homozygosity, or ROH) and between-individual (identity-by-descent, or IBD) regions (Peripolli et al. 2017; Browning & Browning 2012; Blondeau Da Silva et al. 2024). This excludes many palaeogenomes from analyses such as IBD-based demographic inference (Fournier et al. 2023), or their incorporation into ancestral recombination graphs (Speidel et al. 2021).

A long-standing model of domestication is the domestication bottleneck, where a small number of animals are used to found managed herds and interbreeding with wild populations (Zeder et al. 2006). Under this standard model, the resulting managed population is expected to show reduced genetic diversity due to the imposed genetic drift and lack of gene flow (Frantz et al. 2020). A concomitant prediction is an increase of inbreeding events and ROHs, due to the reduced effective population size. Genetic data has required increasingly nuanced modifications to this domestication-diversity model. Genomes from horses (Schubert et al. 2014), dogs (Marsden et al. 2016), and chickens (Wang et al. 2021) have revealed a greater burden of deleterious variation in domestic populations compared to wild, suggestive of a fixation of variants which may otherwise be purged in a larger population.

However, in both sheep (Sandoval-Castellanos et al. 2024) and goat (Daly et al. 2021) mitochondrial diversity is high in the earliest millenia of herding, but subsequently declines. Data from sorghum, maize and barley suggest a post-domestication of erosion of diversity rather than an initial sharp decrease (Allaby et al. 2019). In contrast, ancient horse populations show several distinct population declines, linked to episodes of intensified management and breeding practices (Librado et al. 2024). Nuclear genomes from early herded goats also show a degree of inbreeding, but paradoxically higher estimated heterozygosity values than later domestics (Daly et al. 2021). These apparently contradictory findings necessitate a re-assessment of the standard domestication model (Frantz et al. 2020), informed by species-specific biological and cultural histories. A meaningful re-assessment would require ancient samples with sufficient coverage (information content) to provide the power needed to address these standard domestication models.

A powerful and cost-effective way to increase the information content of ancient samples is genotype imputation. Genotype imputation is the statistical inference of genotypes at unknown or missing sites, using known haplotype variation in a set of reference individuals (Browning & Browning 2016). This method facilitates the inference of diploid genotypes from low coverage ancient genomes, enabling haplotype-aware and genealogical analyses. Genotype imputation is widely employed in ancient humans (Gamba et al. 2014; Martiniano et al. 2017; Cassidy et al. 2020; Hui et al. 2020; Ariano et al. 2022; Ausmees et al. 2022; Ausmees & Nettelblad 2023; Sousa da Mota et al. 2023), ancient pigs (Erven et al. 2022), ancient bovids (Erven et al. 2024; Rossi et al. 2024), ancient canids (Bougiouri et al. 2024), historic-era horses (Todd et al. 2023), and modern livestock (Wang et al. 2022; Ding et al. 2023; Lloret-Villas et al. 2023; Zhang et al. 2023). The method has proven to be a step-change in our increasingly fine-scale understanding of human population history, offering insights into prehistoric social structures at the level of shared haplotypes (Cassidy et al. 2020; Allentoft et al. 2024; Lazaridis et al. 2025), demographic bottlenecks in the establishment of insular populations (Ariano et al. 2022), and the trajectory of selection within different ancient ancestries (Vaughn & Nielsen 2024; Irving-Pease et al. 2024).

The VarGoats project has surveyed commercial and local breeds, the progenitor species *Capra aegagrus* (the bezoar ibex), and other *Capra* species to synthesize a dataset of over 1,100 novel or publicly available genomes (Denoyelle et al. 2021), now expanded to 1,372 individual genomes representing 155 country-breed groupings with a worldwide distribution. This global-scale genomic dataset represents an opportunity to employ imputation on goat palaeogenomes. With the aim to test the long-standing assumption of an early domestication-related population bottleneck in livestock species, we first performed downsampling experiments on high quality goat palaeogenomes to ensure imputation accuracy, and then utilized the imputed genotypes to directly assess changing inbreeding patterns through time.

## Results

We systematically evaluated genotype imputation of ancient low coverage goat genomes, following the GLIMPSE2 pipeline (Rubinacci et al. 2023), schematically described in Figure 1. The accuracy of imputation – the frequency at which imputed genotypes matched non-imputed genotypes for the same sample – was tested by downsampling four ancient high coverage goat genomes. These four genomes comprise different ancestries across domestic and wild goats which are variably represented within our reference panel (Figure 1, Table S1-2). Heterozygous and homozygous alternative genotypes are expected to impute less effectively than homozygous reference (Zhang et al. 2021), so we particularly focused on these former variants in the assessment of accuracy. We additionally assessed the number of variant sites recovered per-sample in the imputed data, relative to the number of variant sites covered by the sequencing data alone.

**Figure 1:**
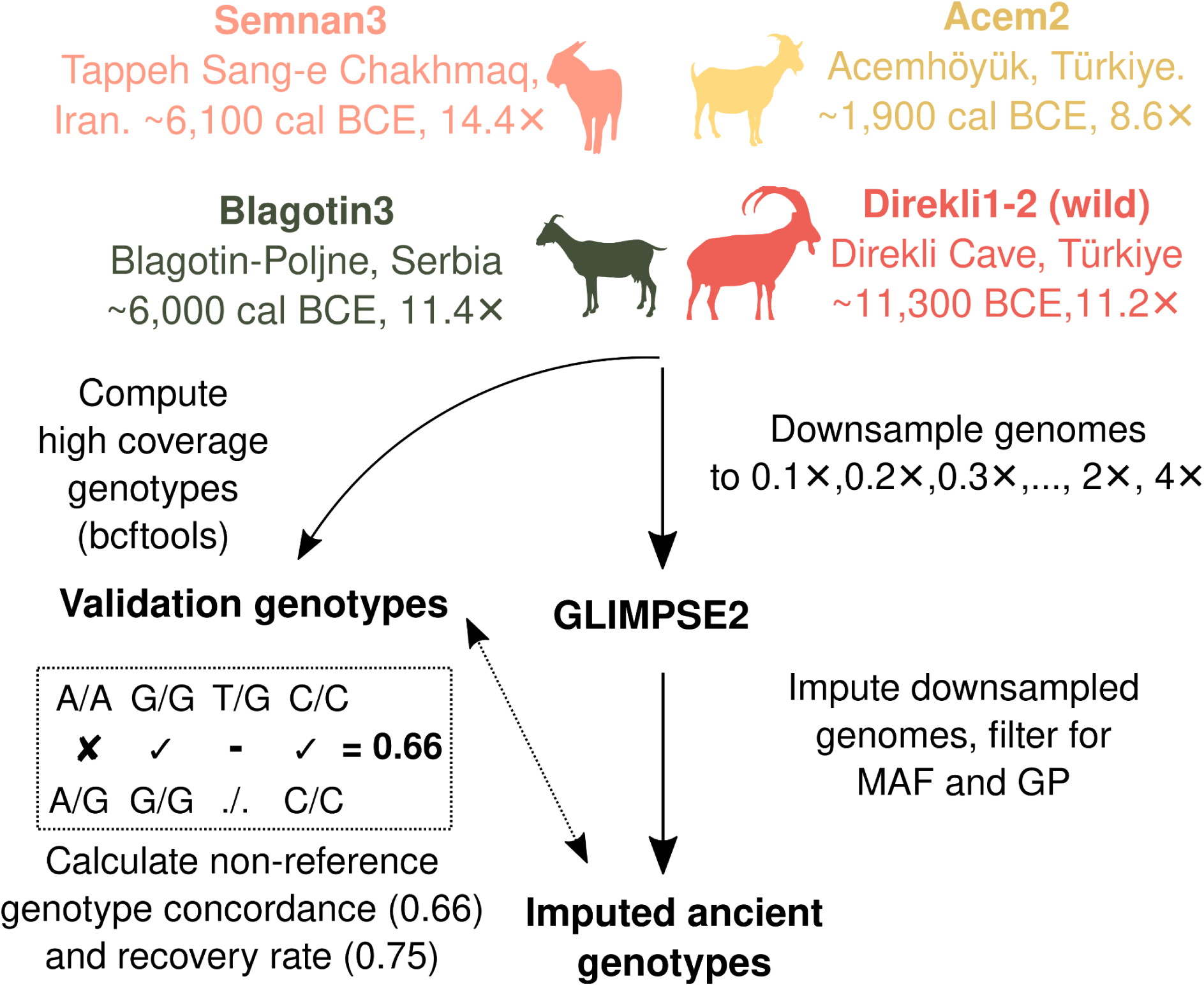
Overview of high coverage goat palaeogenomes and downsampling approach. Goat silhouettes from phylopic.org.

### Imputation accuracy

The imputation accuracy of ancient goats, measured by the genotype concordance of non-reference genotypes between high coverage and imputed genotypes (heterozygous and homozygous alternative) showed a clear dosage dependency (Figure 2, Table S5-6). Consistent with previous findings in ancient humans (Sousa da Mota et al. 2023), cattle (Erven et al. 2024) and canids (Bougiouri et al. 2024), higher coverage led to greater imputation concordance. This effect of coverage is more pronounced in rare alleles (<2% MAF), which show a greater decline in genotype concordance as coverage decreases compared to more common alleles (Figure 2). Genotype concordance plateaus between the 2-5% and 5-10% MAF tranches (Figure 2, S1-2), where it stabilises from 0.5✕ coverage onwards (>0.90 concordance; Figures S1-2, Table S5). We find concordance gains at higher coverage in both heterozygous and homozygous alternative genotypes, with sample-specific effects likely due to uneven representation of ancient ancestries in the reference panel (Supplementary Note 1, Figures S3-4, Tables S7-8).

**Figure 2:**
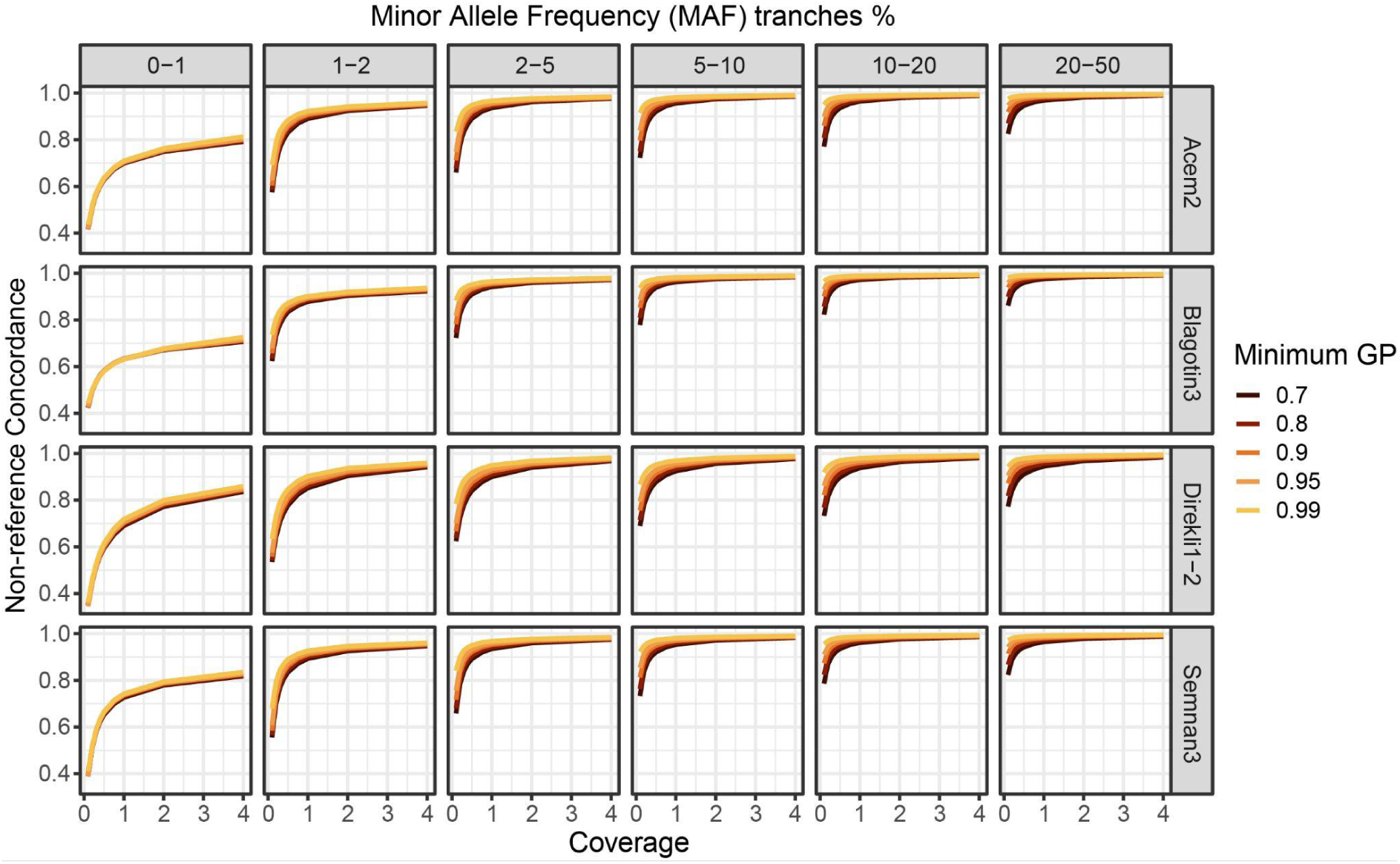
Non-reference concordance across MAF tranches for four samples, with downsampled coverages ranging from 0.1-4✕. Concordance values are calculated for transitions and transversions combined. Colors indicate different genotype probability (GP) thresholds. 0.1-1✕ is shown in Figure S1.

Genotype concordance was additionally sensitive to the minimum genotype probability (GP) threshold, which was applied to the imputed data as a quality filter (Methods – Validation of genotype imputation). A more stringent GP threshold resulted in higher genotype concordance; this gain in genotype concordance was greater in the 0.1✕ imputed data compared to the higher coverage imputed data (>1✕; Figure 2, S1), as observed in humans and cattle (e.g. (Erven et al. 2024; Sousa da Mota et al. 2023)). In general, we observed genotype concordance for common alleles (MAF tranches >5%) exceeding 0.69 and 0.87 at a minimum coverage of 0.1✕ for GP70 and GP99; 0.89 and 0.95 for a minimum coverage of 0.5✕; and 0.96 and 0.98 for a minimum coverage of 2✕ (Table S5).

We observed differences in genotype concordance between transversions and transitions, where transitions showed a lower concordance than transversions (Figures 3, S2, Table S6), particularly at MAF tranches below 2%. Heterozygous genotypes specifically appear to be the primary driver of this divide between transversion and transition concordance rates (Figures S3-4, Table S6). Differences between transversions and transitions in more common alleles (MAF tranches >5%) are minimal (Figures S5-7): transversions have on average a 0.019, 0.014 and 0.011 higher genotype concordance at 0.1✕, 0.5✕ and 2✕, respectively (Table S6). Finally, we observe variation in the difference of transition/transversion concordance rates between individuals (Figure 3). This does not appear to be driven by residual post-mortem deamination error, as we see little evidence of systematic differences in error at CpG sites (Table S2), and may be instead due to a combination of sample specific-error and dissimilarity to the reference panel.

**Figure 3:**
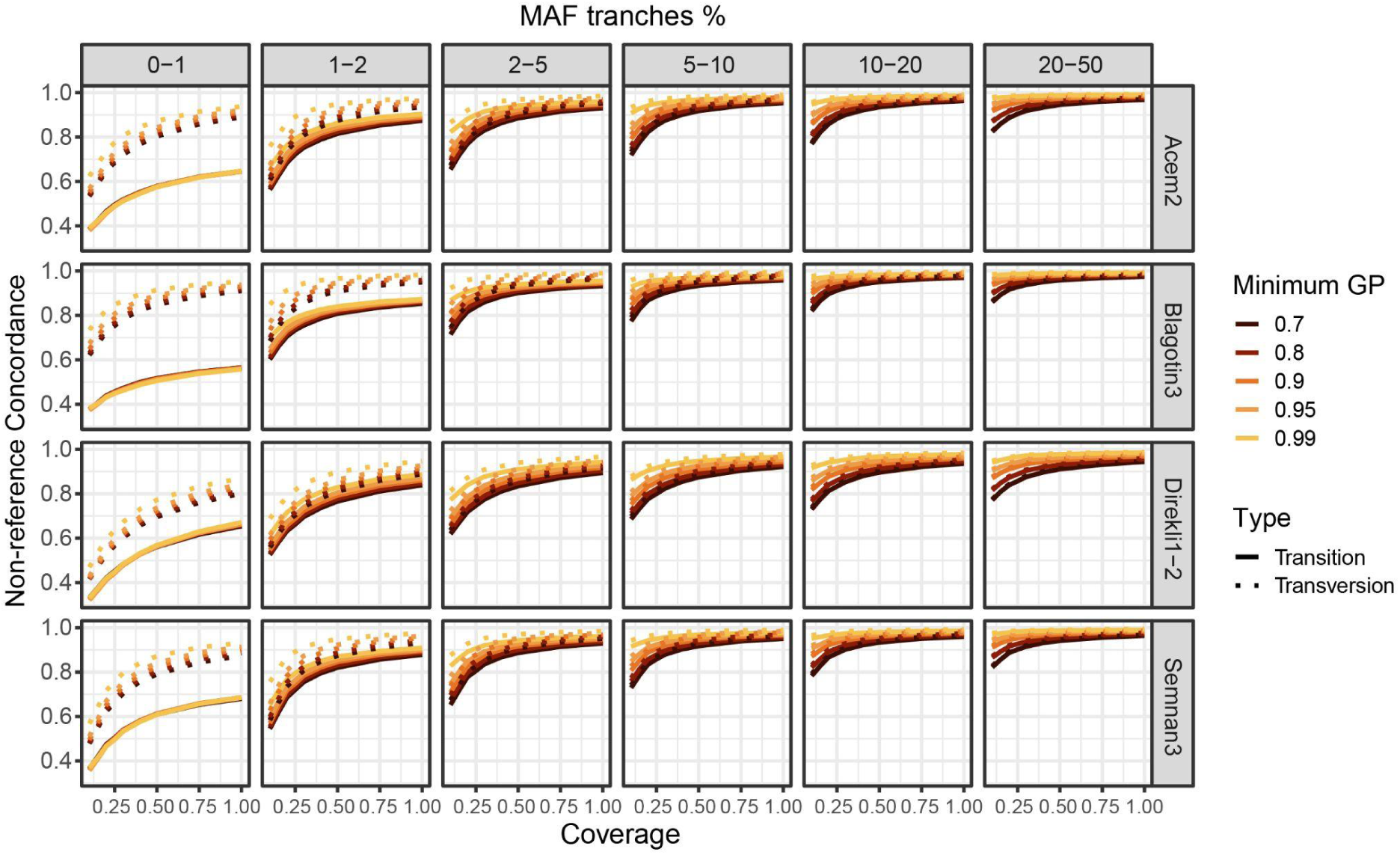
Non-reference concordance across MAF tranches for four samples, with downsampled coverages ranging from 0.1-1✕. Concordance values are calculated for transitions and transversions separately, depicted by Type. Colors indicate different genotype probability (GP) thresholds. 0.1-4✕ is shown in Figure S2.

We then examined heterozygous sites, known to be more challenging to impute (Figure S3; (Sousa da Mota et al. 2023; Erven et al. 2024; Hui et al. 2020; Escobar-Rodríguez & Veeramah 2024)). We calculated the heterozygous false-positive rate (FPR), measured as number of imputed heterozygous called as homozygous in the validation dataset (FP) divided by the sum of the number of FP and correctly-imputed homozygous sites (TN). The heterozygous FPR was sensitive to coverage and GP, with larger decreases in FPR seen at stricter GP thresholds (Figures 4, S8, Tables S7-8). We also see that variation between GP thresholds shows sample-specific patterns (Supplementary Note 2, Figure S8, Tables S7-8). We finally explored the false-negative rate (FNR) of heterozygous sites and found a large effect of the GP filter (Supplementary Note 2).

**Figure 4:**
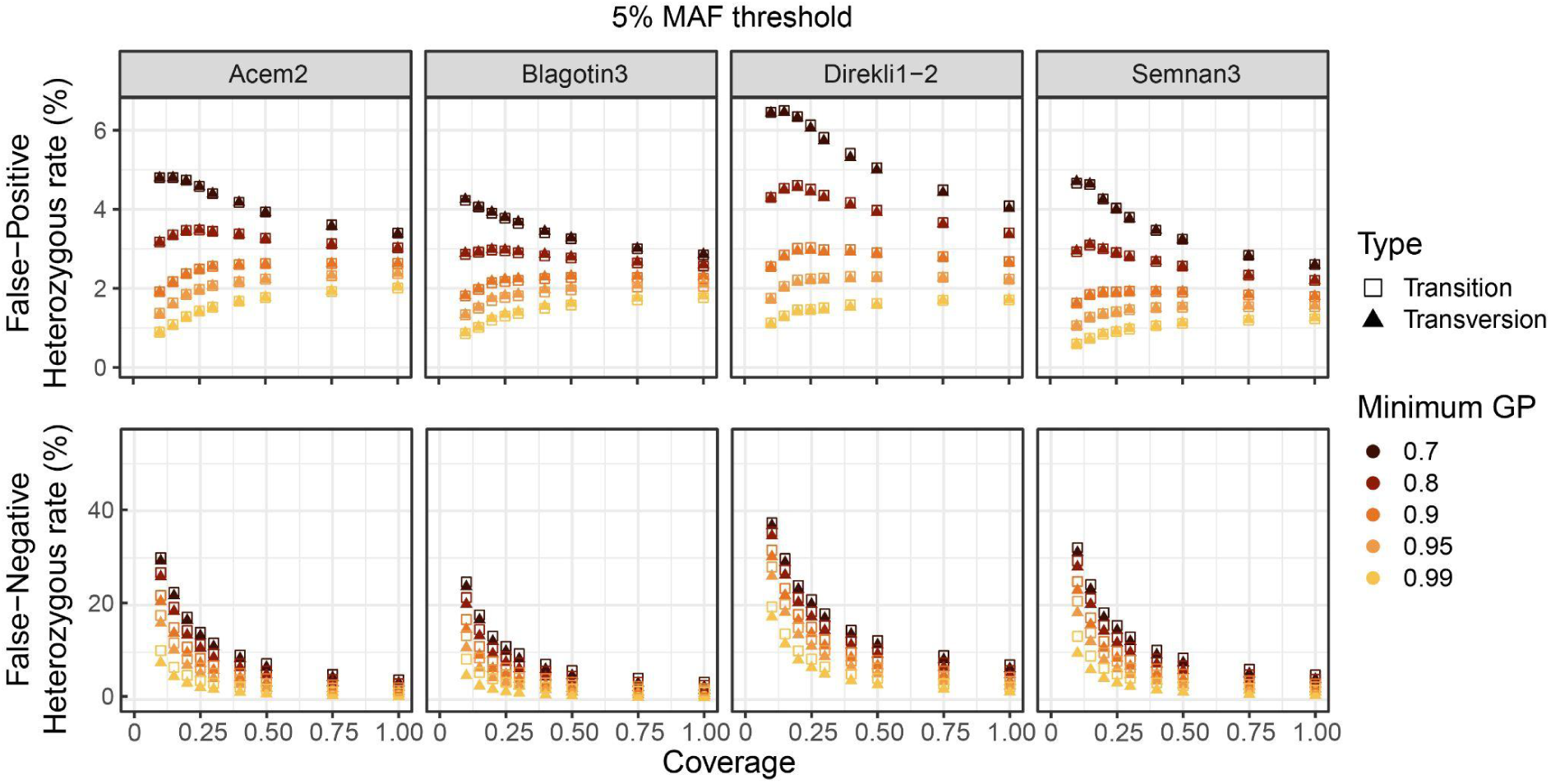
Patterns of imputed heterozygous veracity. A) false-positive rate (FPR) and B) false-negative rate (FNR), at the 5% MAF threshold. “True” heterozygotes are defined using the validation non-imputed genotypes calculated using the entirety of sequencing data available for each test sample.

Given the concordance rates observed across individuals (including the Epipalaeolithic bezoar Direkli1-2) at 0.5✕ and associated suppression of false heterozygous genotypes (FPR), we chose to focus analyses at this coverage level and primarily use a GP99 filter in the following sections.

### Imputation recovery

There is a clear positive trend between coverage, GP threshold, and site recovery; higher coverage and more lenient GP thresholds lead to a greater number of imputed sites (Figures S9-10, Table S9). The total number of non-reference sites recovered for the test samples at 0.5✕, 5% MAF, and the strictest GP (0.99) are 1.9M (Acem2), 2.1M (Blagotin3), 1.5M (Direkli1-2), and 1.9M (Semnan3) (Table S9). Lowering the GP threshold to 0.95 increases the number of recovered sites to a greater degree than subsequent lower GP thresholds (Figures S9-10). On average, the number of recovered heterozygous sites was greater than the recovered homozygous alternative sites, with low coverages (<0.4✕) an exception (Figure S10), possibly due to a higher heterozygous FPR at these coverage levels (Figure 4).

We compared the recovery rate before and after imputation by comparing pseudohaploid (i.e. the genetic data which would be considered if no imputation was performed) and imputed calls, focusing on common alleles (MAF ≥5%). Imputation improves recovery rates, particularly for low coverage samples (<1✕) (Figure S11). Moreover, imputation also enables us to call heterozygotes variants which show a low error rate (∼1.5% FPR for 0.5✕, Figure 4). Finally, we compared different GLIMPSE software to evaluate which pipeline would be most suitable for our data (Supplementary Note 3; Table S10-11). GLIMPSE1 (Rubinacci et al. 2021) showed a slightly higher genotype concordance (Figure S12), while GLIMPSE2 showed higher recovery rates (Figure S13). We opted for GLIMPSE2, due to the substantially higher recovery in imputed sites (e.g. ∼12% gain at MAF 5%, GP99 at 0.5✕).

### Genome-wide concordance and heterozygous FPR

We next explored differences in chromosomal region imputation accuracy. We calculated non-reference concordance for our 0.5✕ downsampled genomes with a 5% MAF and GP99 threshold across the genome by utilising a sliding window approach (500kb window, 100kb step). Non-reference concordance was relatively stable in all four samples, with the lowest observed for the Epipaleolithic wild bezoar Direkli1-2 (Figures S14-17). The bottom 1st percentile of windows were considered as outlier regions; the lowest concordance was observed in Semnan3 on chromosome 27 (Figure S17). The number of individual outlier regions ranged from 91 to 102, with Semnan3 exhibiting the fewest and Direkli1-2 the most. Among the identified outlier regions, 12 were detected in three or more samples (represented as black bars in Figures S14-17).

Notably, four of the imputation outlier regions are located within or overlap with the first and last 2 Mb of chromosomes, likely reflecting the inaccurate imputation of variants mapping to telomeric regions (Table S12). Additionally, three regions contain olfactory receptor or immunity-related genes (GO terms are in Table S12; pooled regions show no specific GO enrichment), consistent with previous findings of difficult to impute regions in both ancient and modern cattle (Zhang et al. 2023; Erven et al. 2024). Finally, we characterized chromosomal distributions of the heterozygous FPR and found a greater degree of sample-specific variation than compared to concordance rates (Supplementary Note 4, Figures S18-21), Table S13). The outlier regions present in the majority of the samples (at least three; Tables S12-13) for both non-reference concordance and heterozygous FPR were excluded from subsequent analysis.

### Downstream analyses

To assess and quantify potential biases introduced by imputation already known to occur (Erven et al. 2022), we performed comparisons between high coverage samples and their imputed counterparts. Both genotype-based analyses, such as Principal Component Analysis (PCA), Runs-of-Homozygosity (ROH) analysis, and the haplotype-based Identity-by-Descent (IBD) analysis were performed to assess imputation accuracy. Reduced concordance and high FPR outlier regions (Tables S12-13) were excluded from downstream analyses.

As a preliminary assessment of the imputed genotypes we performed PCA, comparing high coverage and pseudohaploid data (Figures S22-24). We found consistency with previously-reported affinities, but also observed a large effect of missingness in PCA placement, suggesting that using lower GP thresholds on imputed data may be justified to ensure adequate site number (Supplementary Note 5).

### Outgroup *f_3_* statistics

As a test case of potential applications of using imputed goat palaeogenomes in concert with globally-distributed modern genotypes, we assessed the genetic history of breeds today and their relationship to ancient populations. Evidence from modern goat breed genotype data suggests that the continental gene pool of Europe, Asia, and Africa were primarily established by the dispersal of managed herds from Southwest Asia, with some subsequent admixture events (Colli et al. 2018). To address this we measured shared genetic drift between ancient and modern genomes in the Vargoats dataset by imputing 36 previously published ancient goat palaeogenomes with ≥0.5✕ coverage (Table S3), and subsequently filtering genotypes (GP99). We then computed outgroup *f_3_*statistics (Patterson et al. 2012), calculated between pairs of ancient genomes and also with modern breed groups (Table S14). We plotted these values on a world map indicating the provenance of each genome or breed grouping; we present a subset of these in Figures S25-31 and the remainder at 10.17605/osf.io/av4f9.

The shared drift measurements with modern domestic goats allowed a broad description of how ancient goat genomes relate to breeds today. We observe for the first time that African breeds, particularly those in East Africa, showed greater affinity with a Bronze Age goat from Israel (Yoqneam2; Figure S25) and Central Türkiye (Acem2; Figure S26), likely reflecting their initial dispersal from Southwest Asia (Marshall & Hildebrand 2002; Prendergast et al. 2019). Among African breeds, there are differential affinities to Neolithic genomes. Goats from North and West Africa show greater affinity with Neolithic European genomes, while breeds from East Africa show more affinity with ancient Iranian and Turkish goats, similar to patterns observed from modern data (Colli et al. 2018). Asian goats today, particularly those from Bangladesh, showed high affinity with the Neolithic East Iranian genomes (e.g. Semnan3; Figure S27); goats from South Asia may therefore better represent the ancestry of animals during the initial dispersal of livestock herds into Central and eastern Asia.

There are hints in our data that this may be confounded by admixture with wild bezoar, with modern Asia breeds having low but varying affinity with a pre-domestication genome from Armenia (Hovk1; Figure S28); there are some evidence of admixture from other *Capra* species into domestic goat (Zheng et al. 2020; Li et al. 2022). Breeds from Europe or having European origin showed affinity with the Neolithic Serbian genomes (e.g. Blagotin3; Figure S29), while a Late Bronze Age goat from Potterne, Wiltshire (UK) showed highest affinity with Old Irish Goat (Figure S30). Finally, ancient bezoar showed higher affinity with European goats and least with those from East Asia (Figure S31), in line with previous inferences of Turkish bezoar admixture in West Eurasian domestic genomes (Daly et al. 2018, 2022). Shared drift between pairs of ancient genomes recapitulates other indicators of affinity, including broad affinity among post-Neolithic Iranian genomes and notable affinity of the Bronze Age goat from Israel (Yoqneam2) to post-Neolithic genomes across Southwest Asia.

### Testing runs-of-homozygosity inference

The recovery of imputed diploid genotypes permits the direct testing of the domestication bottleneck hypothesis by tracking the burden of runs-of-homozygosity (ROHs) through time. To do this we first optimized ROH inference by assessing the parameters affecting their estimation in our imputed genotypes. We computed ROHs with no missingness across test samples simultaneously using transitions and transversions (“all sites”) or with transversions-only. We additionally computed ROHs on an individual-basis using transitions and transversions or transversions-only. For each dataset, we further downsampled to 1M, 750K and 500K sites to determine a minimal number of SNPs necessary for accurate ROH calculation. The non-imputed high coverage samples were used as validation of ROH profiles and were computed without downsampling on an individual-basis using either transitions and transversions or transversions-only. Exploring the effect of site number of ROHs (Figure S32, Supplementary Note 6), we opted to calculate ROHs on datasets >750K sites with a minimum of 200 SNPs per ROH, using both the combined datasets with no missingness and individual-based ROH calculation.

We tested the consistency between imputed and high coverage genotypes, when no missingness is allowed within the dataset (i.e. when the same sites are analyzed in high coverage and imputed genotypes). We find high consistency between the imputed and high coverage ROH profiles, indicating that false-positive and false-negative heterozygotes have less effect on ROH profiles at a GP threshold of 0.99 (Figure S33). A similar pattern is observed when using bcftools to calculate ROHs, which produce similar profiles to plink for high coverage and 0.5✕ imputed genomes assessed together with no missingness (Figure S34), although we find instances of erroneously fragmented long ROHs (Figure S35-38, other sample figures available at 10.17605/osf.io/av4f9).

Given the robust parameter space identified, we computed ROHs using imputed ≥5% MAF genotypes of goat palaeogenomes with ≥0.5✕ coverage, applying a GP99 filter and removing sites with any missingness across samples (Figure 5, Table S3, S15). We compared these combined ROH profiles to ROH profiles calculated from transversions-only, with ROHs calculated on a per-individual basis (Figure S32, Supplementary Note 6), downsampling each individual to a random 1.2M and 1.6M sites (matching the lowest number of sites across samples for 0.4✕ and 0.5✕, respectively – Table S15). We observed large differences between ROH profiles generated using plink and bcftools in the combined dataset (Figure S39). Using ROHan to infer reference ROH profiles (Renaud et al. 2019), plink consistently detects fewer ROHs, especially long ROHs (>8Mb). In contrast, bcftools overestimated short ROHs, but correctly called longer ROHs based on comparison to heterozygosity profiles and ROHan profiles (Figure S35-39). These discrepancies were diminished when using higher SNP density datasets (1.6M sites; Figure S40). Analysis of local ROHs and heterozygosity profiles across the different datasets suggests that ROH fragmentation in the combined dataset is likely due to lower density of SNPs rather than an increase of heterozygous SNPs (Figures S36-38), leading to the lower number of long ROHs called by plink. These findings highlight the importance of SNP density in ROH estimation, the potential underestimation of long ROHs when using plink, and overestimation in small ROHs using bcftools.

**Figure 5:**
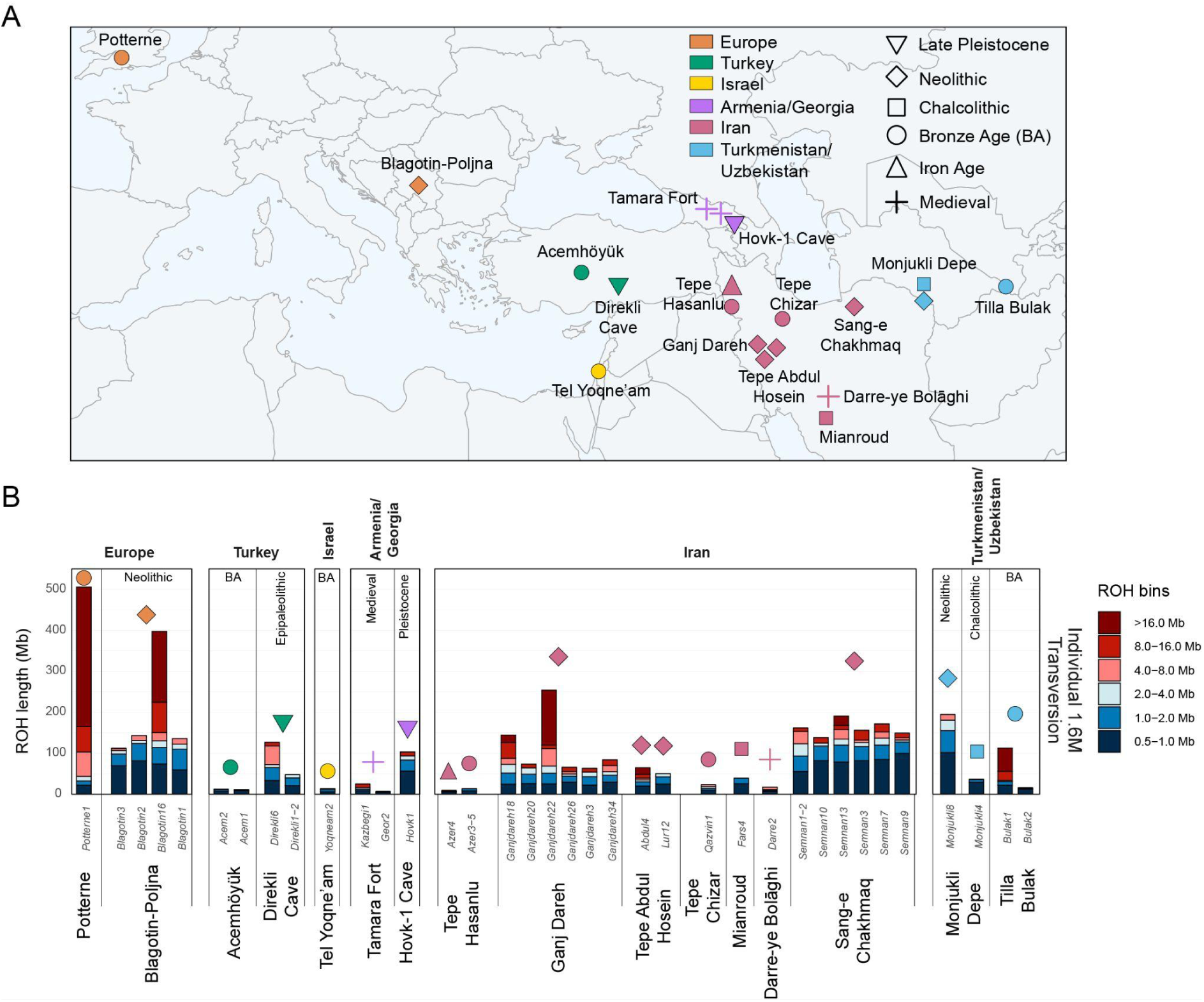
Analysis of ROH for imputed ancient goat genomes with ≥0.5✕ mean coverage. Key indicates the general archaeological period the samples derived from. A) Map of archaeological settlements from which imputed genotypes are analyzed. B) ROH length bin profiles for ≥0.5✕ imputed ancient goats. ROH was calculated with each ancient sample individually, using transversions-only and downsampling to 1.6 million sites, which corresponds to the lowest number of imputed sites in the 0.5✕ imputed test samples. ROH profiles are the result of combining plink ROH and long ROH (>4Mb) from bcftools.

To mitigate these opposing biases we combined approaches, merging bcftools-identified long ROHs (>4Mb) with plink-computed ROHs. This integrative approach resulted in similar ROH profiles for individual-based transversions (Figure 5B) and “combined” transition and transversions datasets (Figure S41). However, it is important to note that bcftools when applied on limited sites (combined dataset: 897K sites) may overcall long ROHs (Figure 5, S35), and results with lower site numbers should be interpreted with caution.

### Spatio-temporal patterns of runs-of-homozygosity in goats

Having established a pipeline for ROH inference, we investigated how these genomic signatures of inbreeding varied during and after the initial phases of goat domestication. Assessing computed ROH patterns, goats from Ganj Dareh, a settlement with some of the earliest known evidence of livestock management ∼8,200-7,600 cal BCE (Zeder & Hesse 2000; Daly et al. 2021), typically show a total ROH amount of <100Mb, with individual ROHs occurring across length categories (Figure 5B). In contrast, two genomes were outliers in having longer ROH stretches (Ganjdareh18 and 22). The long stretches of ROHs in these samples were previously identified (Daly et al. 2021) and linked to potential kin matings within a management-induced small breeding population. In contrast, the overall genetic diversity in Ganj Dareh has been reported as high relative to other ancient goats (Daly et al. 2021). Other Ganj Dareh goats (Ganjdareh3, 20, 26 and 34) showed a low overall cumulative sum of ROHs (Figure 5B). Genomes (Abdul4 and Lur12) from the nearby Tepe Abdul Hosein (inhabited ∼8,250-7,800 and ∼7,600-7,550 cal BCE, (Nishiaki 2022)), show a comparable ROH profile to these lower ROH Ganj Dareh samples.

Goats from the Northeast Iranian settlement of Tappeh Sang-e Chakhmaq (Roustaei et al. 2015; Mashkour et al. 2014), with both Aceramic (∼7,000 cal BCE; sample Semnan1-2 in Figure 5B) and Ceramic (∼6,000 cal BCE; remaining Semnan samples) phases, showed significantly elevated ROH profiles compared to those from the earlier site of Ganj Dareh (Mann-Whitney U test; U=7, one-tailed *p*=0.01465). Furthermore, no outliers with large cumulative sums of long ROHs were present, with analyzed genomes showing an inflation of ROHs across size categories. These consistent elevated ROH profiles are indicative of a bottleneck during the dispersal from a region of domestication and beyond the range of their wild progenitor *C. aegagrus* (Zheng et al. 2020; Daly et al. 2021; Zeder 2024; Petretto et al. 2024).

Following the Neolithic period, goats from archaeological assemblages across Iran, Türkiye, Israel, Armenia, Georgia, Turkmenistan, and Uzbekistan exhibited a reduced cumulative sum of ROHs. For instance, at Monjukli Depe in southern Turkmenistan, the ROH profile of Monjukli8, a sample from Neolithic layers (6,400-5,800 BCE), closely resembled that of contemporaneous goats from Tappeh Sang-e Chakhmaq. This pattern shifts in the subsequent Chalcolithic period (5,100-4,500 BCE; also known as Aeneolithic) as evidenced by a later genome from Monjukli Depe (Monjukli4), which displayed a lower cumulative sum of ROHs. This pattern, observed across Southwest Asia, indicates that herd genetics were shaped by increased outbreeding or a sustained lack of inbreeding over the intervening millennia. Alternatively, bezoar brought under human control may already carry inbreeding signals, due to their non-panmictic polygynous reproduction system (i.e. isolated populations with male competition prior to reproduction). Notably, contrasting ROH profiles can be observed within goats in Bronze Age Uzbekistan (Tilla Bulak, ∼2,000-1,700 BCE), where Bulak1 shows an excess of long ROHs compared to Bulak2. Additional samples from Central Asia are required to draw broader regional conclusions.

ROH profiles from the westward dispersal of managed goats to Europe also suggested a population bottleneck. At Blagotin-Poljna (Serbia) ∼6,000 cal BCE, we saw one outlier (Blagotin16) with a high amount of long ROHs, indicative of recent inbreeding. While this is not observed in the other Blagotin-Poljna goats, their overall summed ROHs (average ∼197 Mb) was larger than those from the Central Zagros Mountains (8,200-7,600 cal BCE; on average ∼100 Mb), however this was not statistically significant, likely due to the small sample size (Mann-Whitney U test; U=6, one-tailed p=0.05455). Our dataset contained one European sample postdating the Neolithic in Europe, Potterne1 (Potterne, Wiltshire, UK; ∼800 cal BCE), which showed a notable excess of long ROHs, suggestive of recent inbreeding.

### Identity-by-Descent

To assess our ability to recover shared chromosomal segments we identified Identity-by-Descent (IBD) regions (Supplementary Note 7, Table S16). Aware of limitations in our IBD recovery for ancestries not well represented in the phasing panel (e.g. Direkli1-2), we analysed pairwise IBD sharing among imputed genotypes of goat palaeogenomes with ≥0.5✕ (Table S3), applying a GP99 and MAF 5% filter, to compare within-settlement relatedness and to evaluate IBD sharing between-settlements. We restricted this to the Neolithic period due to the low sample size available for other time periods (Table S3). Normalized relatedness was calculated for settlements with a minimum of three samples. Individuals were considered related if they shared at least three IBD segments >3cM, with a cumulative IBD length of at least 21cM.

Normalized relatedness was calculated for Ganj Dareh (Iran, ∼8,200-7,600 cal BCE), Tappeh Sang-e Chakhmaq (Iran ∼7,000-6,000 cal BCE), and Blagotin-Poljna (Serbia ∼6,000 cal BCE), shown in Figure 6A. Among these settlements, Tappeh Sang-e Chakhmaq showed the highest normalized relatedness (1), indicating that all of the tested individuals were related (Table S17). We excluded from these group comparisons the single genome from the Aceramic West Mound (∼7,000 cal BCE) at Tappeh Sang-e Chakhmaq, Semnan1-2, which incidentally has short IBD with the ∼6,000 cal BCE East Mount goats (Table S18). The Blagotin-Poljna assemblage also showed a relatively high normalized relatedness (0.55), indicating that roughly half of the individuals were related (Table S17). In contrast, goats from Ganj Dareh showed a normalized relatedness of 0, indicating no detectable genetic relatedness among individuals.

**Figure 6:**
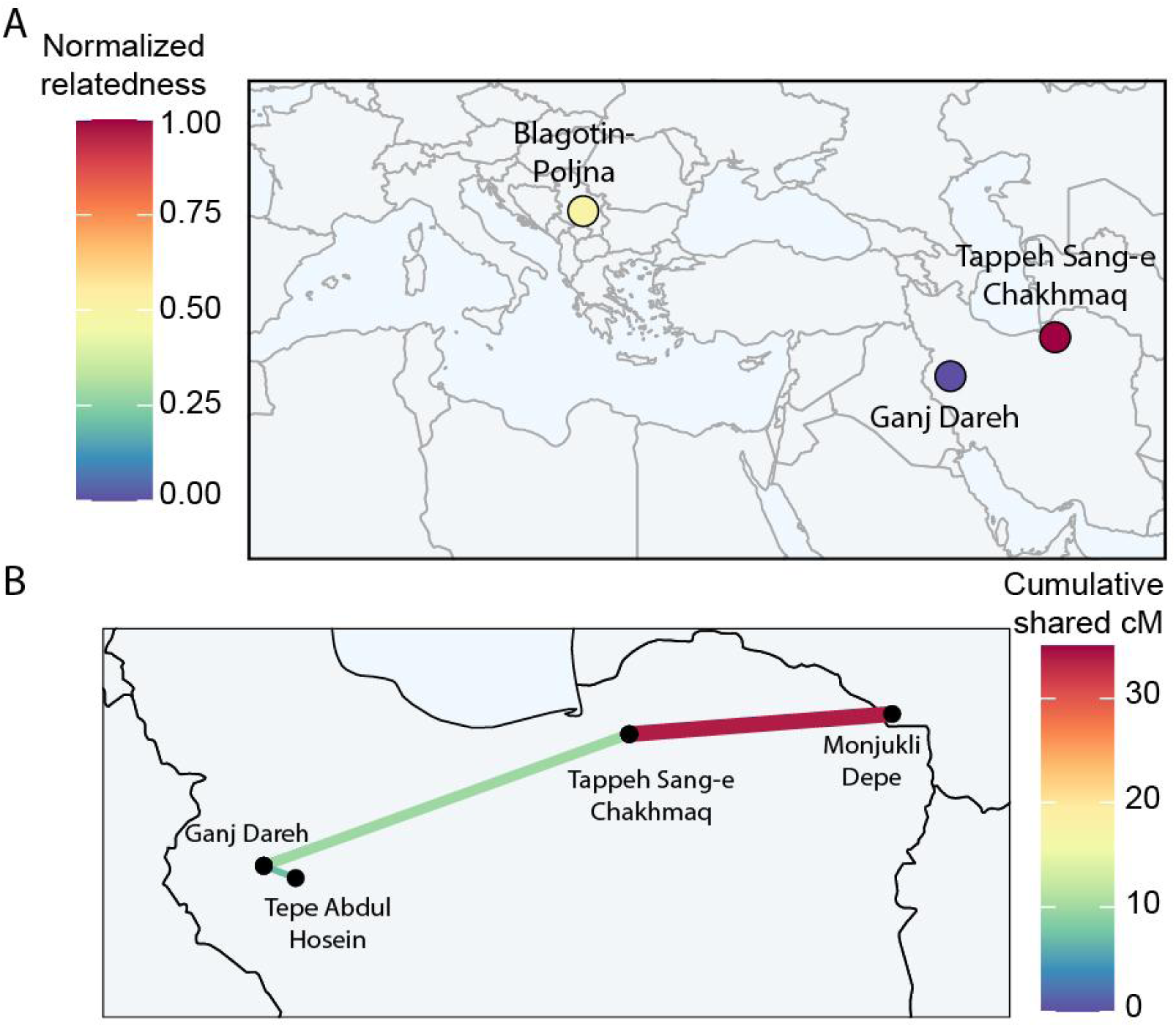
IBD sharing A) within-settlement, normalizing relatedness for assemblages with ≥3 imputed genomes with ≥0.5✕ and B) between-settlement cumulative shared IBD (cM). For Tappeh Sang-e Chakhmaq, we excluded a single ∼7,000 cal BCE individual from the West Mound, Semnan1-2. Between-settlement IBD is represented by lines between-settlement pairs, and does not imply direct cultural connectivity or dispersal patterns.

Between-settlement IBD broadly matched geographic proximity (Figure 6B), with little long-range IBD detected. We saw notable IBD sharing in Neolithic East Iran, specifically between Tappeh Sang-e Chakhmaq and Monjukli Depe in nearby Turkmenistan (Figure 6B; the older Semnan1-2 genome also showed IBD sharing with Monjukli8, see Table S18), sites showing cultural affinities (Roustaei et al. 2015). IBD sharing between Tappeh Sang-e Chakhmaq and Ganj Dareh was notably lower, suggesting a weaker connection at least partially driven by the temporal difference (∼2,200-600 years) between the assemblages.

## Discussion

We report the successful imputation of ancient goat genomes. Using the large-scale VarGoats reference datasets and GLIMPSE2 (Rubinacci et al. 2023), non-reference concordance rates for both transition and transversion variants reach 0.7 for the lowest tested coverage and GP (0.1✕, at GP70 – Figures 2-3, Table S7). Once stratified for common variants (MAF ≥5%) and GP99, concordance increases to >0.9 at the lowest tested imputation coverage (0.1✕). Higher MAF thresholds (i.e. 10%) produce similar concordance values to those seen at 5% (Figure 2, Table S7). For genomes imputed here, concordance is >0.99 at 0.5✕ for common variants. Notably, we imputed common variants in an Epipaleolithic wild bezoar (Direkli1-2) with a non-reference genotype concordance of 0.97 at 0.4✕ and 0.5✕, demonstrating that variation shared between wild and domestic populations can be accurately imputed even with a largely-domestic reference panel. This suggests that ancient domestic populations not studied here will impute with similar accuracy using the VarGoats reference genotypes, when restricted to common variants (≥5% MAF).

We note some limitations in our imputation approach. Variants below the frequency threshold of 5% impute less accurately (Figure 2), particularly at lower coverages; studies across species have observed that alleles less common in a reference panel are more difficult to impute (Sousa da Mota et al. 2023; Erven et al. 2024; Bougiouri et al. 2024; Yao et al. 2025). As a consequence, variants unique to ancient populations or rare today (that is, having a MAF below 5%) are not recovered accurately (Figures 2, 3). We additionally observe a difference between transitions and transversions in rare alleles (Figure 2), also reported in ancient humans and dogs (Sousa da Mota et al. 2023; Bougiouri et al. 2024) but to a lesser extent. A potential reason for this could be differences in how genotype concordance was calculated – comparing all genotypes versus only heterozygous and homozygous alternatives in this study. Alternatively, those imputation reference panels may have better representation of the ancestry of tested samples compared to the present study, including at rarer alleles. Future work should consider incorporating high quality ancient genomes into imputation reference panels, thus making the panel more representative of the divergent ancestry to be imputed, potentially improving accuracy (Escobar-Rodríguez & Veeramah 2024).

Filtering criteria for imputed genotypes necessitate a trade-off between genotype concordance and the number of genotypes available for analysis. Heterozygous errors - both spurious (FPR) and missed (FNR) - are apparent in our data (Figure 4). We find stringent GP thresholds can mitigate these errors, at the cost of genotype number. As such, we recommend that for this and comparable datasets, the GP threshold be tailored to the downstream analysis. For analyses in which accuracy is paramount, such as ROHs or IBD, a high GP filter is necessary to detect shared haplotypes or regions depleted of variants. For PCA analyses, we find missingness to be a larger source of error than imputation (Supplementary Note 4); a lower GP filter such GP80 or GP90 may therefore be reasonable.

We demonstrate the potential of imputed genotypes to infer ROH profiles accurately and report broad trends of inbreeding in this ancient livestock. Our exploration of plink-based ROH parameters suggest individual-level ROH calculation, restricted to transversion genotypes and matching variant numbers by random downsampling to match the sample with the lowest variant count, captures many of the same patterns as a “combined” dataset of all individuals. This may circumvent an issue of “combined” ROH calculation, which is limited by decreasing site number when a “no missingness” filter is used and the number of individuals increases. In general, our testing highlights the sensitivity of plink-calculated ROH to variant density and to the parameters used (Supplementary Note 5). Contrasting this are ROH profiles inferred using bcftools, which appear to accurately recover longer ROHs but can over-call short homozygosity stretches (Figure S36-38). We therefore suggest a novel approach to ROH estimation, by merging long (≥4 Mb) bcftools-computed ROHs with plink-computed ROHs, mitigating the opposing bias of both tools.

Our initial description of the inbreeding landscape of ancient goat populations (Figure 5) reveals distinct patterns during the Neolithic period: increased inbreeding and total ROHs in genomes from locations which managed goats were introduced to, as observed in ancient cattle (Rossi et al. 2024). This finding is consistent with a growing body of palaeogenomic-informed research (Allaby et al. 2019; Frantz et al. 2020) indicating that domestication of many species involved multiple wild populations, geographically and temporally dispersed (Daly et al. 2018; Verdugo et al. 2019; Frantz et al. 2019; Bergström et al. 2022; Kaptan et al. 2024; Erven et al. 2025). Some signals of inbreeding are detected within Aceramic Neolithic assemblages in the Central Zagros Mountains as previously reported (Daly et al. 2021), indicating a degree of management-induced or standing inbreeding level. In contrast, ROH lengths rise as managed herds are transferred to more distant locales, such as Southeast Europe (Blagotin-Poljna) and Northeast Iran (Tappeh Sang-e Chakhmaq), providing parallel direct evidence for the accumulation of inbreeding in a livestock species during its human-induced dispersal in the Neolithic period. This may have been further exacerbated by the reduction or lack of gene flow with wild bezoar populations, particularly in Europe where *C. aegagrus* was not present.

Today, goat breeds from more isolated regions show elevated ROH signals (Bertolini et al. 2018), illustrating the general consequences of limited gene flow. For example, goats today from Ireland and Britain (and including a single Bronze Age genome from Potterne, Wiltshire, Figure 5) are among the least genetically diverse of breeds surveyed (Petretto et al. 2024), suggesting prolonged genetic isolation. Inbreeding signals appear notably reduced - with some exceptions - from the Chalcolithic onwards. This may be due to a combination of many factors: improved husbandry practices, growing effective population sizes, and developed animal trade networks. Additional genomes appropriate for imputation, from regions and time periods not represented in this study, will shed light on the overall trend and regional variation of this observation.

Our application of IBD analysis using imputed genotypes (Figure 6) echoes the ROH findings, and together develop a more complete picture of the early demographic history of managed goats. By inferring shared haplotypes between genomes, IBD provides extra information about shared relatedness between genomes: within a population or settlement, or between those population pairs. The length of these shared haplotypes allows the inference of the age since a shared ancestor, identify cryptic relatedness otherwise missed using summary statistics, construct pedigrees, track ancestral haplotypes, and reconstruct ancestral migration paths.

The absence of detectable relatedness in the Aceramic Neolithic settlement of Ganj Dareh, or between Ganj Dareh and the nearby (∼60km) contemporaneous Tepe Abdul Hosein (Figure 6B) is in itself notable, given the demographic and genetic evidence of herd control (Zeder & Hesse 2000; Daly et al. 2021). This may be attributed to an extended initial phase of domestication, during which managed stock were more freely interbreeding with the unmanaged, genetically distinct animals (Daly et al. 2021). The long inhabitation period of both settlements (Ganj Dareh: 200-600 years, Tepe Abdul Hosein, ∼700 years (Meiklejohn et al. 2017; Daly et al. 2021; Nishiaki 2022)) may also explain the lack of within- and between-settlements IBD detection, although we do find IBD between pairs at Tappeh Sang-e Chakhmaq separated by ∼1,000 years (Table S3, S18). The absence of detected Ganj Dareh-Tepe Abdul Hosein IBD could be taken as evidence for limited animal exchange between the two settlements, or that such exchange was small relative to the herd sizes, potentially inflated with recruitment of local wild goats.

In contrast, herds later in the Neolithic period, in regions into which herded goat were introduced such as Europe (Blagotin-Poljna) or Northeast Iran (Tappeh Sang-e Chakhmaq), show greater within-settlement IBD, and indications of regional between-settlement IBD (Monjukli Depe, Figure 6). These may reflect “founder herd” effects, i.e. the establishment of local Neolithic herds from a narrow pool of animals, during the centuries-long expansion of agriculture in Eurasia. Such herds may have limited outbreeding opportunities, with nearby settlements’ flocks also established from the same founders. Indeed, both sheep and goat breeds today show an inverse relationship between genetic connectivity and IBD (Blondeau Da Silva et al. 2024).

## Conclusion

Our analyses confirm the potential of imputation methods to leverage emerging large-scale non-human genome datasets and maximize genotype recovery in palaeogenomes for common alleles (MAF ≥5%). Rare alleles (MAF <5%) showed notably reduced imputation accuracy, highlighting the importance of haplotype and ancestry diversity within the reference panel being at least partially representative of the target genomes. Future work could explore how ancient genomes can be included within imputation panels, possibly through joint imputation of ancient samples (while circumventing batch effects). Such efforts will be crucial in broadening the scope of which species ancient imputation can be applied. The observed trade-off between genotype number and genotype accuracy must also be grappled with depending on the intended question and analysis.

With stringent filtering and a minimum coverage of 0.5✕, we show that ROH and IBD can be recovered from genomes ∼10,000 years old, and ROHs in wild bezoar genomes preceding evidence of goat husbandry. Imputed genotypes here have facilitated an initial picture of the demographic processes of the first millennia of goat management. Notably, parallel founder events are inferred in goats later in the Neolithic process; demographic costs of domestication may be more ascribed to dispersal rather than initial herd management. Imputed genotypes will likely facilitate other haplotype-based analysis of ancient livestock, such as Local Ancestry Inference, to build an increasingly detailed picture of how human activity has impacted the genetic trajectory of partner species.

## Methods

### Modern genotypes

The VarGoats project dataset is currently composed of 1,372 animals across 155 country-breed groups and 7 wild *Capra* species (Table S1), including bezoar. Genotypes were called across all samples as per (Denoyelle et al. 2021), resulting in a set of 77 280,295 SNPs. This set of SNPs was further filtered, following 3 steps: i) 85 animals were removed if their Whole Genome Sequencing parameters were harbouring MeanDP <6, MeanGQ <19 and %Missing >10 or their 60k genotyping data identified them as outliers in PCA and Admixture analysis (i.e. mislabelled animals, data not shown). ii) only biallelic variants with GQ>20 and subsequent CR>0.9 were retained. iii) the remaining 28,645,747 high quality SNPs were recovered for the entire dataset. Details about this procedure could be found in [paper in prep]. Phasing was performed using Beagle 5.3 (Browning et al. 2021). Details about the methodology and relevant codes are available at https://github.com/goatimpuation/GoatWGSimputation.

To account for individuals with high genomic relatedness, prior to imputation individuals with pairwise π^HAT^ >0.4 with more than 2 individuals were removed using plink v1.07 (Purcell et al. 2007). For any remaining pair with π^HAT^ >0.4, a random member of the pair was removed, leaving 1,155 individuals in the imputation reference panel (Table S1). VCF files were then processed following the GLIMPSE2 documentation (Rubinacci & Delaneau).

### Ancient genomes

To evaluate the potential of GLIMPSE in the imputation of ancient goats, we performed a downsampling experiment on four high coverage goat genomes (Daly et al. 2018). These represented distinct time periods and ancestries, offering different perspectives to the reference panel’s representation of past diversity (Table S1-2). Direkli1-2 (mean genome coverage 11.21✕), a wild *C*. *aegagrus* from the Epipalaeolithic faunal assemblage at Direkli Cave in the Taurus Mountains of southern Türkiye (Arbuckle & Erek 2012), dates to the late 12th millennium BCE, prior to evidence of goat herd management. Two genomes represent herds early in their dispersal beyond their initial domestication region: Semnan3 (14.44✕), a ∼6,100 BCE goat from the West mound of Tappeh Sang-e Chakhmaq (Roustaei et al. 2015; Mashkour et al. 2014) in Northeast Iran; and Blagotin3 (11.40✕), a goat dating to ∼6,000 BCE from Blagotin-Poljna in Serbia (Greenfield & Jongsma-Greenfield 2014). Finally, Acem2 (8.63✕) represents an admixed or heterogenous ancestry from the Central Turkish settlement of Acemhöyük, and dates to ∼1,900 cal BCE.

The four high coverage goat bams, aligned to the ARS1 reference genome (Bickhart et al. 2017) following steps described previously (Daly et al. 2021), were individually downsampled to a range of coverages using samtools view: 0.1✕, 0.15✕, 0.2✕, 0.25✕, 0.3✕, 0.4✕, 0.5✕, 0.75✕, 1✕, 2✕, and 4✕. All bam files of ancient goats were soft-clipped 4bp at either end of reads using a python script prior to analyses, as per (Daly et al. 2021). Mpileup files were generated for the high coverage and downsampled bams for the 28,242,942 autosomal variants in the VarGoats call set using samtools mpileup (v1.13, parameters -B -q 30 -Q 30, (Li et al. 2009)) and bcftools call (v1.13, parameters “-mO -C alleles”, (Danecek et al. 2021)). The high coverage validation dataset (i.e. full coverage) was further filtered for Genotype Quality (GQ) of 30 and for sites with ≥6 and ≤40 covering reads using a custom python script, producing diploid genotypes against which imputed genotypes were compared.

The range of downsampled coverages were pseudohaploidized using ANGSD version 0.938 (Korneliussen et al. 2014) doHaploCall, with the following parameters: doHaploCall 1, doCounts 1, dumpCounts 1, minimum base quality of 30 (-minQ 30), minimum mapping quality of 25 (-minMapQ 25), retain only uniquely mapped reads (-UniqueOnly 1), remove reads flagged as bad (-remove_bads), remove triallelic sites (-rmTriallelic 1e-4), downscale mapping quality of reads with excessive mismatches (-C 50), discarding 5 bases of both ends of the read (-trim 5), and remove transitions (-rmTrans 1). The abovementioned sites in the VarGoats reference panel were used as input for ANGSD using the parameter –sites. As a sanity check, the major/minor alleles of the low coverage ancient were compared to the modern reference panel and were removed if there were any discrepancies. ANGSD haplo files were transformed to plink tped files with the haploToPlink function from ANGSD version 0.938 and recoded into ped files with PLINK v.1.90 (Chang et al. 2015). Transitions were removed because transitions, unlike transversions, are affected by postmortem deamination of DNA, which might increase individual error rates.

Previously published goat palaeogenomes with ≥0.5✕ coverage (Table S3) were aligned to the ARS1 reference genome (Bickhart et al. 2017) as described previously (Daly et al. 2021). All bam files of ancient goats were soft-clipped 4bp at either end of reads using a python script prior to analyses, as per (Daly et al. 2021).

### GLIMPSE imputation

Prior to imputation with GLIMPSE2 v2.0.0 (Rubinacci et al. 2023), 4Mb chunk files were generated for ARS1 using GLIMPSE2 chunk (--window-mb 4 --buffer-mb 0.2), utilising a sex-averaged megabase-scale recombination map (Etourneau et al. 2025). The ARS1 reference fasta was split using GLIMPSE2_split_reference and the recombination map. Each downsample bam file was then separately imputed using GLIMPSE2 using the chunk files and recombination map described above. The resulting bcf files were ligated using GLIMPSE2_ligate and converted to vcf files using bcftools convert. Previously published goat palaeogenomes (Table S3) were also imputed separately.

The same steps and parameters were used for GLIMPSE1 (Rubinacci et al. 2021) following their guideline (https://odelaneau.github.io/GLIMPSE/glimpse1/), GLIMPSE1 was performed individually on the downsampled bcftools generated genotypes.

#### Sheep/outgroup genotypes

For outgroup *f_3_* analyses, an outgroup population was created by aligning domestic sheep to the goat reference ARS1. Ten domestic sheep or wild Asiatic mouflon (Table S4) were aligned to ARS1 using bwa mem (Li 2013). Genotypes were called using the same pipeline as used for the validation of ancient genotypes. Genotypes were then filtered for GQ30 and for sites covered by 6-40 reads. VCF files containing the VarGoats reference panel and sheep genotypes were then merged with bcftools.

#### Validation of genotype imputation

To evaluate the effect of post-imputation quality filtering, a range of genotype probability (GP) filters (0.7, 0.8, 0.9, 0.95, 0.99) were applied to the downsampled-imputed genotypes using a custom python script, with genotypes failing the filtering set to missing (”./.”). For each of the four higher coverage individuals, the number of matching sites and concordance rates between validation genotypes and the GP-filtered imputed genotypes of varying coverages and GP filters were calculated using a custom R script; only heterozygous and homozygous-alternative sites (non-reference) in the validation dataset. To assess the effect of minor allele frequency (MAF, here representing the frequency in the Vargoats 1372 individual panel) on imputation accuracy, concordance was calculated at different minimum MAF thresholds: 0%, 1%, 5%, 10% and in MAF tranches: 0-1%, 1-2%, 2-5%, 5-10%, 10-20%, 20-50%.

Non-reference concordance was calculated by dividing the total number of correctly imputed heterozygous and homozygous alternative genotypes by the total number of heterozygous and homozygous alternative genotypes in the validation dataset. Heterozygous false-positive rate (FPR) was calculated by dividing the number of imputed heterozygous called as homozygous in the validation dataset (FP) by the sum of FP and the number of correctly-imputed homozygous sites (TN). The false-negative rate (FNR) was calculated by dividing the number of imputed homozygous called as heterozygous in the validation dataset (FP) by the total sum of called heterozygotes in the validation dataset.

#### Local concordance and FPR

Non-reference concordance and FPR throughout the genome were calculated in sliding windows of 500 kb with a step size of 100 kb; sliding windows were created with BEDTools (Quinlan & Hall 2010) makewindows. Sliding windows consisting of correctly imputed and total number of genotypes were created with BEDTools map, using the sliding windows (500 kb with a step of 100 kb) created earlier. This approach was repeated for the FPR. The outlier non-reference concordance and FPR regions were extracted by filtering for the bottom and upper 1 percentile, respectively. These outlier regions were then merged with BEDTools merge. Genes in outlier regions were determined using RefSeq annotation GCF_001704415.1_ARS1, downloaded from UCSC genome browser (last accessed 29-05-2018), using BEDTools closest. To determine regions that were outliers in the majority of the samples, BEDTools merge was used on the individual bed files, regions that overlapped were counted, with the parameter -o sum, and these regions were subsequently filtered to only be present in a majority of the samples (at least three). A Gene Ontology (GO) enrichment analysis was performed to identify overrepresented biological processes and molecular functions of the genes in each individual outlier region. GO enrichment analysis was done using both g:Profiler (version: e112_eg59_p19_25aa4782 –https://biit.cs.ut.ee/gprofiler/gost) and STRING-DB (Version 12.0 – https://string-db.org/) both accessed on 13-03-2025.

### PCA

The downsampled pseudohaploid ancient samples were merged with the data set containing the imputed and high coverage genotypes and VarGoats reference panel, filtered for MAF 5% with a missingness filter of 1.5%; this was done for imputed samples filtered for GP99, GP95 and GP80, separately. Smartpca version 16000 was used with default parameters (Patterson et al. 2006; Price et al. 2006). The first ten principal components were calculated using the modern reference panel; the pseudohaploid, imputed, and high quality genotypes were projected (lsqproject:yes, autoshrink:yes).

### Outgroup f_3_

To estimate the degree of shared genetic drift between modern goat breeds and ancient genomes, outgroup *f_3_* statistics (Patterson et al. 2012) were calculated using ADMIXTOOLS version 7.0.2 between imputed genotypes of each ancient genome with ≥0.5✕ mean coverage, and each domestic breed grouping in the VarGoats dataset. We used more granular groupings (i.e. “Distinct_split-off_breed”) where available, and breed groupings otherwise (Table S1). Domestic sheep were used as the outgroup (Table S4).

### ROH

We performed ROH exploration on our downsampled 0.5✕ imputed genomes using four datasets filtered for MAF ≥5% and GP99. ROH were computed without missing data across test samples using either transitions and transversions (“all sites”) or with transversions-only. Additionally, individual-based ROH calculations (i.e. one ancient sample at a time, using sites passing filtering steps for that sample) were conducted using either all sites or transversions-only.

For each dataset, we further downsampled to 1M, 750K and 500K sites to determine a minimal number of SNPs required for accurate ROH estimation. The non-imputed high coverage samples were used as validation ROH profiles and were computed on an individual-basis using either all sites or transversions-only, without additional downsampling. ROHs were estimated with PLINK v1.90 (Chang et al. 2015) with the parameters –homozyg --homozyg-density 50 --homozyg-gap 100 --homozyg-kb 500 --homozyg-snp 50 --homozyg-window-het 1 --homozyg-window-snp 50 --homolog-window-threshold 0.05, according to earlier studies (Gamba et al. 2014; Cassidy et al. 2016; Martiniano et al. 2017; Sousa da Mota et al. 2023). The minimum SNP threshold (--homozyg-snp; default of 50) was changed to 150, 200, 250 and 300, to test the effect on small ROHs within our dataset. We further explored ROH profiles for 0.25✕, 0.5✕, 0.75✕, and 1✕ imputed genomes on a per-sample basis, using the parameters described above, with --homozyg-snp set to 200.

Additionally, ROHs were computed using imputed genotypes from published goat palaeogenomes ≥0.5✕ coverage, appling a MAF ≥5% and GP99 filter. Two approaches were tested: one combined dataset with no missing data and another using transversions only on a per-individual basis, with each individual downsampled to 1.2 and 1.6 million sites (matching the lowest number of sites for the 0.4✕ and 0.5✕ imputed genomes, Table S3) to ensure consistent SNP density.

Bcftools ROH was used as a complementary method for ROH detection and run with standard parameters aside from *-G 30 --AF-dflt 0.4*. The output files were filtered to retain regions containing at least 200 SNPs, a quality score greater than 10, and a minimum length of 500kb. Long ROHs identified by bcftools (>4Mb) were merged with the plink ROH profiles using mergeBed, counting the number and keeping the sizes of merged ROHs with the parameters -c and -o count, collapse. Finally, ROHan (Renaud et al. 2019) was also used as an alternative method for ROH detection for medium to high coverage ancient genomes. The pipeline for ancient samples was followed, by first detecting possible damage, using the estimateDamage.pl script provided with ROHan. ROHan was run on the autosomes with a heterozygosity cut-off similar to the heterozygosity cut-off in (Daly et al. 2021). The parameters used for ROHan were --rohmu 5.7e-5, and the deamination profiles from the estimateDamage.pl script.

Heterozygosity was calculated using 50-SNP windows by segmenting the VCF file into consecutive windows of 50 SNPs and counting the number of heterozygous sites within each window. This analysis was performed using a custom AWK script.

### IBD

To identify identity-by-descent (IBD) segments between individuals, we used RefinedIBD (Browning & Browning 2013), which assumes no genotype errors and relies on phased haplotypes. Since phasing accuracy is sensitive to missing data, imputed genomes were filtered for GP >0.99 and individual genomes with >20% missingness were removed. The remaining imputed samples were imputed again and phased together as a single “batch” (i.e. not individually) using Beagle5 with standard parameters aside from setting impute=true and specifying a random seed. The second round of imputation with Beagle5 was performed to impute and phase individuals jointly, to minimize missingness while optimizing SNP count within the dataset. Genotypes below GP99 were set to missing, and SNPs were further filtered to keep only bi-allelic SNPs without missing data. We further filtered variants which were MAF 5%, transversions and had no missing data in imputed samples >2✕, resulting in 1,037,536 SNPs. This dataset included imputed 0.5✕ test genomes and previously published ancient palaeogenomes (>0.5✕).

RefinedIBD was run with default settings. To correct for breaks or short gaps in IBD segments, segments were merged with merge-ibd-segments.17Jan20.102.jar with a sex-averaged megabase-scale recombination map (Etourneau et al. 2025), a maximum of one discordant homozygote, and gaps that are less than 0.6 cM in length. The phasing, refinedIBD and merge-ibd were repeated a total of 3 times to account for potential variance in phasing. The first 2 Mb of chromosome 18 was excluded due to a very large estimated recombination rate region on the start of the chromosome, most likely indicative of an assembly error (Etourneau et al. 2025). The repeated runs were combined and filtered for a minimum LOD score of 3 and a length threshold of 3cM. To obtain IBD segments for a pair of individuals, IBD segments for pairs of individuals were extracted from the IBD files and merged with bedtools MergeBed. For IBD detection of imputed 0.5✕ test genomes and their high coverage counterparts both IBD1 and IBD2 segments were considered (indicating that both haplotypes are in IBD), merging IBD segments within 4cM. For previously published ancient palaeogenomes, only IBD1 segments were taken into account without merging IBD segments. Close relatives up to the 8th degree were identified using a threshold of at least 3 shared IBD segments with a cumulative length of at least 21 cM. For within- and between-population comparisons, we excluded the Abdul4 genome from the Tepe Abdul Hosein assemblage, due to its wild genomic affinity (Daly et al. 2021), and Semnan1-2 from the Tappeh Sang-e Chakhmaq assemblage (deriving from the West Mound, (Roustaei et al. 2015)), due to the ∼1,000 year gap between the remaining, younger genomes (from the East Mount).

## Acknowledgments

The authors would like to thank Victoria E. Mullin and Emily Breslin for helpful discussions, and Daniel G. Bradley for mentorship and guidance; Rachel Rupp for work done on the recombination map; and Prof. Akira Tsuneki (University of Tsukuba) for having authorized sampling of the Tappeh Sang e Chakhmaq material from the Japanese excavations of this assemblage. Finally, we thank the VarGoats sample providers acknowledged in Denoyelle et al (Denoyelle et al. 2021).

## Data and script availability

All steps and scripts necessary to replicate the pipeline described are available at https://github.com/JolijnErven/Goat_imputation_manuscript. Imputed unfiltered genotypes are available at <u>10.17605/osf.io/av4f9</u>. Detailed methodology and relevant scripts for the creation of the VarGoats reference panel are available at https://github.com/goatimpuation/GoatWGSimputation. The VarGoats genotypes are under embargo until 31st December 2025, and will then be available at ENA under Project accession PRJEB90141. Previously published data was used for this work (Daly et al. 2021, 2018).

