## Supplementary_Notes_S1-7_Figures_S1-41 for "Inferring domestic goat demographic history through ancient genome imputation"

**Supplementary Note 1 - Sample Specific Effects of Concordance Gains**

A minor allele frequency (MAF) threshold for more common alleles (MAF >5%) results in concordance gains for both heterozygous and homozygous alternative genotypes (Figures S5-7, Tables S7-8). This concordance gain varied between samples, with the ~13,000 year old wild Turkish bezoar goat Direkli1-2 and the ~8,000 year old Iranian domestic goat Semnan3 having a higher concordance gain compared to the ~8,000 year old Serbian domestic goat Blagotin3 and the ~4,000 year old Turkish goat Acem2 (Figures S5-7, Tables S7-8). Concordance for Direkli1-2 at 0.1✕ for all variants with genotype probability GP99 and without a MAF filter were 0.73 and 0.80 for transitions and transversions, respectively. This increased to 0.91 and 0.92 with a 5% MAF threshold (for transitions and transversions; Table S8). This concordance gain between no MAF and a 5% MAF threshold is dosage dependent, where a greater gain is observed in the lower coverages compared to higher (Figures S5-7, Tables S7-8). This concordance gain for common alleles in Direkli1-2 is likely related to the ancestry of this individual, as wild bezoar are underrepresented in the VarGoats dataset relative to domestic animals (16 bezoar total, Table S1). For the 4 test samples (Acem2, Blagotin3, Direkli1-2, and Semnan3), concordance at the lowest coverage (0.1✕) for all sites with GP99 and 5% MAF reaches 0.95, 0.97, 0.92, and 0.96, respectively. This increases to 0.98, 0.99, 0.97, and 0.98 at 0.4✕ coverage, and 0.99, 0.99, 0.97, and 0.99 at 0.5✕ (Table S7), with concordance being slightly higher for transversions only (Table S8). We see similar trends for a minimum MAF of 10% (Table S7-8).

**Supplementary Note 2 - False-Positive Rate and False-Negative Rates**

Direkli1-2 and Semnan3 show a higher heterozygous false-positive rate (FPR) with lower GP threshold; higher GP thresholds (i.e. 0.99) mitigate this sample-specific variation. There is little-to-no difference between transition and transversion FPR, across samples or GP thresholds (Figures 4, S8, Table S8). Interestingly, FPR increases with coverage under stricter GP thresholds, plateauing at 0.5✕. This trend is likely due to the low number of imputed heterozygous genotypes at lower coverages for stricter GP thresholds (<10,000 sites; Table S8). At 0.5✕ and the strictest GP (0.99), the FPR is 1.77%, 1.6%, 1.6% and 1.1% for Acem2, Blagotin3, Direkli1-2, and Semnan3, respectively (Figure 4, Tables S7-8). These roughly correlate with the coverage of the validation genomes, suggesting missed heterozygous calls at lower coverages inflate the measured FPR of imputed genotypes. In contrast, FPR at 0.5✕ for the lowest GP (0.7) is higher, reaching 3.9%, 3.3%, 5% and 3.2% for the same samples (Figure 4, Tables S7-8).

The false-negative rate (FNR) was also computed, measured as the number of imputed homozygous called as heterozygous in the validation dataset (FP) divided by the total sum of called heterozygotes in the validation dataset. Heterozygous FNR decreases for both coverage and stricter GP and starts levelling out at 0.5-0.75✕ (Figure 4). The FNR drops below 1% at 0.75✕ for Acem2, 0.4✕ for Blagotin3, 2✕ for Direkli1-2, and 1✕ for Semnan3, all under the strictest GP threshold (0.99). At the most lenient GP threshold (0.7), the FNR only drops below 1% for Acem2 and Blagotin3 at 4✕ (Tables S7-8). At a GP threshold of 0.95, FNR drops below 1% at 2✕ for Acem2, 1✕ for Blagotin3, 4✕ for Direkli1-2, and 2✕ for Semnan3 (Tables S7-8). This highlights that heterozygous concordance is sensitive to the GP threshold, where a stricter GP limits false-negative and false-positive heterozygotes.

**Supplementary Note 3 - Imputation concordance and recovery; GLIMPSE2 versus GLIMPSE1**

We calculated genotype concordance for GLIMPSE1 to enable a direct comparison with GLIMPSE2. Firstly, we compared GLIMPSE1 using similar filters to GLIMSPE2 (GP filter only), to compare both under the same parameters. GLIMPSE2 achieves higher genotype concordance for non-reference alleles, considering no MAF threshold and low coverages (Figure S12; Table S10). This trend is reversed when looking at a MAF threshold of 5%, where GLIMPSE1 had a higher genotype concordance. These differences are minimal, particularly when looking at the strictest GP filters (0.95 and 0.99). The slightly higher genotype concordance in GLIMPSE1 does not translate to a greater recovery rate; for all GPs and particularly for low coverage, GLIMPSE2 has a higher number of imputed genotypes compared to GLIMPSE1 (Figure S13; Table S10).

We concluded that there is a trade-off between GLIMPSE1 and GLIMPSE2, in terms of the number of recovered sites and concordance rates. Studies must evaluate which outcome - more imputed sites or greater imputation concordance - is most appropriate in each case. Here, we selected GLIMPSE2 as the primary imputation pipeline for our study, as we concluded that the greater number of sites was advantageous for ROH inference across the greatest number of samples.

A previous study showed that the accuracy of GLIMPSE1 improved with the addition of an INFO filter (Erven et al. 2024); we therefore also explored this for the GLIMPSE1 run. A great loss in recovery was observed with the addition of an INFO filter (40-50% recovery loss for 0.5× imputed genomes; Table S10). Moreover, the number of imputed homozygous genotypes stayed constant despite the filtering for varying GP and INFO filters, indicating that the INFO filter was not functioning as intended for homozygous genotypes. We tested an additional GLIMPSE1 run with varying GP filters while restricting the INFO filter for heterozygous genotypes only (Table S11). The recovery of heterozygotes was higher when applying a GP-only filter compared to the combined GP and heterozygote-specific INFO filter (Table S11). Interestingly, heterozygote concordance was also higher under the GP-only filter. In contrast, both the homozygous reference and homozygous alternative genotypes showed higher concordance for the combined GP and heterozygote-specific INFO filter. Moreover, the heterozygous FPR was higher with the GP-only filter, indicating that the INFO filter removes false-positive heterozygotes but also true heterozygotes, highlighting the challenges when filtering post-imputation.

The INFO score is related to the allele frequency of a reference panel (Marchini & Howie 2010); as a result, the INFO score values can vary depending on the size, diversity, and composition of the reference panel. It is therefore essential that this is evaluated for different reference panels.

**Supplementary Note 4 - Chromosomal Distribution of Imputation Quality**

We calculated heterozygous FPR for our 0.5× with a 5% MAF and GP99 threshold across the genome by utilising a sliding window approach. Semnan3 had the lowest overall heterozygous FPR among the 4 samples (Figures S18-21). The upper 1 percentile of windows were considered as outlier regions, the highest FPR was observed in Semnan3 and Blagotin3 on chromosomes 12 and 4, respectively (Figures S19, S21). Heterozygous FPR showed greater variation in the outlier regions detected than compared with non-reference concordance (that is, there is a smaller number of regions consistently showing high spurious heterozygotes compared to overall imputation accuracy). Acem2 and Blagotin3 had the most outlier regions (157 and 155), whereas Semnan3 had the fewest (138). Despite the higher total number of outlier regions, fewer occur in the majority of samples compared to the non-reference concordance. Only 2 outlier regions occur in the majority of the samples, located on chromosomes 6 and 17 (Table S12). These regions are not strong outliers, barely passing the upper 1 percentile (represented as black bars in Figures S18-21). Overall, this highlights that heterozygous FPR is more variable compared to non-reference concordance, with sample-specific effects, and is likely not caused due to genome architecture.

**Supplementary Note 5 - Principal Component Analysis With Imputed Genotypes**

The first 10 principal components were calculated to assess imputation accuracy; all tested samples fall in their expected continental groupings in the first two principal components (Figure S22). However, a clear positive trend between accuracy and increasing downsampled coverage for imputed genomes is observed, where the 0.25× imputed genomes show the greatest deviation from the high coverage genome. To determine whether these discrepancies were due to bias introduced by imputation or instead to missingness, we conducted a PCA using more lenient GP thresholds of 0.95 and 0.80 for the imputed genomes (Figures S22-S24). We find that lower GP thresholds decrease this deviation, particularly at a GP threshold of 0.80 (Figure S24), indicating that the deviation observed might be linked to missingness rather than imputation bias. Additionally, downsampled imputed genomes shift towards 0,0, with this being most extreme in the strictest GP threshold of 0.99 and lowest coverage 0.25× (Figure S22). This shift to 0,0 in the PCA space has previously been linked to missingness (Meisner et al. 2021). When missing data is excluded in individual-based PCAs, no bias is observed between the high coverage and the imputed genomes (figure available at 10.17605/osf.io/av4f9). However, a bias is evident in the pseudohaploid genomes, which increases with decreasing coverage.

At lower coverage levels for imputation (e.g. 0.25×), even the more lenient GP threshold (0.80) deviates from the high coverage placement. This trend is also observed in ancient humans, where imputed genomes ≤0.25× show deviations in the first 4 principal components (Meisner et al. 2021). The 0.25× imputed genomes still place in their expected continental clusters, but a potential deviation from the validation should be expected. Coverages ≥0.5× show greater consistency and less deviation from the high coverage genome.

**Supplementary Note 6 - Minimum SNP Number in Plink ROH Analysis**

We observed that lower numbers of sites produced an inflation in the smallest ROH bin (0-0.5Mb; Figure S32). This inflation is most noticeable for the combined datasets (due to the low number of sites; 1.1 million for all-sites and 330K for transversions), and for individual datasets under 1 million sites (Figure S32). One approach to mitigate this inflation is to increase the minimum number of SNPs required in a ROH (--homozyg-snp). However, the impact of this parameter depends on the total number of sites used to compute ROH. Our analysis indicates that a minimum threshold of 200 SNPs per ROH eliminates excessive small ROH in datasets >750K sites (Figure S32). Additionally, we see that in individual-based datasets at the 200 SNP threshold, ROH profiles more closely match the high coverage samples than the combined all-sites dataset (i.e. ROH calculated for all samples together using transitions and transversions). For subsequent ROH analysis on imputed data, we decided to calculate ROH on datasets >750K sites with a minimum of 200 SNPs per ROH, tested both on combined datasets with no missingness and individual-based ROH calculation.

**Supplementary Note 7 - Validation of Identity-By-Descent Regions**

To assess our ability to recover shared chromosomal segments, we identified Identity-by-Descent (IBD) regions between our imputed 0.5✕ test genomes and their high coverage counterparts. Sample-specific differences were detected, with Blagotin3 showing the highest IBD sharing (3252.68 cM) and Direkli1-2 the lowest (16.1 cM; Table S16). Notably, these samples are also the closest and most distant from the reference panel, respectively. These differences may result from phasing errors and inaccurate haplotype assignments, which are likely more frequent in underrepresented individuals (Browning & Browning 2011; Hanks et al. 2022; Shi et al. 2023). Further exploration of phasing errors and haplotype missassignments could improve our understanding of processes that introduce artificial breakpoints, leading to fragmentation of IBD segments.


Browning SR, Browning BL. 2011. Haplotype phasing: existing methods and new developments. Nat. Rev. Genet. 12:703–714. doi: 10.1038/nrg3054.

Erven JAM et al. 2024. A high-coverage Mesolithic aurochs genome and effective leveraging of ancient cattle genomes using whole genome imputation. Mol. Biol. Evol. 41. doi: 10.1093/molbev/msae076.

Hanks SC et al. 2022. Extent to which array genotyping and imputation with large reference panels approximate deep whole-genome sequencing. Am. J. Hum. Genet. 109:1653–1666. doi: 10.1016/j.ajhg.2022.07.012.

Marchini J, Howie B. 2010. Genotype imputation for genome-wide association studies. Nat. Rev. Genet. 11:499–511. doi: 10.1038/nrg2796.

Meisner J, Liu S, Huang M, Albrechtsen A. 2021. Large-scale inference of population structure in presence of missingness using PCA. Bioinformatics. 37:1868–1875. doi: 10.1093/bioinformatics/btab027.

Shi M et al. 2023. Genotype imputation accuracy and the quality metrics of the minor ancestry in multi-ancestry reference panels. bioRxiv. doi: 10.1101/2023.05.30.542466.

### Supplementary Figures

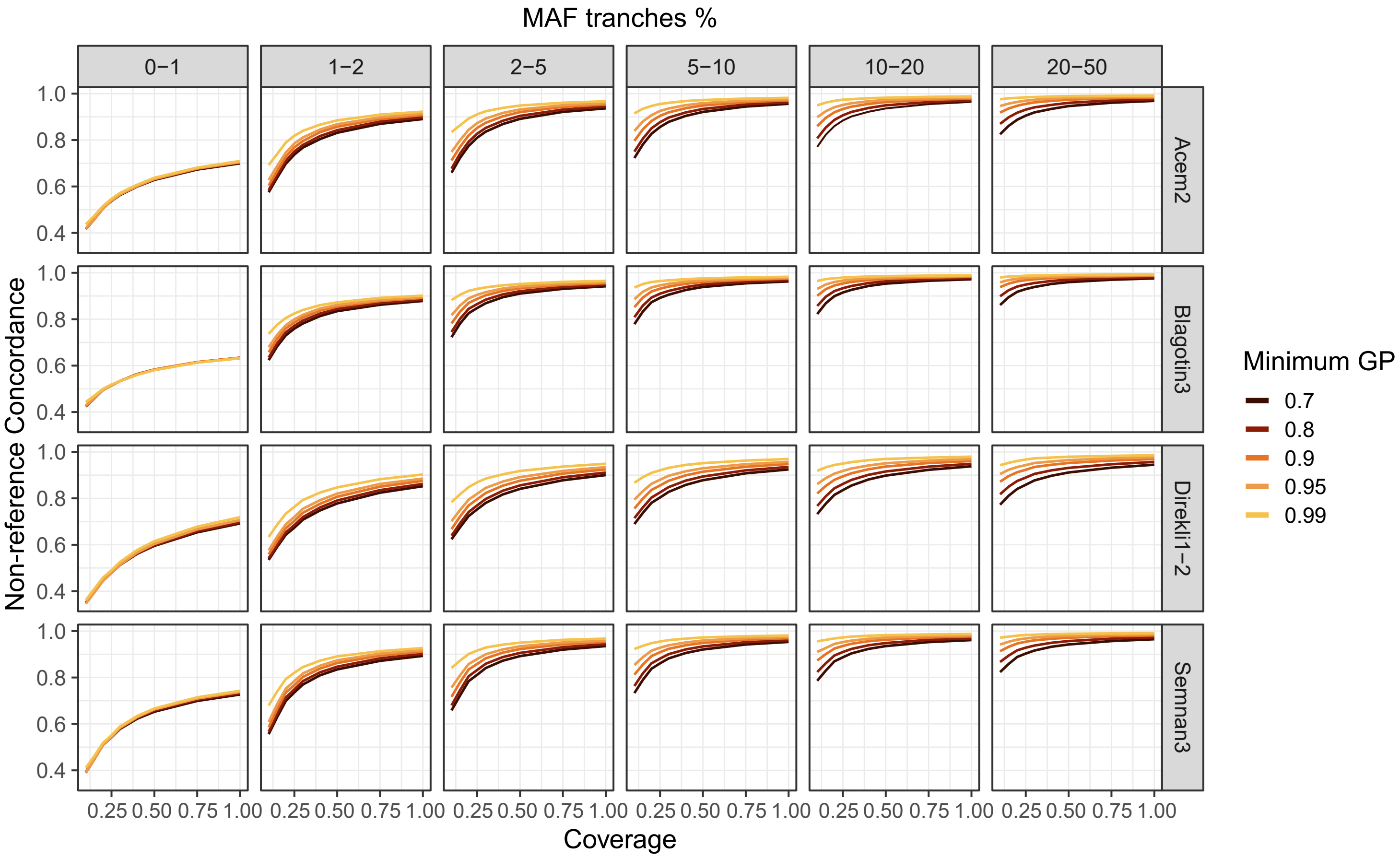


**Figure S1: Non-reference concordance across MAF tranches for four samples, with downsampled coverages ranging from 0.1-1X.** Concordance values are calculated for transitions and transversions combined. Colors indicate different genotype probability (GP) thresholds.

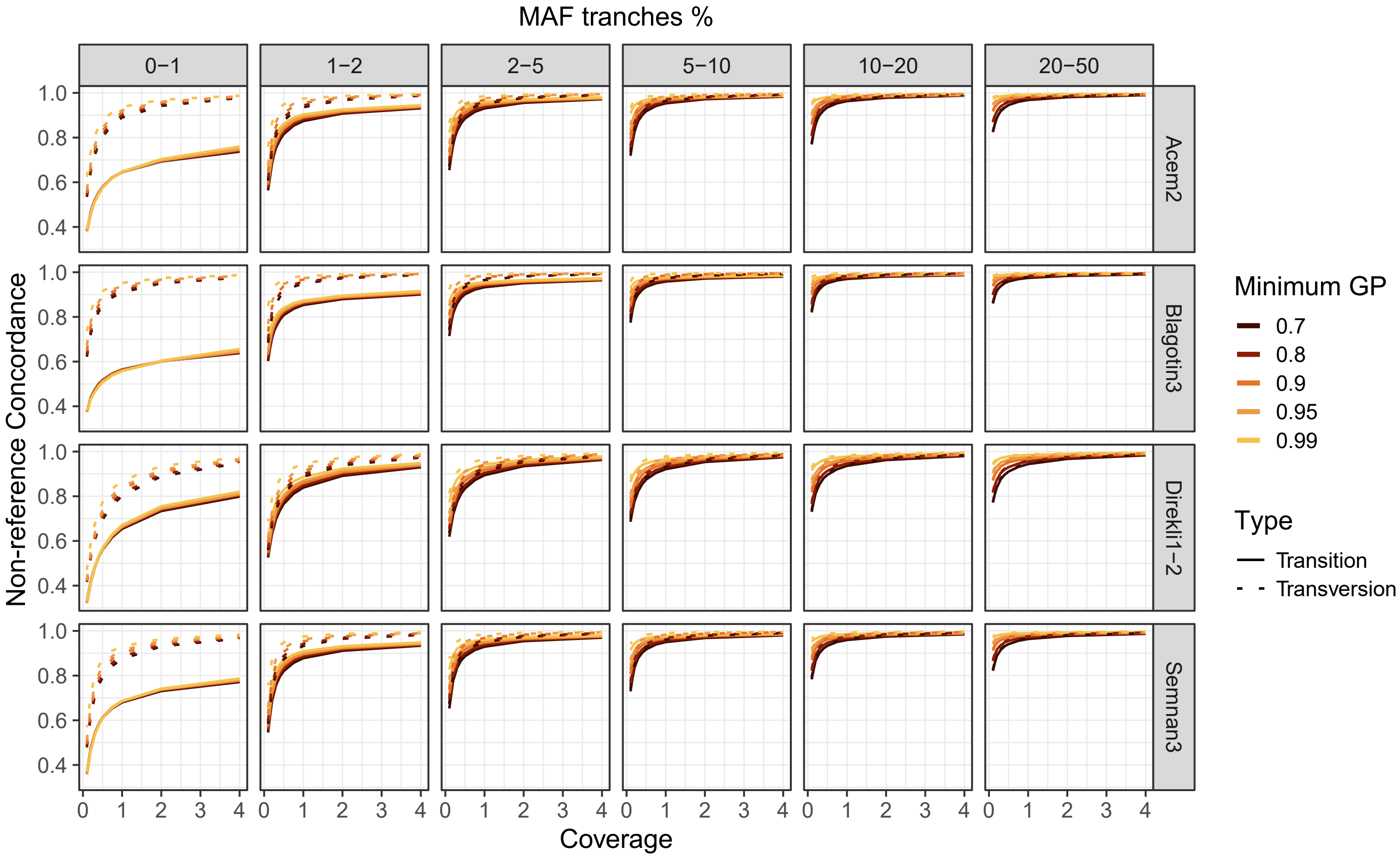


**Figure S2: Non-reference concordance across MAF tranches for four samples, with downsampled coverages ranging from 0.1-4X.** Concordance values are calculated for transitions and transversions separately, depicted by Type. Colors indicate different GP thresholds.

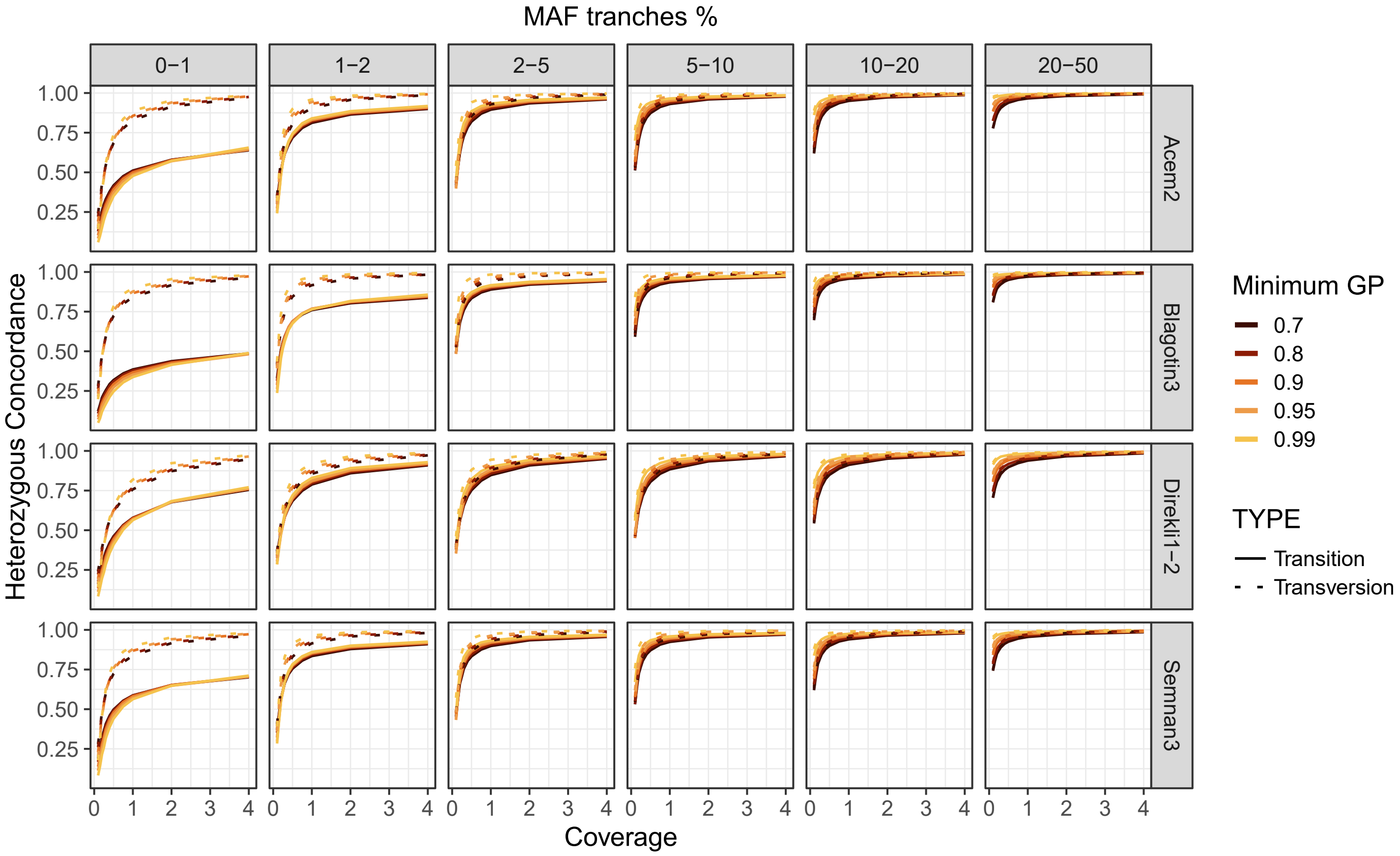


**Figure S3: Heterozygous concordance across MAF tranches for four samples, with downsampled coverages ranging from 0.1-4X.** Concordance values are calculated for transitions and transversions seperately, depicted by type. Colors indicate different genotype probability (GP) thresholds.

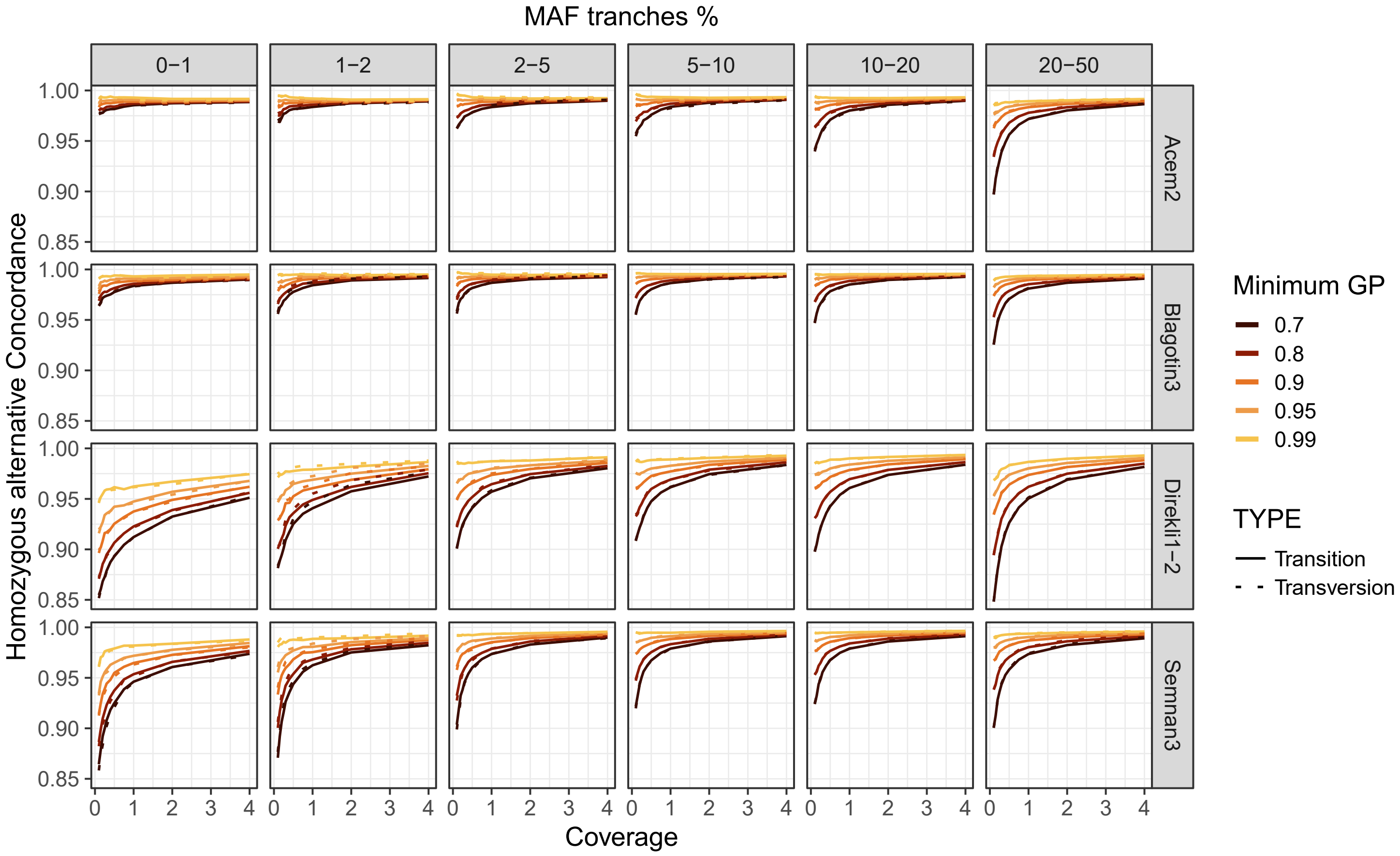


**Figure S4: Homozygous alternative concordance across MAF tranches for four samples, with downsampled coverages ranging from 0.1-4X.** Concordance values are calculated for transitions and transversions seperately, depicted by type. Colors indicate different genotype probability (GP) thresholds.

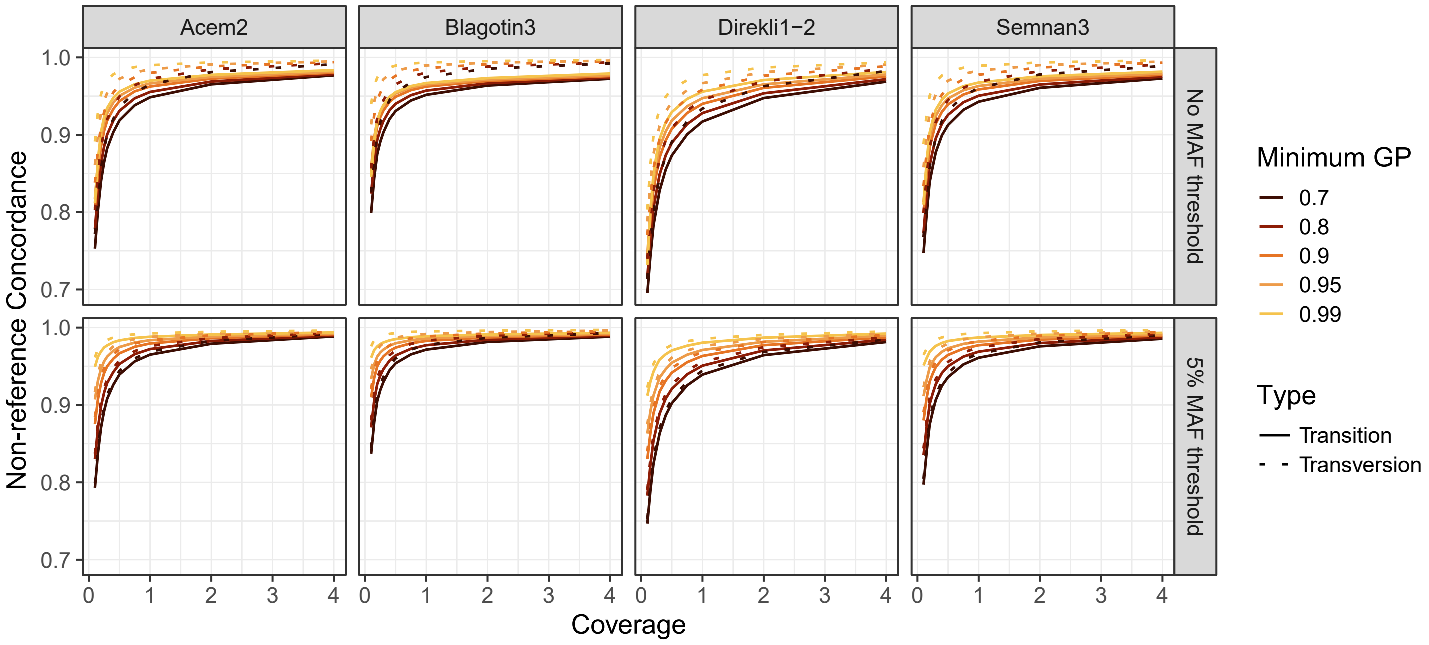


**Figure S5: Non-reference concordance across MAF thresholds for four samples, with downsampled coverages ranging from 0.1-4X.** Concordance values are calculated for transitions and transversions separately, depicted by Type. Colors indicate different GP thresholds.

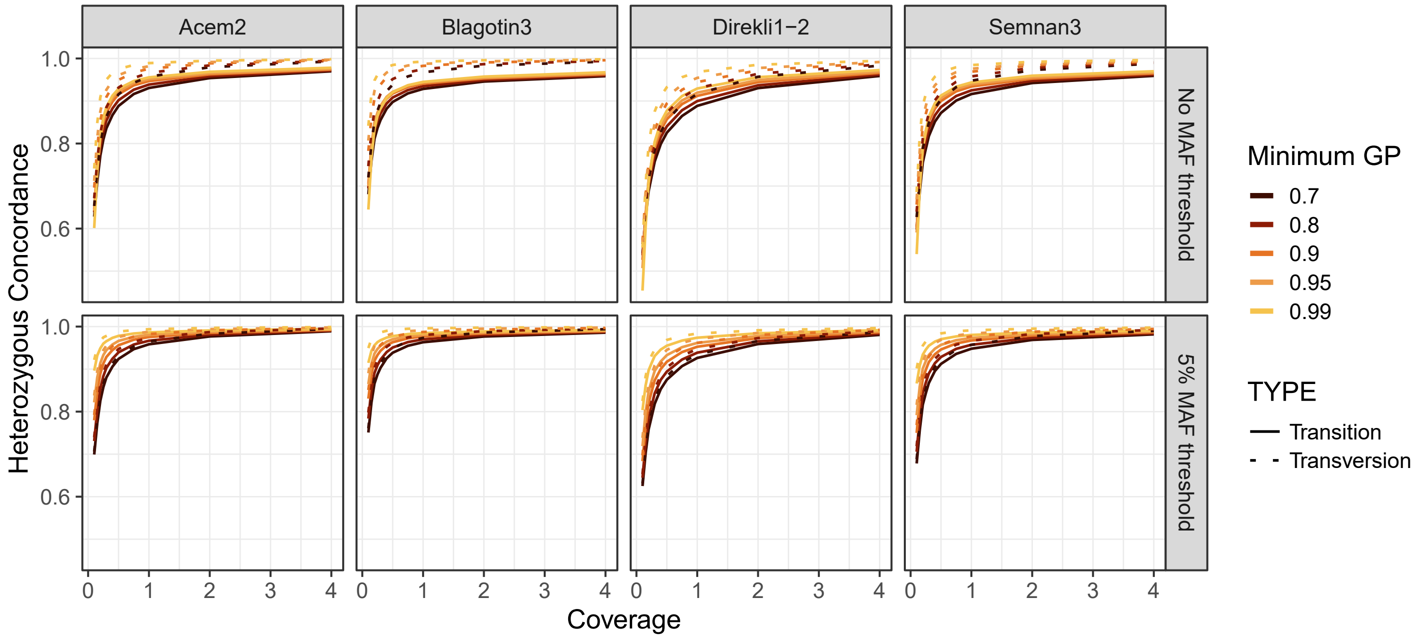


**Figure S6: Heterozygous concordance across MAF thresholds for four samples, with downsampled coverages ranging from 0.1-4X.** Concordance values are calculated for transitions and transversions separately, depicted by Type. Colors indicate different GP thresholds.

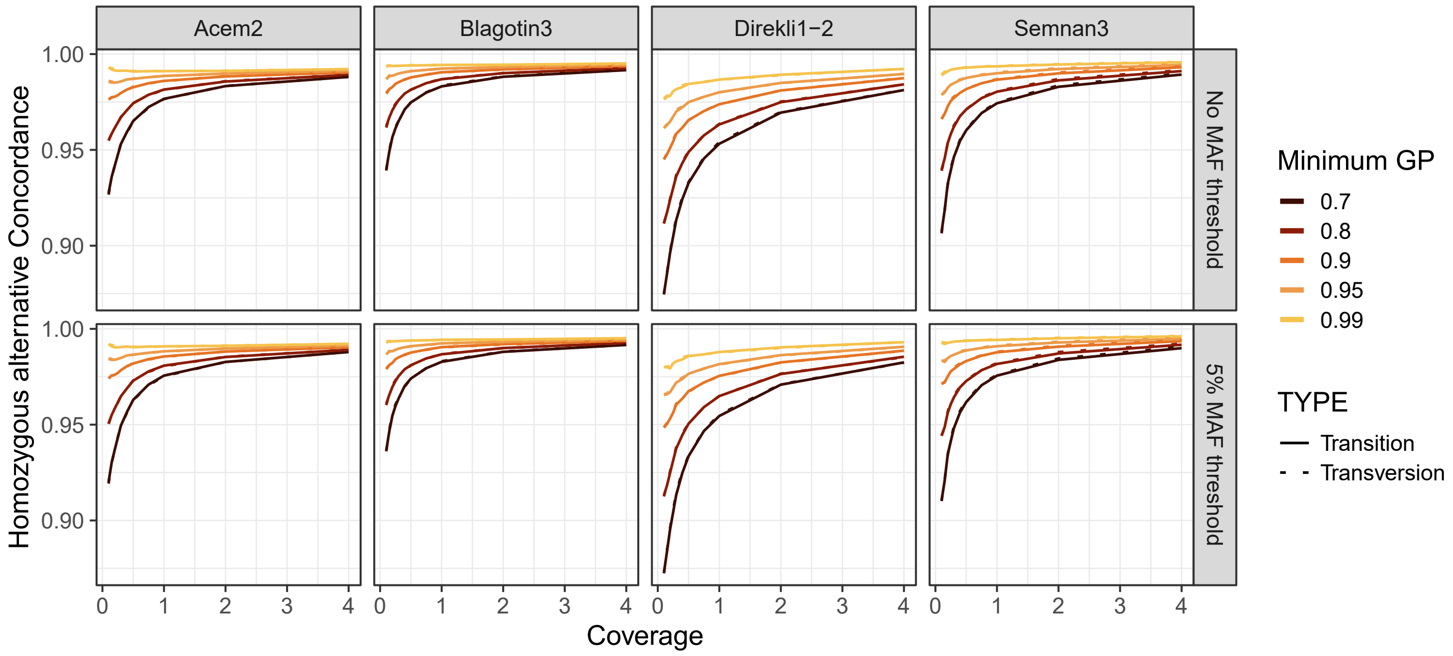


**Figure S7: Homozygous alternative concordance across MAF thresholds for four samples, with downsampled coverages ranging from 0.1-4X.** Concordance values are calculated for transitions and transversions separately, depicted by Type. Colors indicate different GP thresholds.

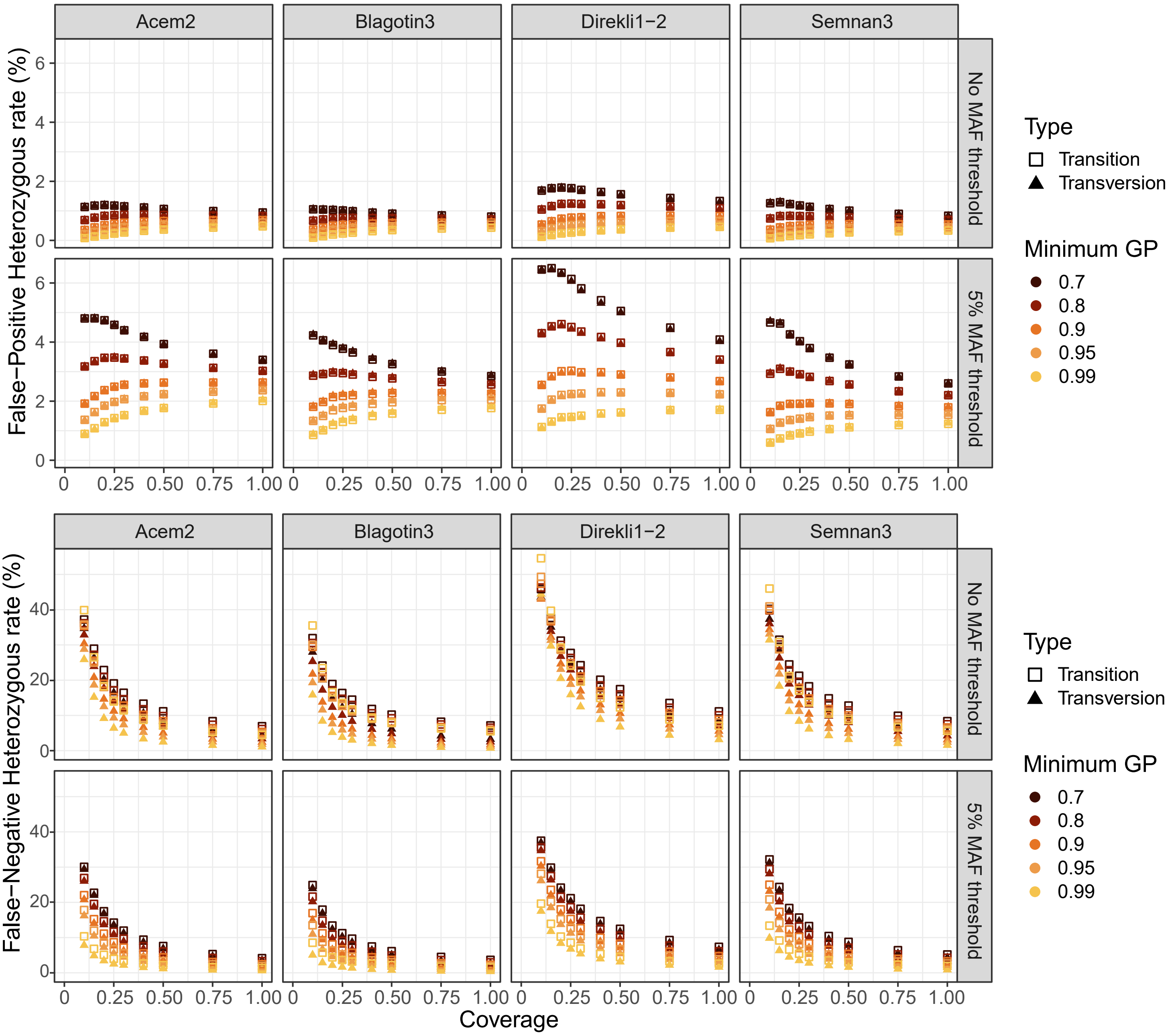


**Figure S8: Patterns of imputed heterozygous veracity for different MAF thresholds. A) false-positive rate (FPR) and B) false-negative rate (FNR).** "True" heterozygotes are defined by the validation non-imputed genotypes calculated using the entirety of sequencing data available for each test sample.

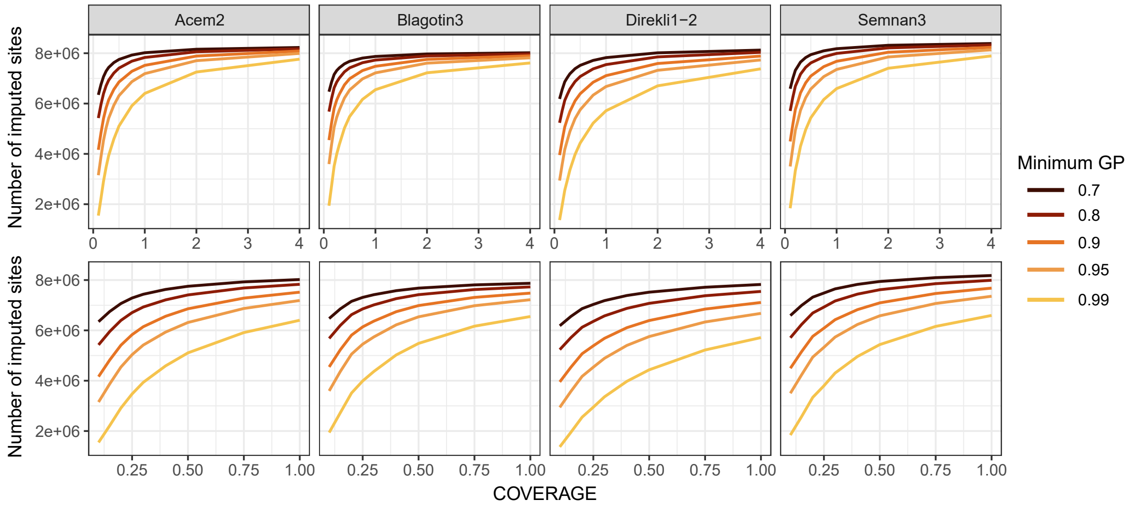


**Figure S9: Recovery of the total number of imputed sites at MAF 5% for four samples, with downsampled coverages ranging from 0.1-4X (upper) and 0.1-1X (lower).** Recovery is calculated for transitions and transversions combined. Colors indicate different GP thresholds.

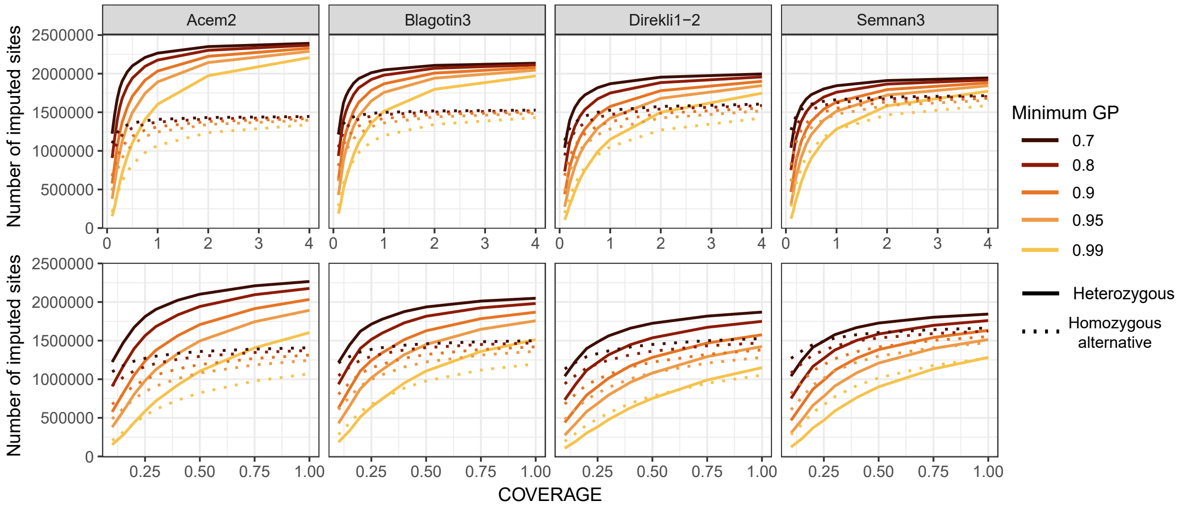


**Figure S10: Recovery of the total number of imputed sites at MAF 5% for four samples, with downsampled coverages ranging from 0.1-4X (upper) and 0.1-1X (lower).** Recovery is calculated for transitions and transversions combined. Heterozygous and Homozygous alternative genotypes are sepera, depicted by type. Colors indicate different GP thresholds.

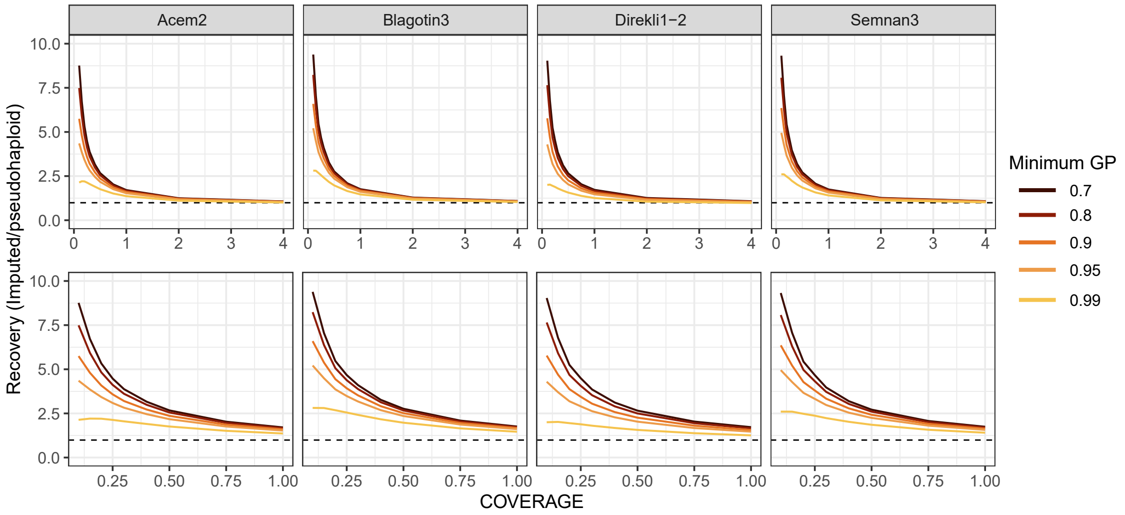


**Figure S11: Recovery of the total number of imputed sites at MAF 5% against their pseudohaploid calls for four samples, with downsampled coverages ranging from 0.1-4X (upper) and 0.1-1X (lower).** Recovery is calculated for transitions and transversions combined. Colors indicate different GP thresholds. The black dashed line indicates 1 i.e. no difference between the number of imputed and pseudohaploid calls. Values greater than 1 indicated an increase in the number of variant sites in the imputed data relative to the non-imputed data; values less than 1 a decrease.

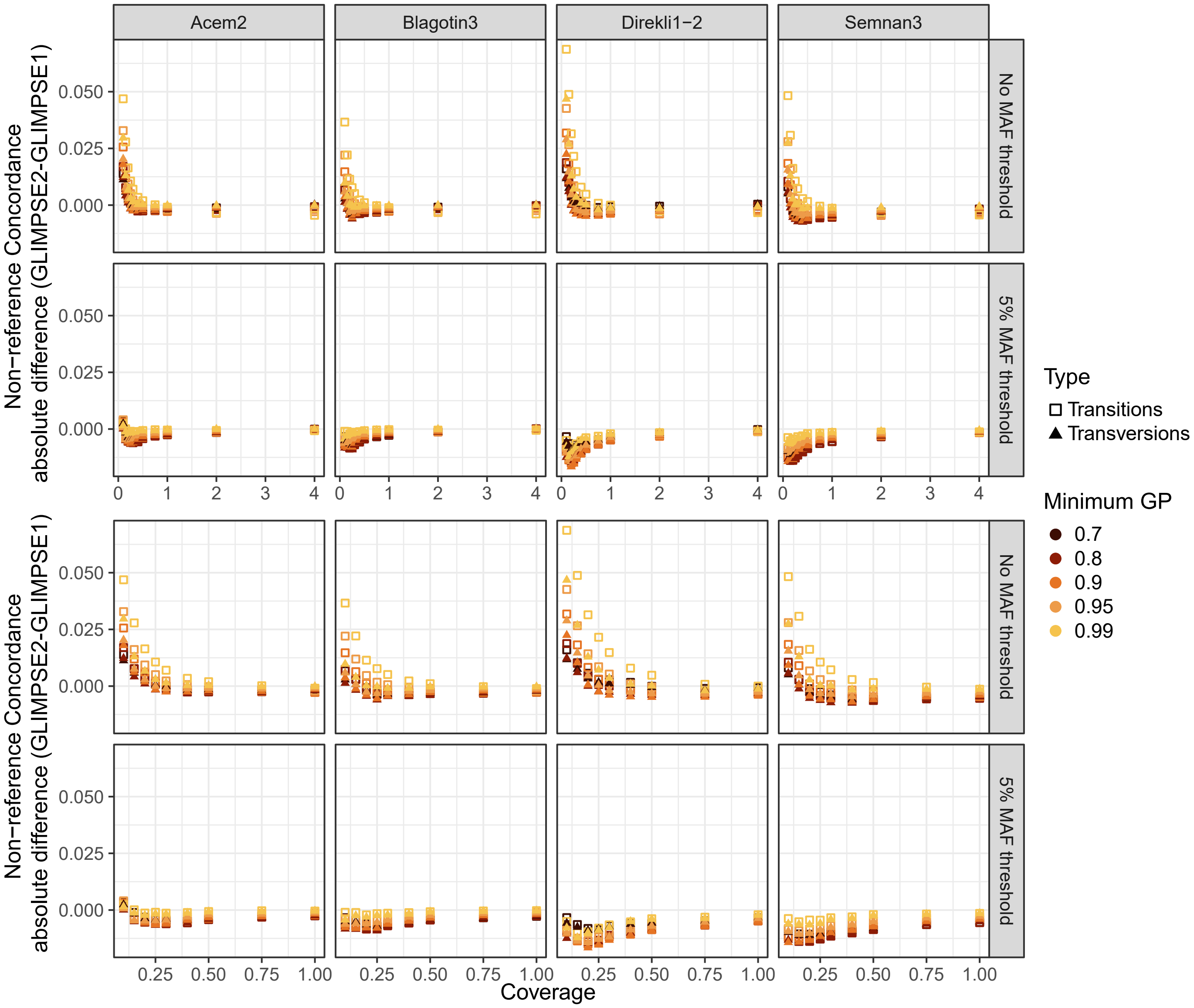


**Figure S12: Absolute difference between GLIMSPE2 and GLIMPSE1 non-reference concordance rates.** Positive indicates a higher GLIMSPE2 concordance, negative indicates a higher GLIMPSE1 GP filter concordance.

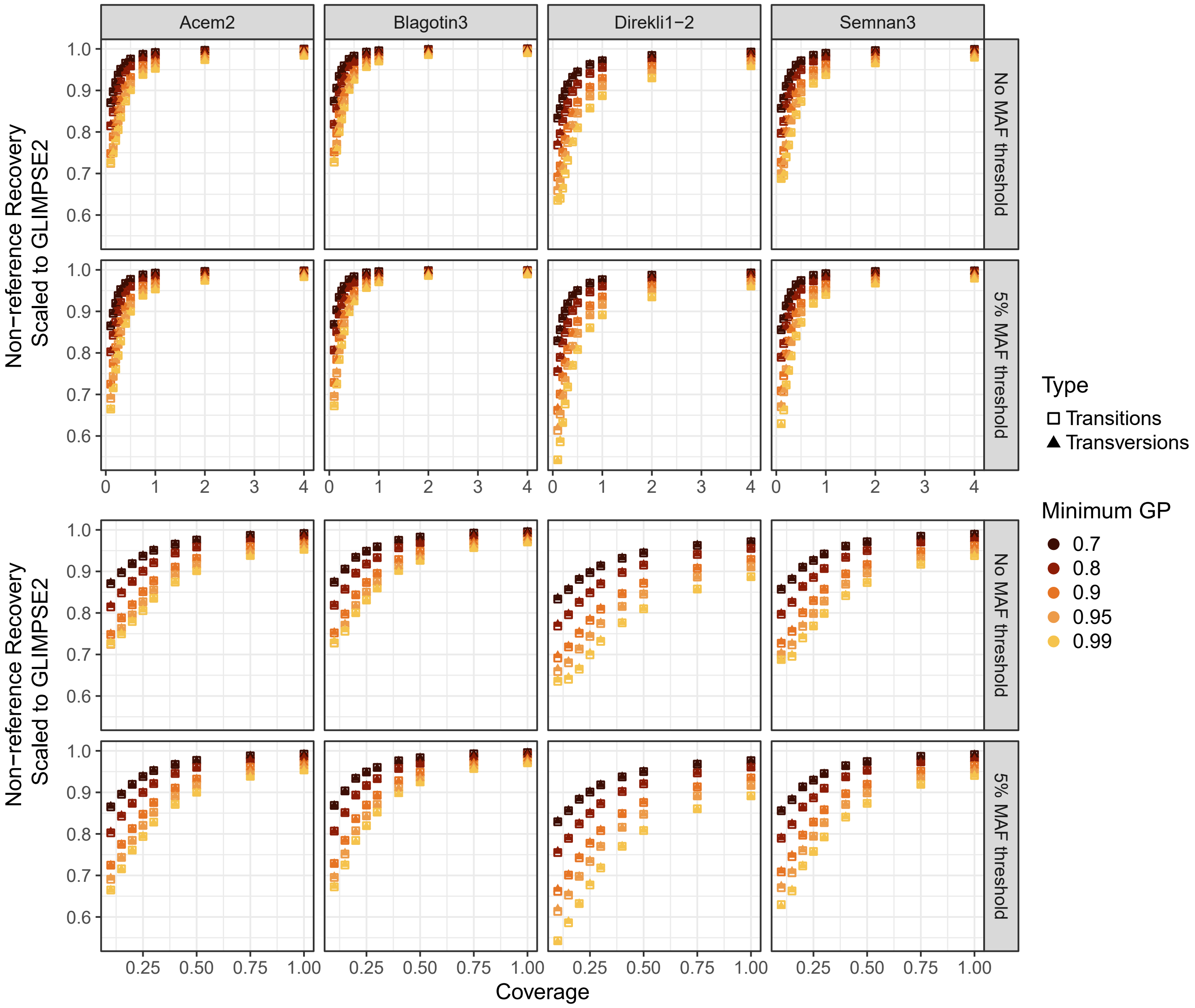


**Figure S13: Recovery of non-reference genotypes using GLIMPSE1 scaled to recovery using GLIMPSE2.** 1 indicates identical site number recovery; below 1 indicates a higher recovery of GLIMPSE2.

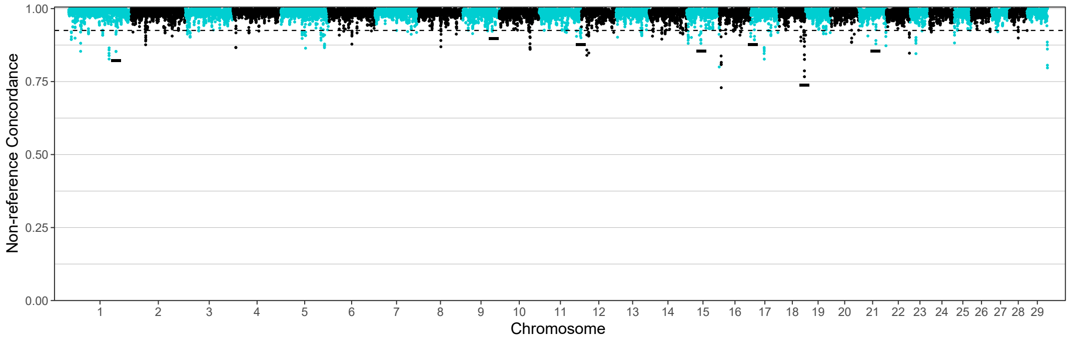


**Figure S14: Non-reference concordance across chromosomes for Acem2, divided in sliding windows.** The dotted line equals the 1 percentile. Each dot represents a 100kb step-size window. Black bars denote reduced accuracy regions shared across the majority of the test samples.

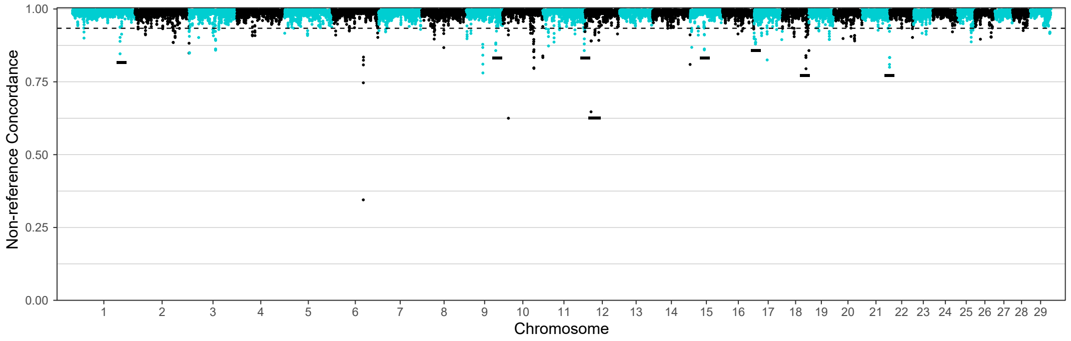


**Figure S15: Non-reference concordance across chromosomes for Blagotin3, divided in sliding windows.** The dotted line equals the 1 percentile. Each dot represents a 100kb step-size window. Black bars denote reduced accuracy regions shared across the majority of the test samples.

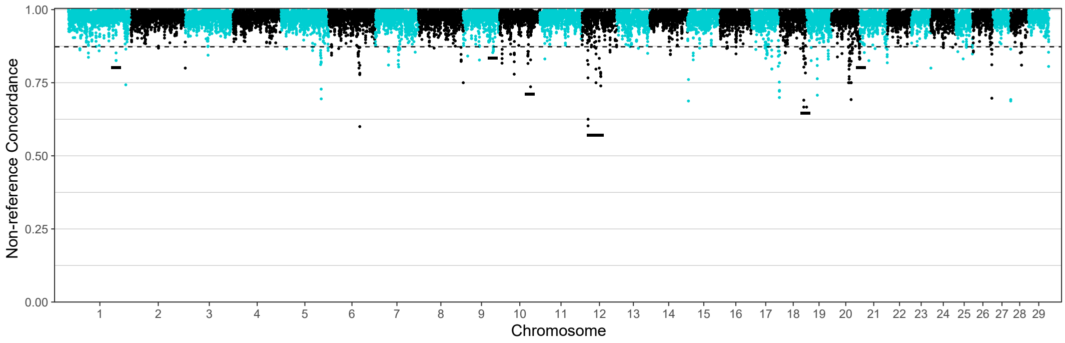


**Figure S16: Non-reference concordance across chromosomes for Direkli1-2, divided in sliding windows.** The dotted line equals the 1 percentile. Each dot represents a 100kb step-size window. Black bars denote reduced accuracy regions shared across the majority of the test samples.

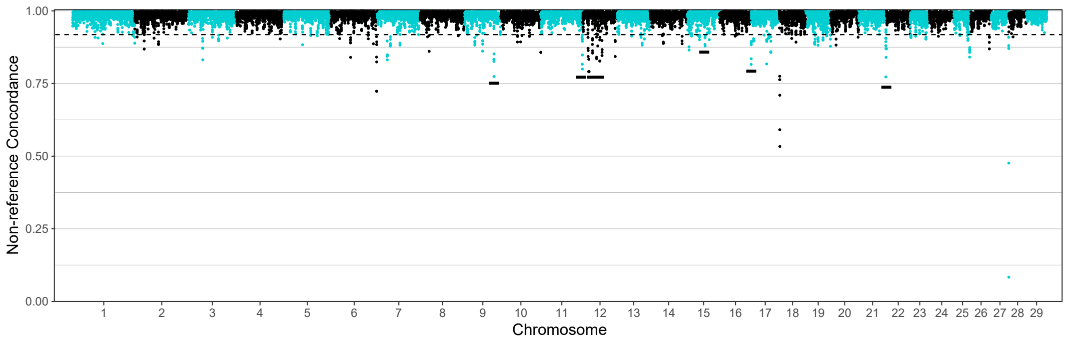


**Figure S17: Non-reference concordance across chromosomes for Semnan3, divided in sliding windows.** The dotted line equals the 1 percentile. Each dot represents a 100kb step-size window. Black bars denote reduced accuracy regions shared across the majority of the test samples.

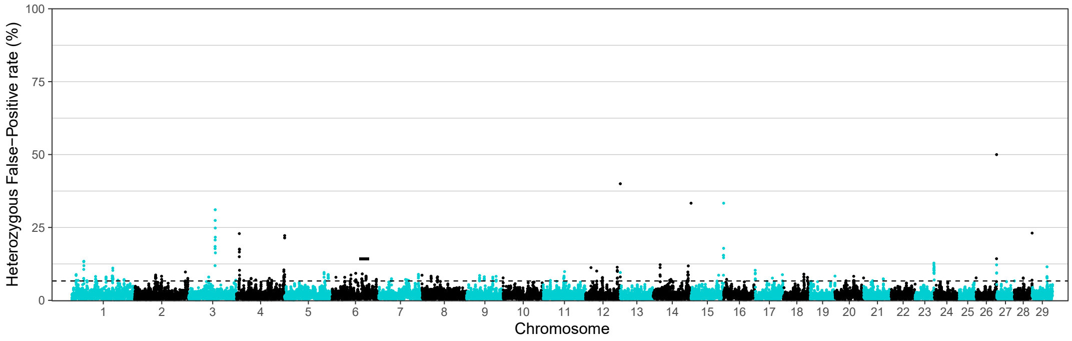


**Figure S18: Heterozygous false-positive rate (FPR in %) across chromosomes for Acem2, divided in sliding windows.** The dotted line equals the 1 percentile. Each dot represents a 100kb step-size window. Black bars denote reduced accuracy regions shared across the majority of the test samples.

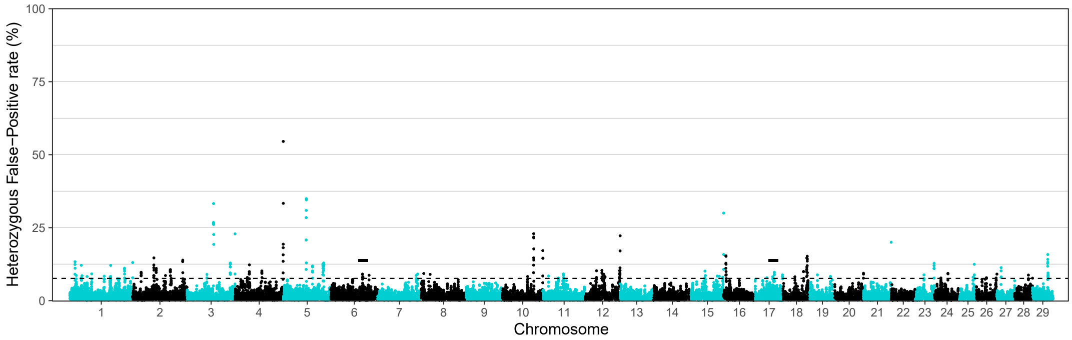


**Figure S19: Heterozygous false-positive rate (FPR in %) across chromosomes for Blagotin3, divided in sliding windows.** The dotted line equals the 1 percentile. Each dot represents a 100kb step-size window. Black bars denote reduced accuracy regions shared across the majority of the test samples.

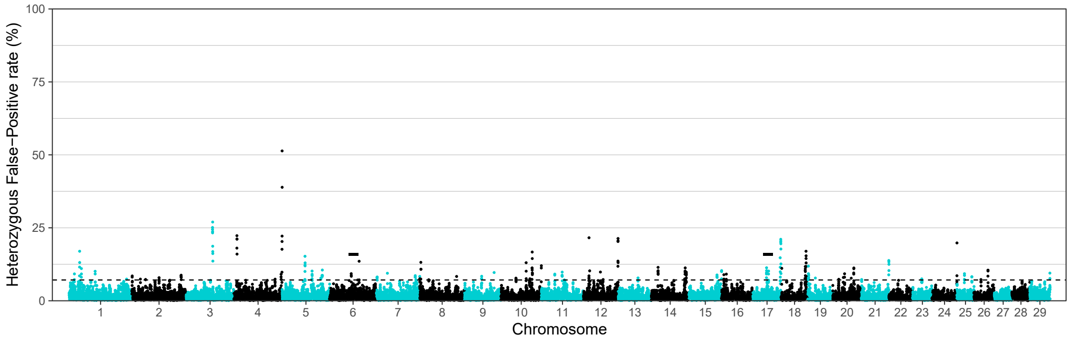


**Figure S20: Heterozygous false-positive rate (FPR in %) across chromosomes for Direkli1-2, divided in sliding windows.** The dotted line equals the 1 percentile. Each dot represents a 100kb step-size window. Black bars denote reduced accuracy regions shared across the majority of the test samples.

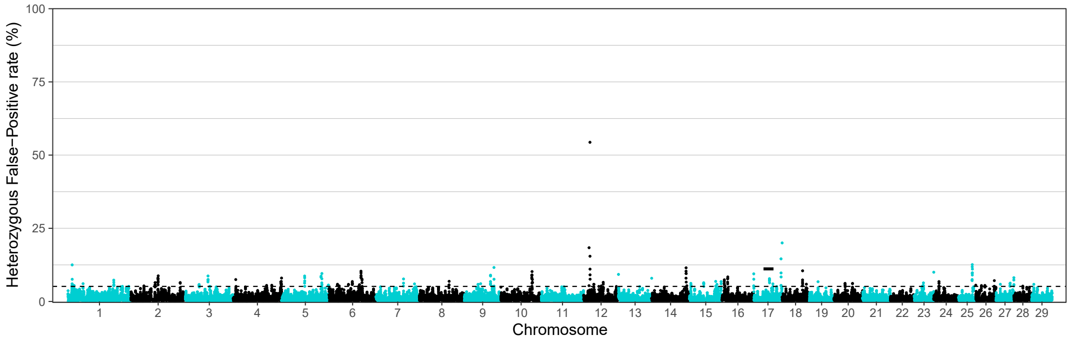


**Figure S21: Heterozygous false-positive rate (FPR in %) across chromosomes for Semnan3, divided in sliding windows.** The dotted line equals the 1 percentile. Each dot represents a 100kb step-size window. Black bars denote reduced accuracy regions shared across the majority of the test samples.

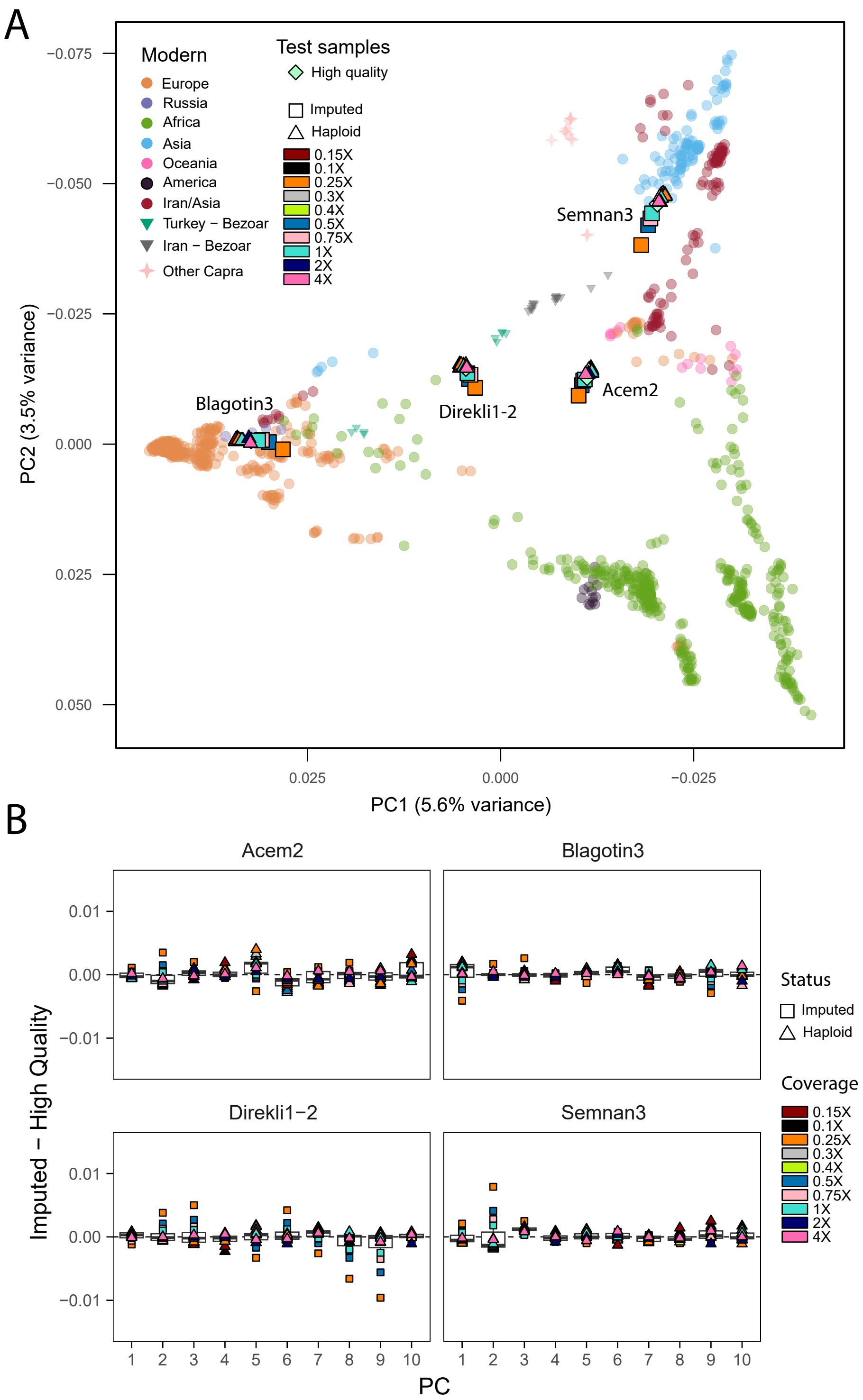


**Figure S22: Principal Components Analysis (PCA) of imputed, pseudohaploid and high coverage genotypes of the four test individuals onto the modern Vargoats reference Panel, imputed genotypes are filtered for GP99.** A) Imputed, pseudohaploid and high quality genotypes of the test samples were projected onto the modern reference panel along the first two eigenvectors. Dataset was filtered for a MAF ≥5%. Key indicates geographical region and ancestry for modern individuals, key indicates coverage and sampling strategy for test samples. B) Boxplots of the normalised differences in the coordinates of the imputed, pseudohaploid and high coverage genomes for the first ten principle components. The hori-zontal lines of the box plot represent the first quartile, median and third quartile, whiskers represent 1.5 times the quartile range.

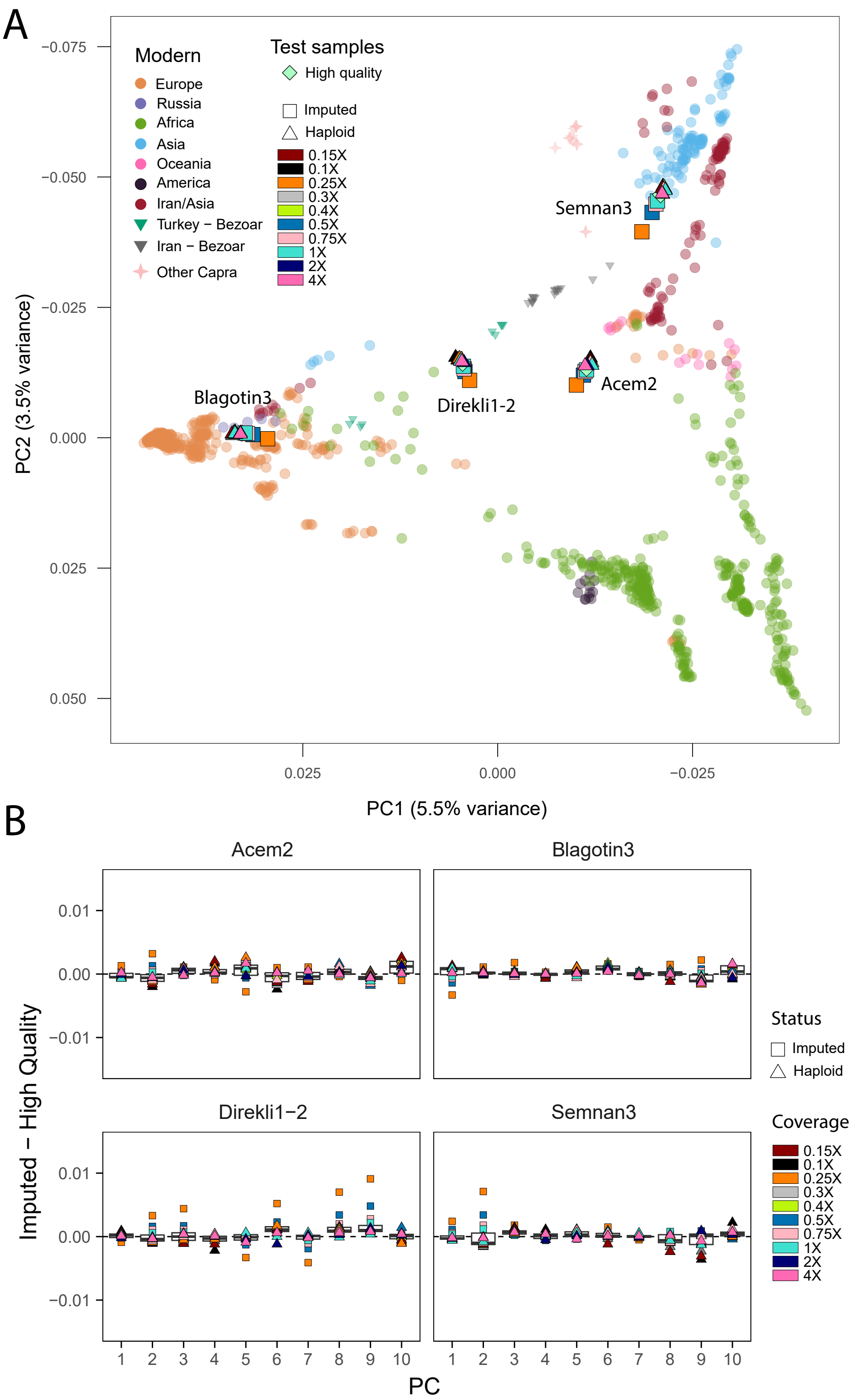


**Figure S23: Principal Components Analysis (PCA) of imputed, pseudohaploid and high coverage genotypes of the four test individuals onto the modern Vargoats reference Panel, imputed genotypes are filtered for GP95.** A) Imputed, pseudohaploid and high quality genotypes of the test samples were projected onto the modern reference panel along the first two eigenvectors. Dataset was filtered for a MAF ≥5%. Key indicates geographical region and ancestry for modern individuals, key indicates coverage and sampling strategy for test samples. B) Boxplots of the normalised differences in the coordinates of the imputed, pseudohaploid and high coverage genomes for the first ten principle components. The hori-zontal lines of the box plot represent the first quartile, median and third quartile, whiskers represent 1.5 times the quartile range.

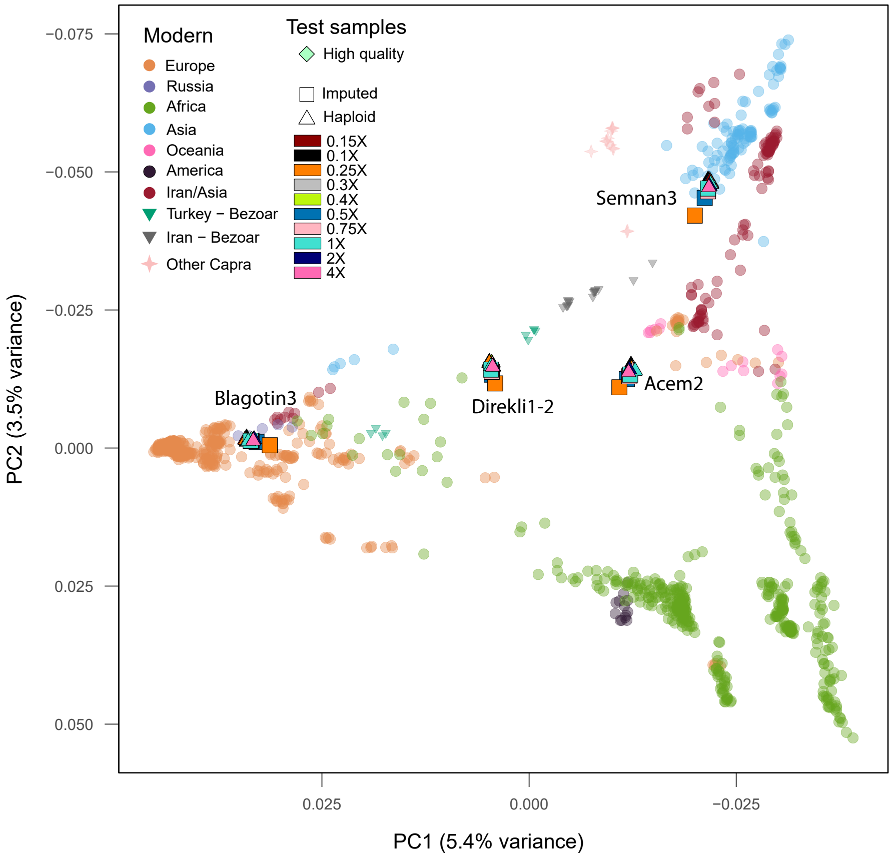


**Figure S24: PCA of imputed, pseudohaploid and high coverage genotypes of the four test individuals onto the modern Vargoats reference Panel, imputed genotypes are filtered for GP80.** Imputed, pseudohaploid and high quality genotypes of the test samples were projected onto the modern reference panel along the first two eigenvectors. The dataset was filtered for a MAF ≥5%. Key indicates geographical region and ancestry for modern individuals, key indicates coverage and sampling strategy for test samples.

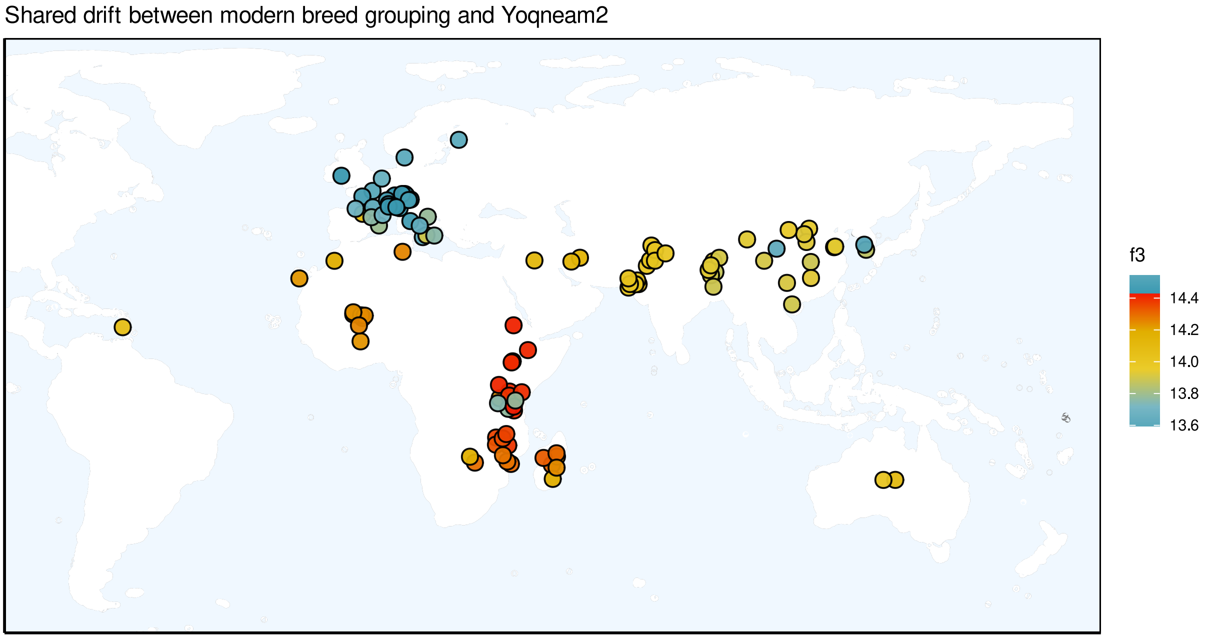


**Figure S25:** Geographical representation of outgroup f3 values of a Bronze Age goat from Tel Yoqne'am Israel (Yoqneam2) and modern domestic goat breeds, measuring the relative shared drift between Yoqneam2 and modern breeds.

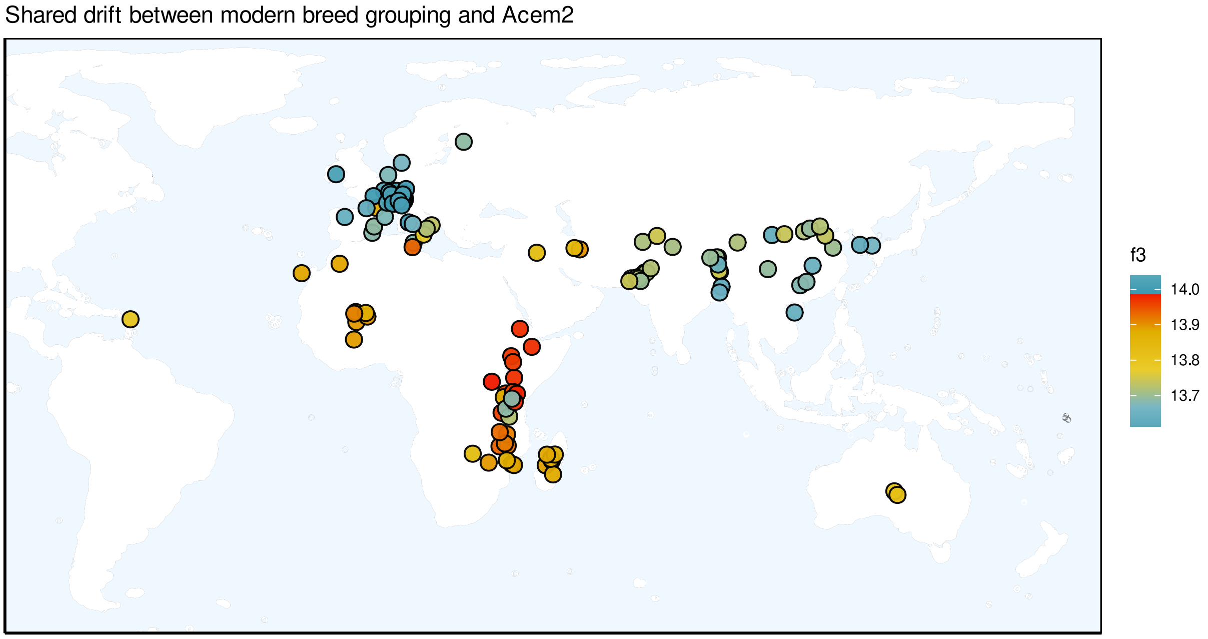


**Figure S26:** Geographical representation of outgroup f3 values of a Bronze Age goat from Acemhöyuk, Central Türkiye (Acem2) and modern domestic goat breeds, measuring the relative shared drift between Acem2 and modern breeds.

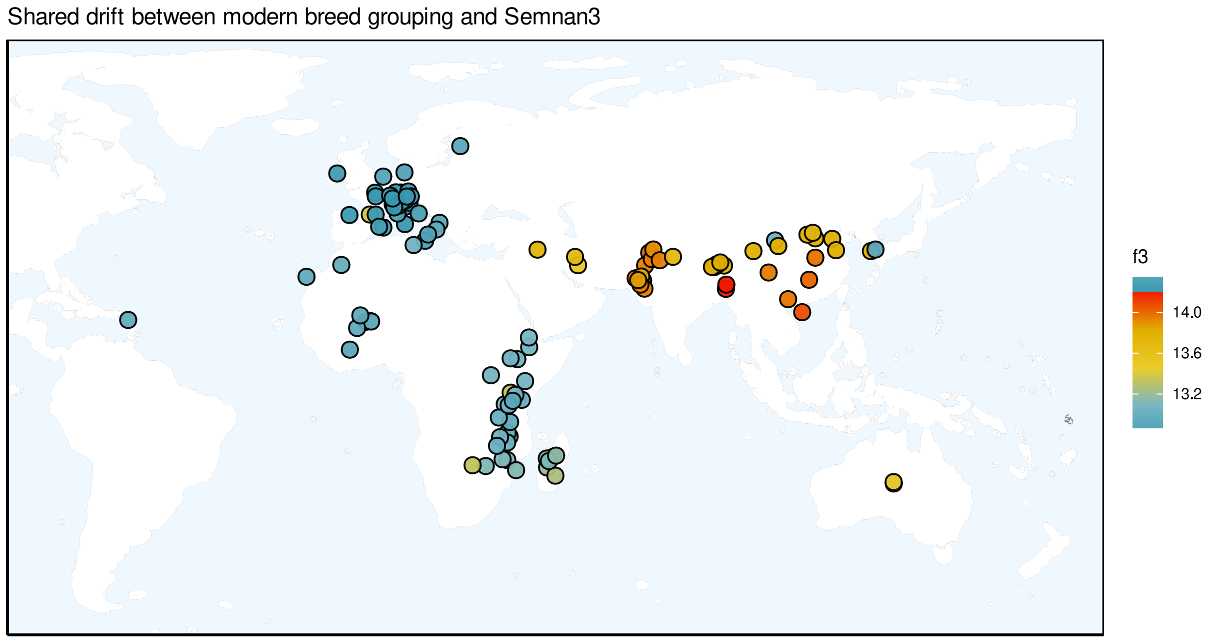


**Figure S27:** Geographical representation of outgroup f3 values of a Neolithic goat from Sang-e Chakhmaq, Iran (Semnan3) and modern domestic goat breeds, measuring the relative shared drift between Semnan3 and modern breeds.

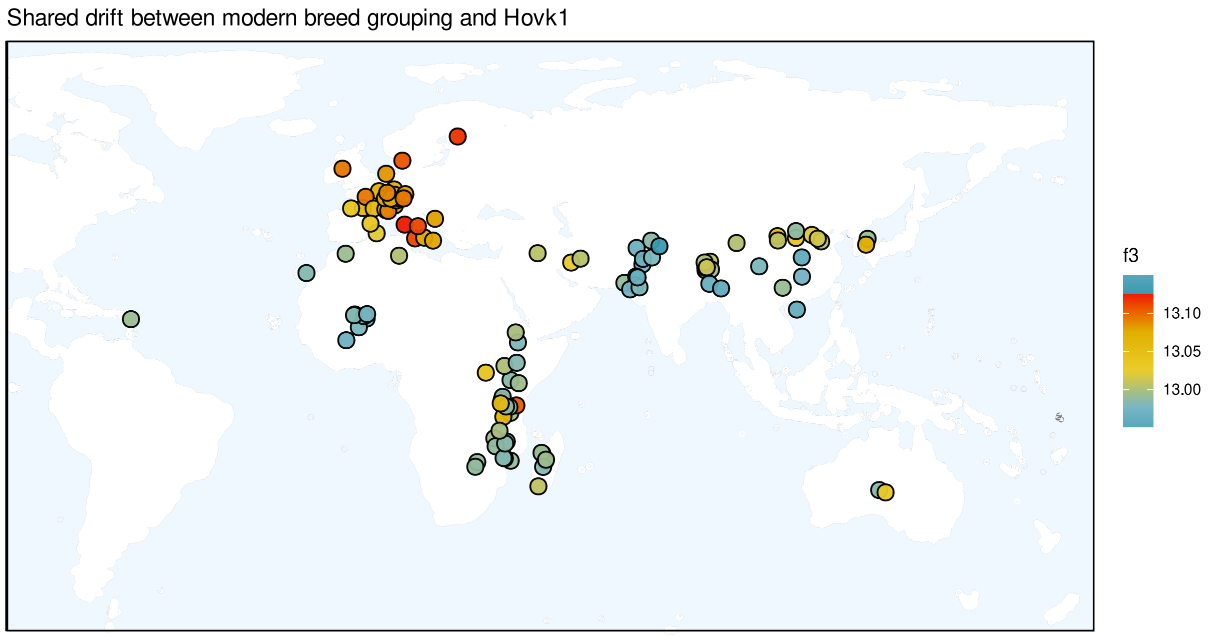


**Figure S28:** Geographical representation of outgroup f3 values of a Late Pleistocene *Capra* from Hovk-1 Cave, Armenia (Hovk1) and modern domestic goat breeds, measuring the relative shared drift between Hovk1 and modern breeds.

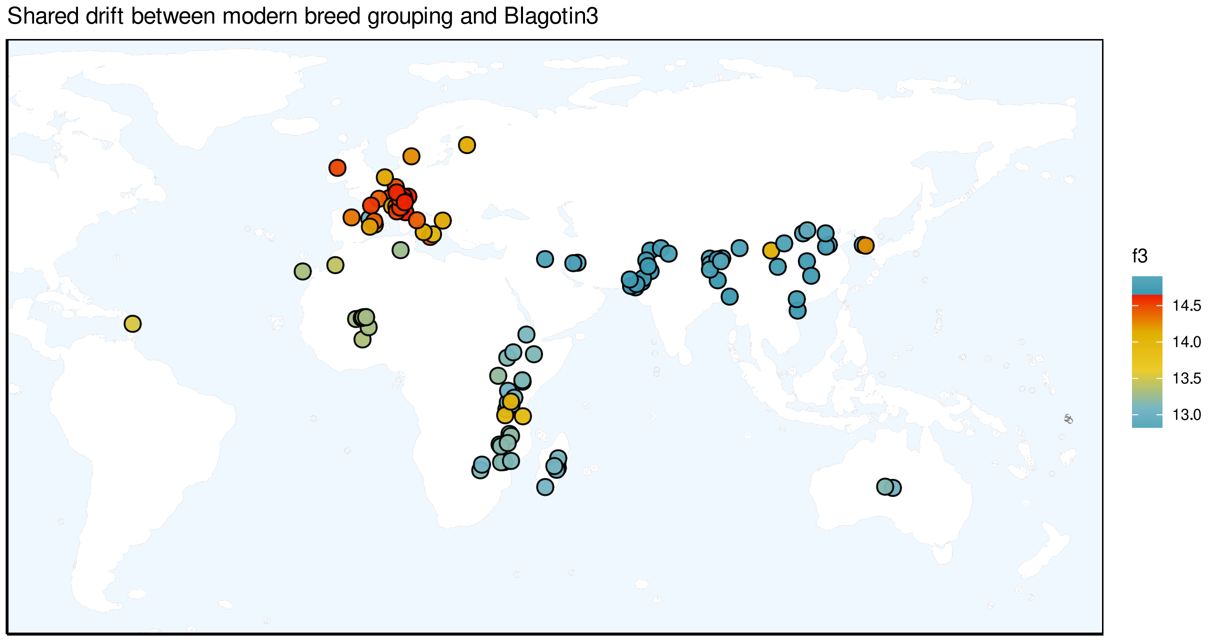


**Figure S29:** Geographical representation of outgroup f3 values of a Neolithic goat from Blagotin-Poljna, Serbia (Blagotin3) and modern domestic goat breeds, measuring the relative shared drift between Blagotin3 and modern breeds.

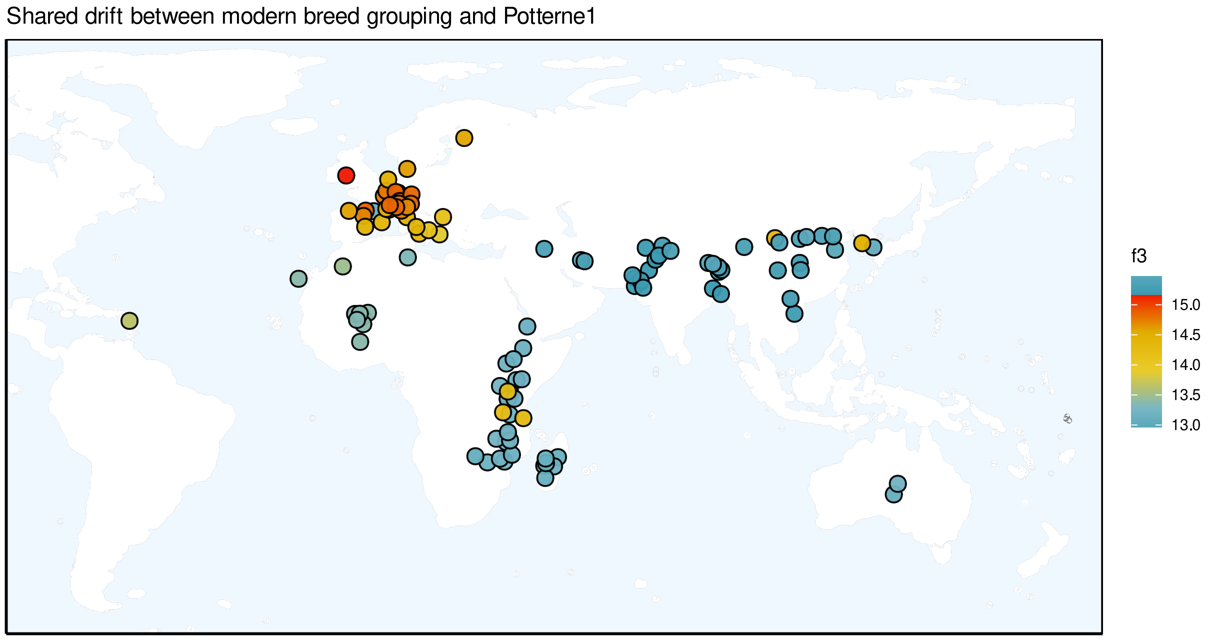


**Figure S30:** Geographical representation of outgroup f3 values of a Bronze Age goat from Potterne, UK (Potterne1) and modern domestic goat breeds, measuring the relative shared drift between Potterne1 and modern breeds.

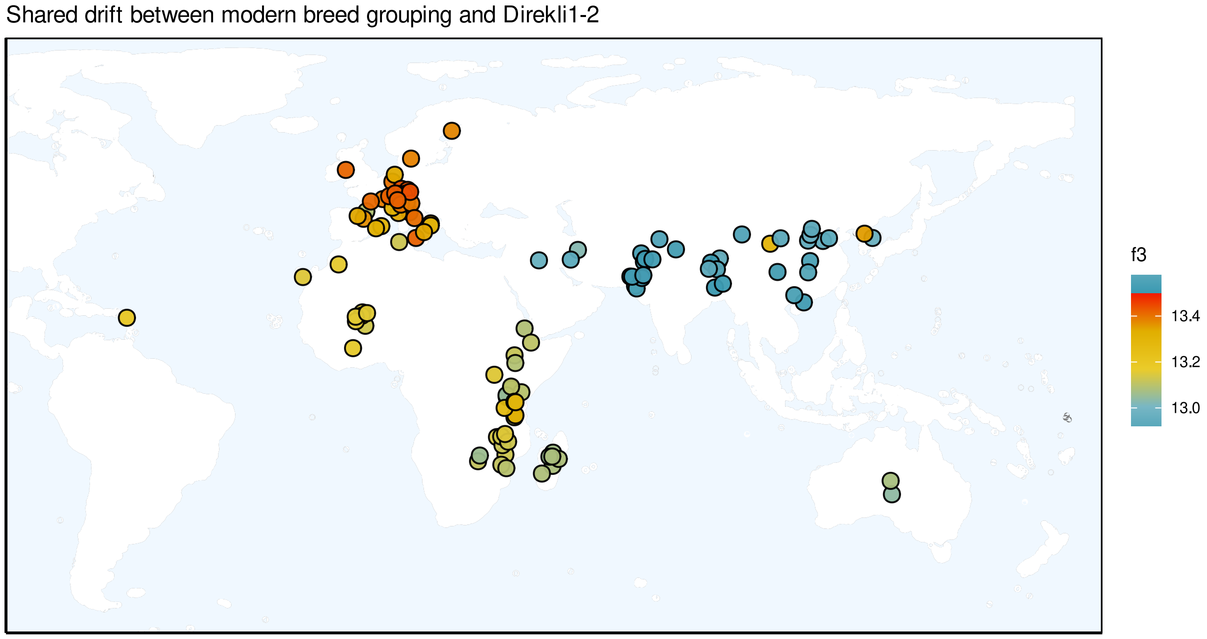


**Figure S31:** Geographical representation of outgroup f3 values of a Late Epipaleolithic bezoar from the Direkli Cave, Taurus Mountains of Türkiye (Direkli1-2) and modern domestic goat breeds, measuring the relative shared drift between Direkli1-2 and modern breeds.

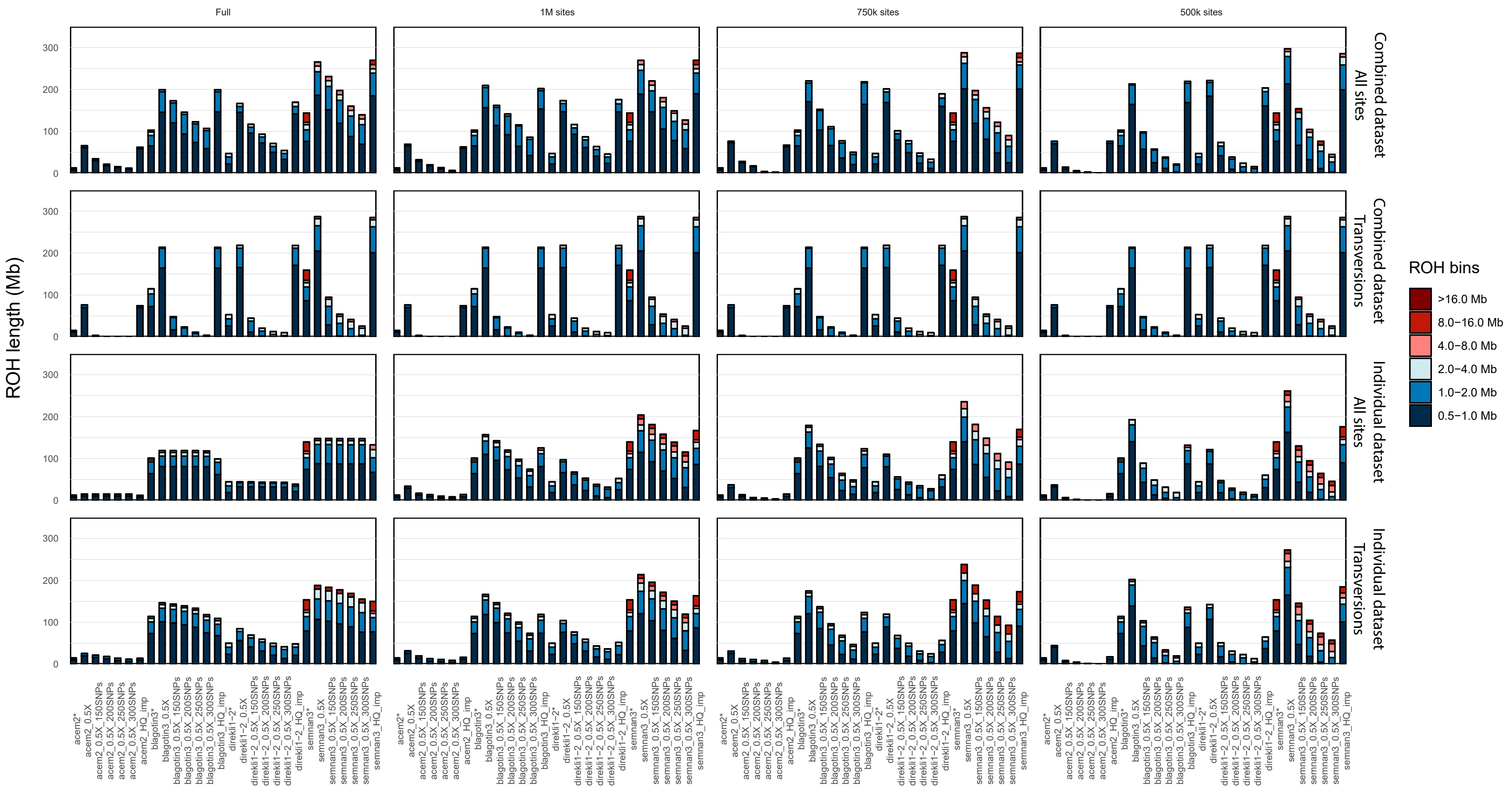


**Figure S32: Runs of Homozygosity (ROH) analysis across 0.5X and high coverage imputed genomes using different datasets, downsampling strategies, and SNP configurations.** ROH profiles are categorized by length bins, represented in color. All datasets were filtered with a MAF 5% and GP99 threshold. The columns represent the downsampling tests, and the rows correspond to the different datasets used. The SNP thresholds (--homozyg-SNP, only performed on the 0.5X imputed genomes) are indicated on the x-axis. *For non-imputed high coverage samples, ROH was computed individually, filtered for all-sites and transversions only, without downsampling.

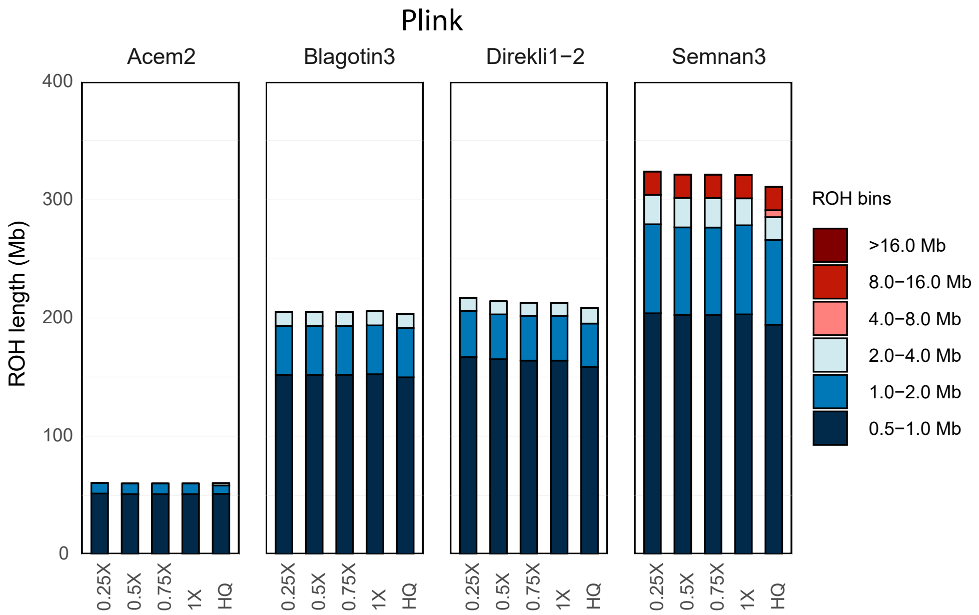


**Figure S33: ROH length bin profiles from Plink for 0.25X, 0.5X, 0.75X, 1X imputed and high coverage ancient goats, stratified by sample.** Samples were tested individually, with no missingness allowed in the dataset.

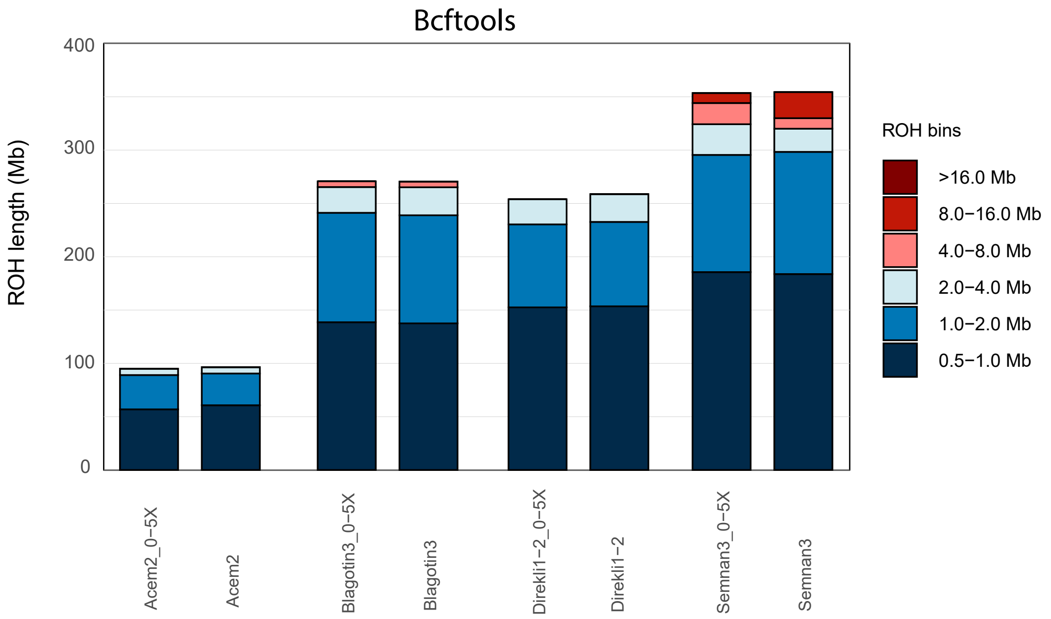


**Figure S34: ROH length bin profiles for 0.5X imputed and high coverage ancient goats, stratified by sample calculated with bcftools.** No missingness allowed in the dataset.

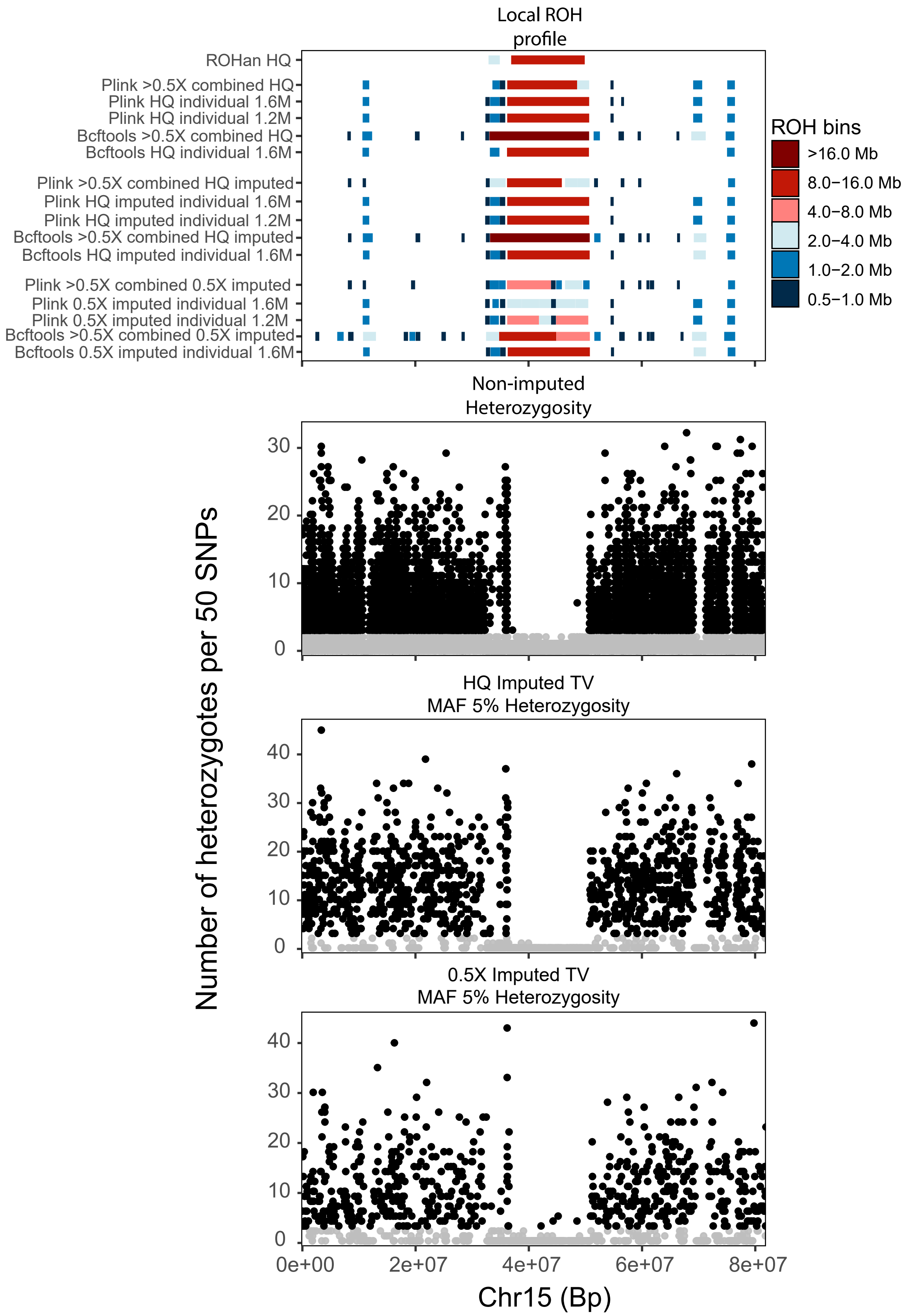


**Figure S35: Local ROH bin length and heterozygous profile on chromosome 15 for imputed 0.5X and high coverage (14.4X) , and non-imputed high coverage (14.4X) Semnan3 genomes across different datasets.** Bars represent ROH bin lengths, represented in color. Dots represent the number of heterozygotes in a 50 SNP window, black indicates >2 heterozygotes and grey indicates ≤2 heterozygotes.

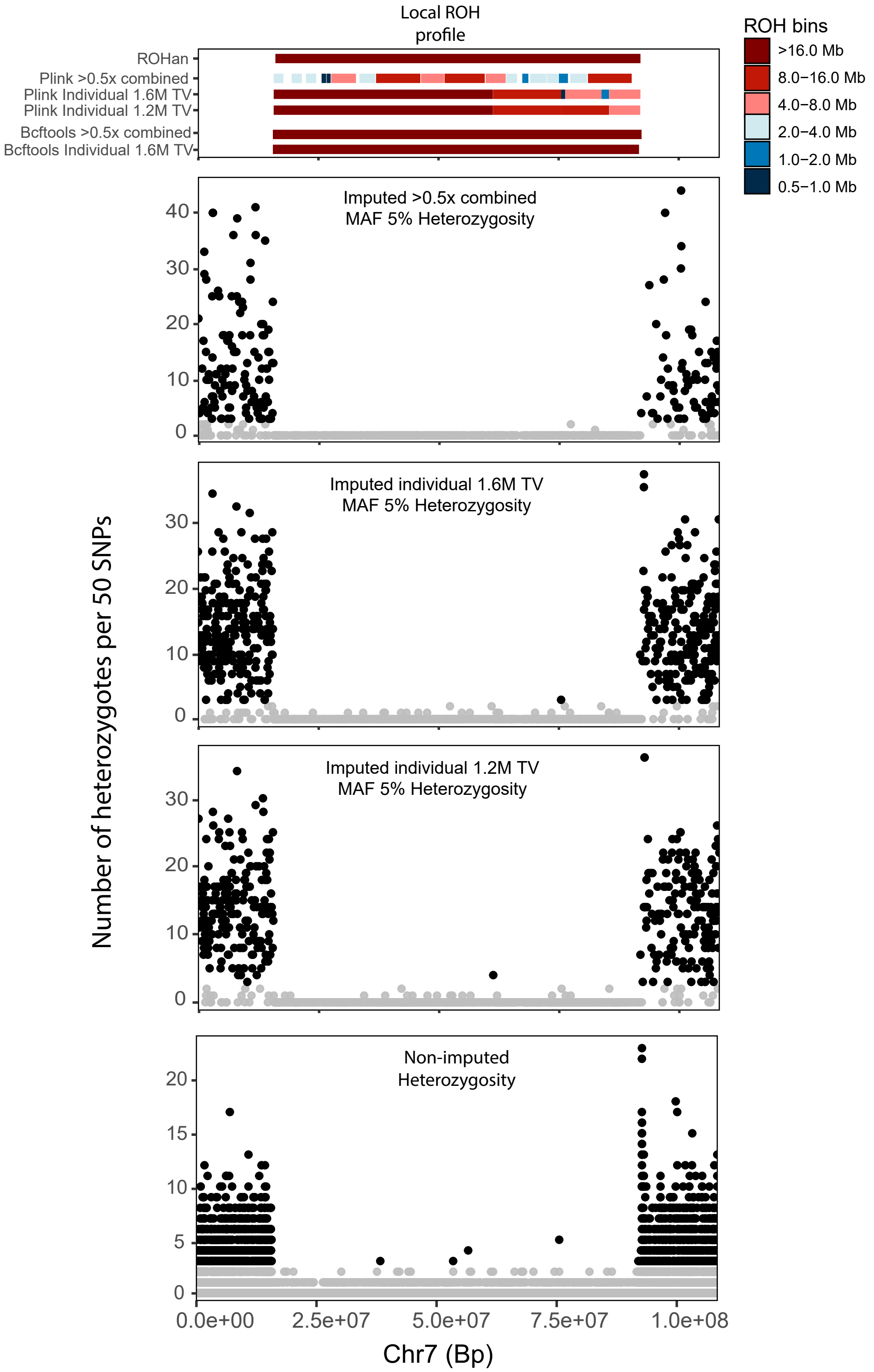


**Figure S36: Local ROH bin length and heterozygous profile on chromosome 7 for imputed and non-imputed Potterne1 (3.6X) genomes across different datasets.** Bars represent ROH bin lengths, represented in color. Dots represent the number of heterozygotes in a 50 SNP window, black indicates >2 heterozygotes and grey indicates ≤2 heterozygotes.

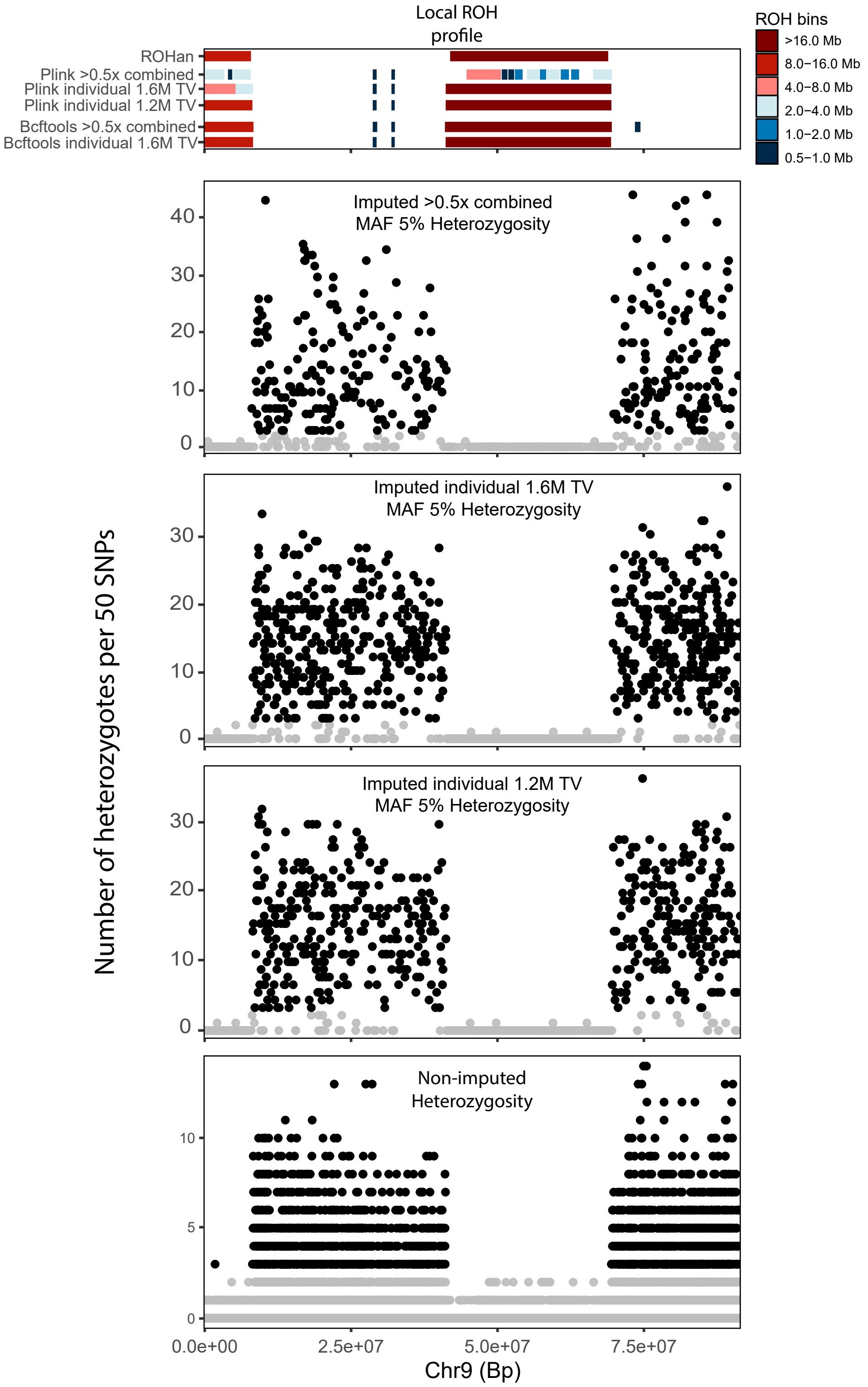


**Figure S37: Local ROH bin length and heterozygous profile on chromosome 9 for imputed and non-imputed Blagotin16 (3.5X) genomes across different datasets.** Bars represent ROH bin lengths, represented in color. Dots represent the number of heterozygotes in a 50 SNP window, black indicates >2 heterozygotes and grey indicates ≤2 heterozygotes.

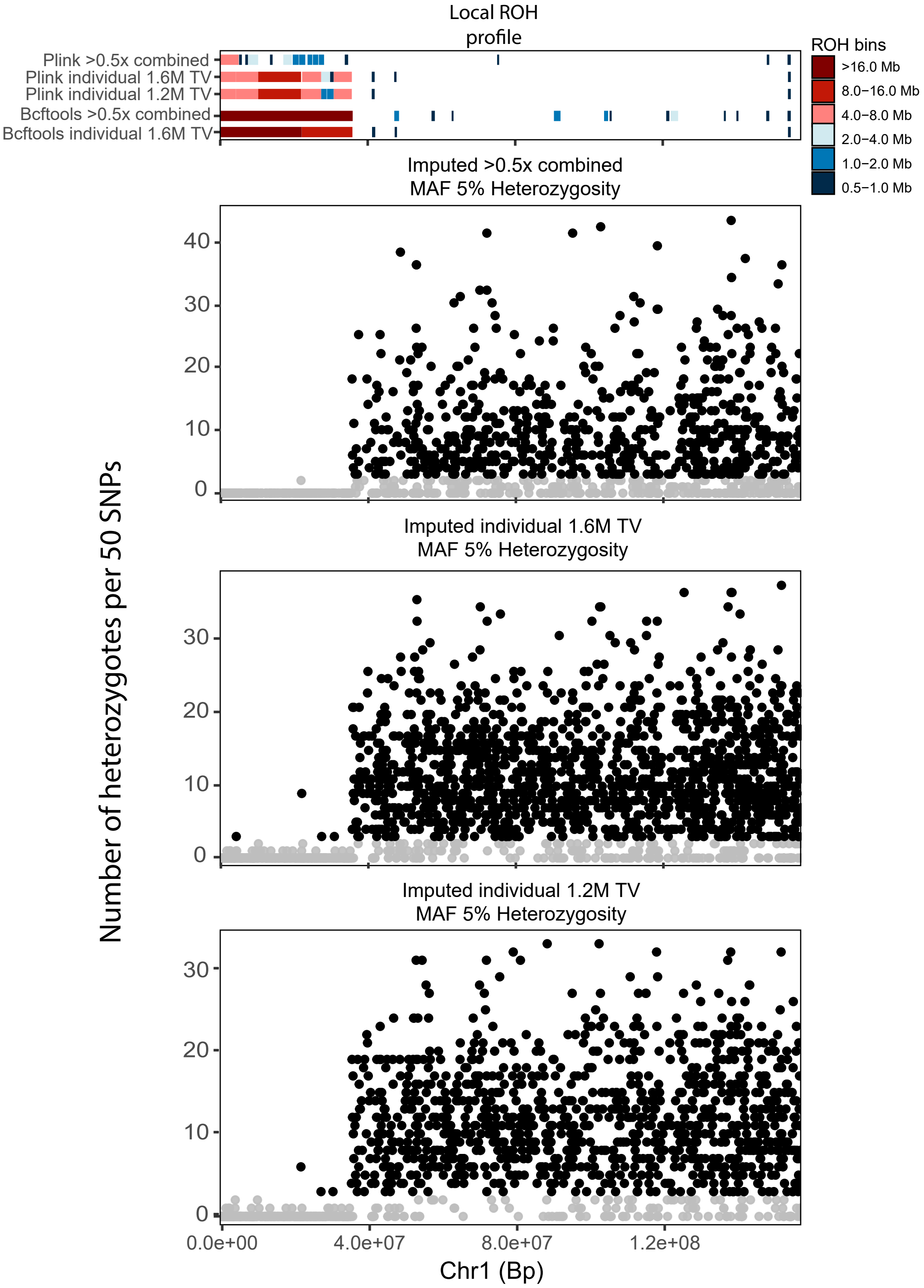


**Figure S38: Local ROH bin length and heterozygous profile on chromosome 1 for the imputed Bulak1 (0.9X) genome across different datasets.** Bars represent ROH bin lengths, represented in color. Dots represent the number of heterozygotes in a 50 SNP window, black indicates >2 heterozygotes and grey indicates ≤2 heterozygotes.

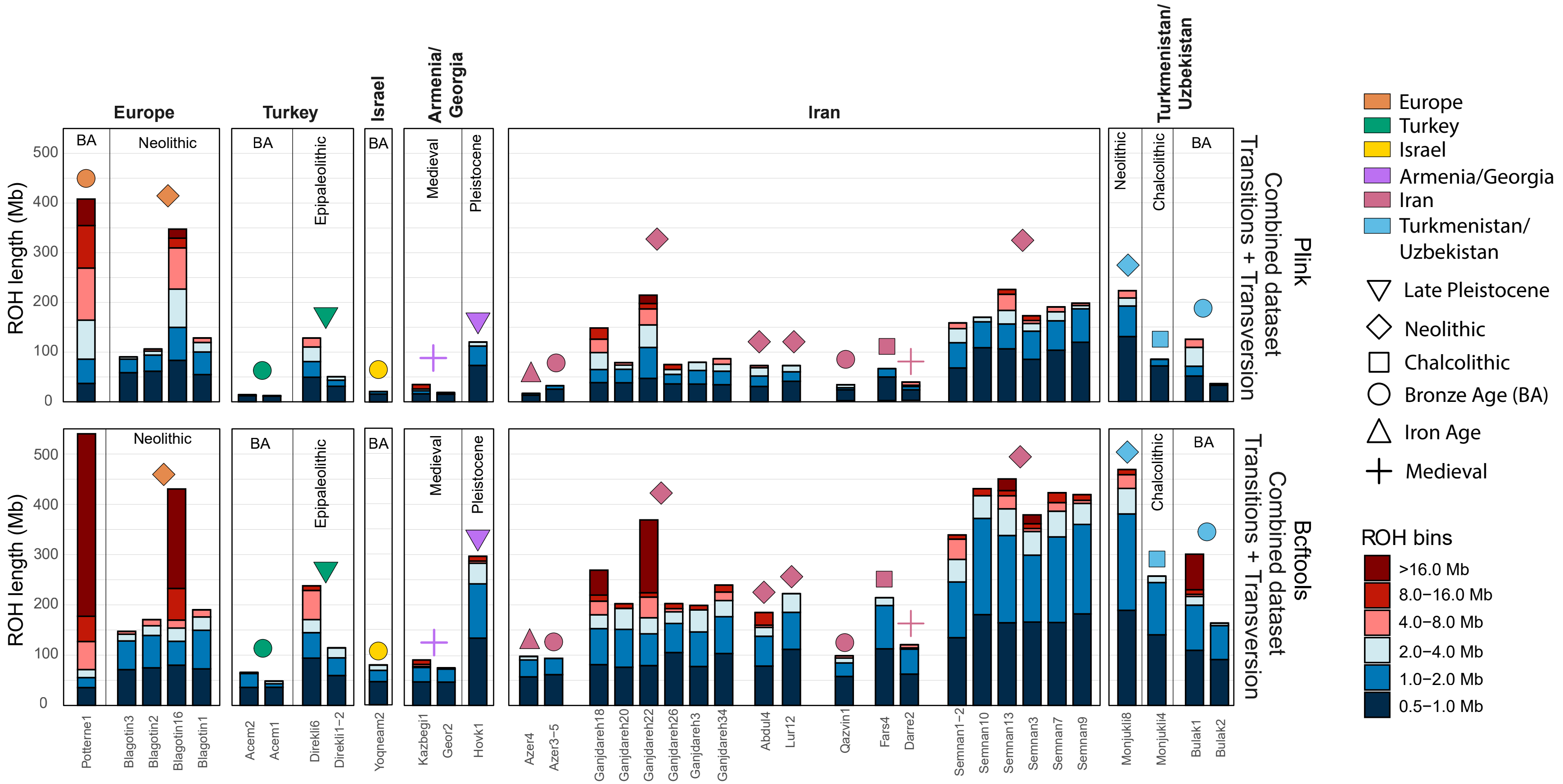


**Figure S39: ROH length bin profiles for ≥0.5X imputed ancient goats for Plink and Bcftools.** ROH was calculated combined and were conducted using both Plink and Bcftools. BA = Bronze Age.

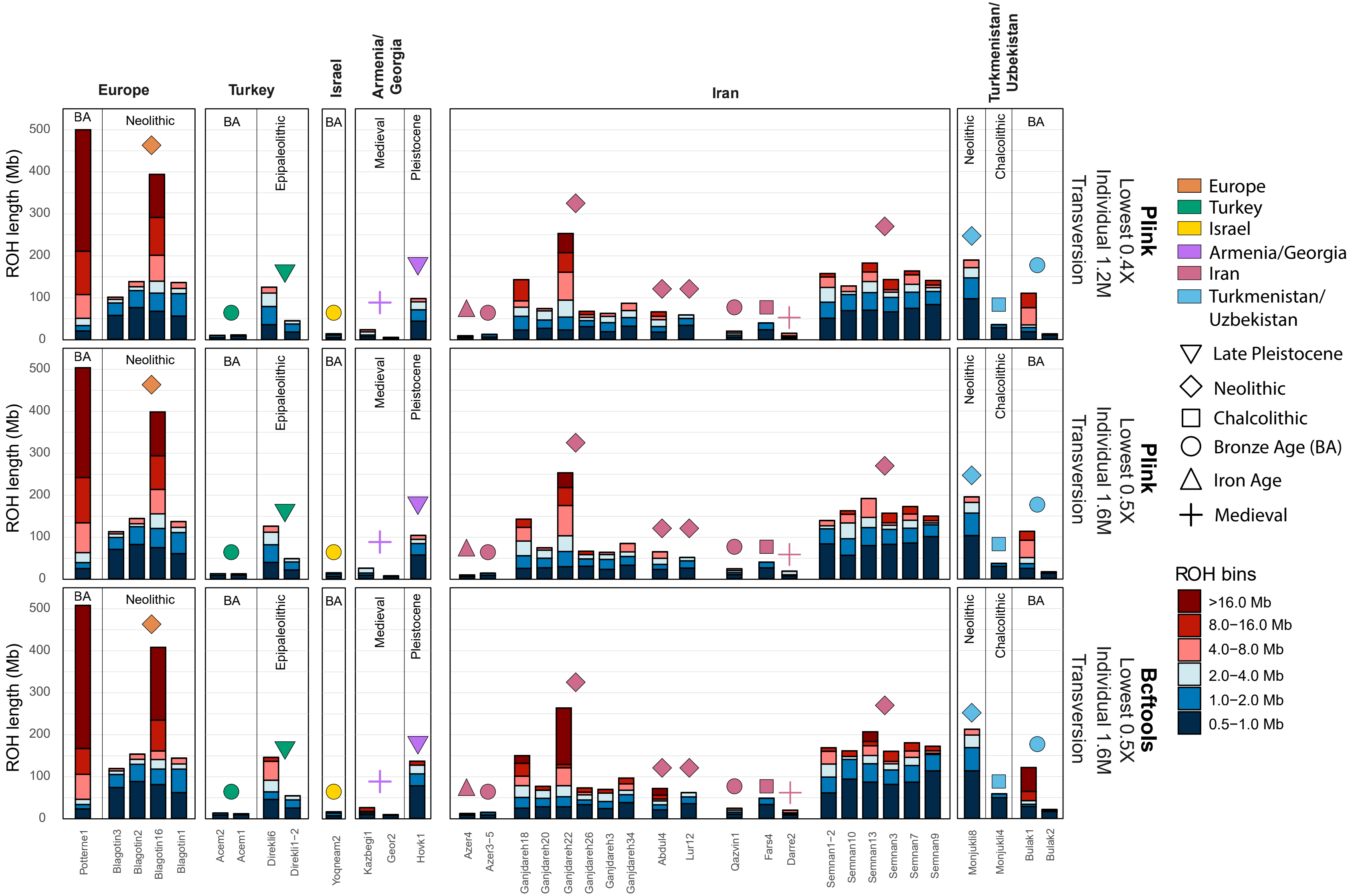

**Figure S40: ROH length bin profiles for ≥0.5X imputed ancient goats across different datasets.** ROH was calculated individually for each ancient sample using transversions only. Downsampling was performed at 1.2 million and 1.6 million sites, corresponding to the lowest number of imputed sites in the 0.4X and 0.5X imputed test samples, respectively. ROH calculations were conducted using both Plink and Bcftools on the 1.6 million-site dataset. BA = Bronze Age.

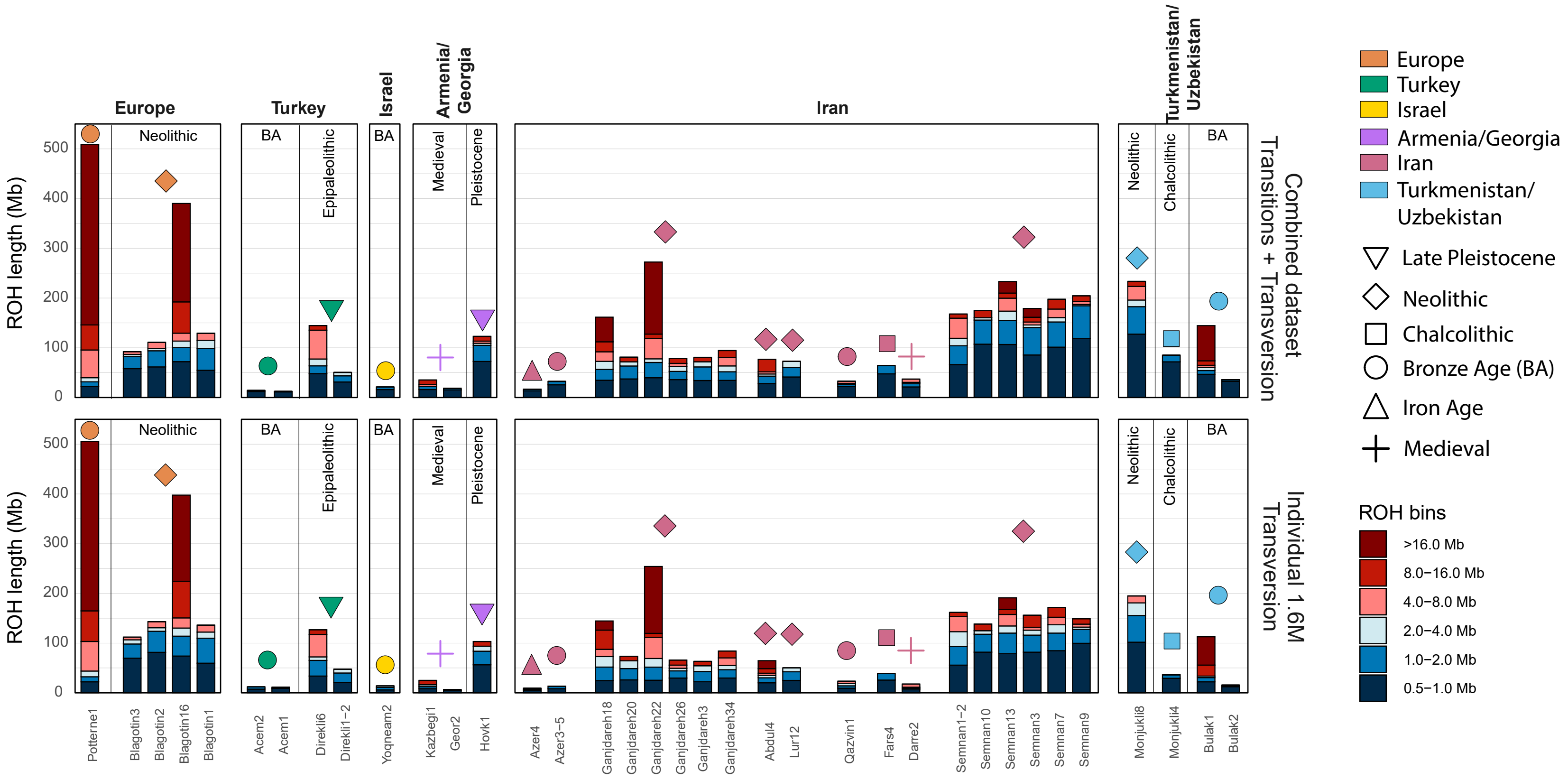


**Figure S41: ROH length bin profiles for ≥0.5X imputed ancient goats, stratified by dataset used.** The upper plot shows ROH calculated using the same transition and transversion genotypes across all samples, with no missingness (897K sites). The lower plot shows ROH calculated with each ancient sample individually, using transversions-only and downsampling to 1.6 million sites, corresponding to the lowest number of imputed sites in the 0.5X imputed test samples. ROH profiles result from combining plink ROH and long ROH (>4Mb) from bcftools.
